# The Receptor Kinase MEE39/ATHE Mediates Cell Wall Integrity Surveillance During Root Vascular Pathogen Infection

**DOI:** 10.64898/2026.03.05.708959

**Authors:** Juan Carlos Montesinos, Marina Martín-Dacal, Hsin-Yao Huang, Gloria Sancho-Andrés, Fortesa Rama, Laura Carrillo, Anurag Kashyap, Álvaro Jiménez-Jiménez, Francisco M. Gámez-Arjona, Caroline Broyart, Huanjie Yang, Nuria S. Coll, Julia Santiago, Cyril Zipfel, Clara Sánchez-Rodríguez

## Abstract

Plant cell wall (CW) integrity signaling enables early detection of microbial invasion, yet the receptors involved and their spatial and temporal dynamics during infection remain largely unknown. We identify ATHENA (ATHE)/MEE39, a previously uncharacterized malectin-like leucin-rich repeat receptor kinase (Mal-LRR-RK) that contributes to defense against the root vascular pathogen *Fusarium oxysporum* (Fo), particularly in outer root layers where colonization begins. ATHE abundance, localization, and endocytic trafficking are rapidly remodeled during infection, and loss of ATHE compromises basal immunity and early pathogen-induced transcriptional reprogramming. ATHE responds to altered cellulose synthesis, cellulose-derived oligosaccharides, mechanically induced CW perturbations, and the fungal secreted peptide Fo-RALF. In most of these contexts, ATHE acts together with the LRR-RK MIK2, forming a pathogen-strengthened RK complex that fine-tunes root responses to Fo. This represents the first example of a receptor complex visualized subcellularly *in vivo* during a plant-microbe interaction. Although *Brassicaceae*-specific, heterologous expression of *ATHE* enhanced tomato resistance to Fo, highlighting its functional relevance across plant lineages and its potential use for crop engineering. Our work reveals a previously unrecognized strategy by which plants decode microbial threats through dynamic CW-integrity surveillance.

## INTRODUCTION

Cells continuously monitor their extracellular matrix to coordinate development with environmental and internal cues. In plants, this process is particularly critical because the cell wall (CW) constitutes both a structural barrier and a dynamic signaling interface. During plant-microbe interactions, the CW acts as a source of nutrients for the intruder and a primary barrier that must be modified for microbial growth. Conversely, plants detect CW damage, shaping the infection outcome ^1,2^. Most plant-associated microbes inhabit the apoplast, which is filled by primary CWs, including root-infecting vascular pathogens such as the fungus *Fusarium oxysporum* (Fo) *sps.* ^3,4^. Fo enters the root through epidermal cells and expands through apoplastic spaces to reach the vasculature, causing xylem occlusion and wilting symptoms that severely impact crop yields ^3,5–9^. During infection, Fo secretes CW-modifying enzymes, including cellulases, xylanases, esterases, and polygalacturonases, to promote apoplastic invasion ^10–13^. In the *Arabidopsis thaliana* (hereafter Arabidopsis)-Fo5176 model, the fungus disrupts host CW integrity by removing cellulose synthase complexes from the plasma membrane (PM), disassembling cortical microtubules, and downregulating cellulose biosynthesis genes, collectively reducing cellulose deposition and shaping the outcome of Fo-plant interactions ^14^. CW degradation generates damage associated molecular patterns (DAMPs) that activate immune responses ^8,15,16^ and, together with mechanical stress caused by hyphal pressure, disrupts cell-cell adhesion ^17^. Despite this complex mixture of apoplastic cues, relatively little is known about the receptors that initiate these signaling pathways.

Hence, it is expected that a set of PM-localized pattern-recognition receptors (PRRs) detect CW-integrity (CWI) alterations caused by vascular pathogens and activate pattern triggered immunity. Many PRRs contain an extracellular leucine-rich repeat (LRR) domain and can also contain malectin, malectin-like, lectin, EGF-like, or LysM domains that determine ligand specificity ^18,19^. Multiple PRRs are receptor kinases (RKs) ^20^, yet few have been functionally linked to vascular-pathogen responses or CWI signaling. Among them, THESEUS1 (THE1), a member of the *Catharantus roseus* RLK1-like (CrRLK1L) family, participates in responses to cellulose synthesis defects ^21^, although its contribution to resistance against Fo5176 remains debated ^22^. The LRR-RK MALE DISCOVERER 1 - INTERACTING RECEPTOR LIKE KINASE 2 (MIK2) exhibits both overlapping and distinct functions with THE1 in CWI sensing and Fo5176 resistance ^22^. MIK2 is also the receptor for Serine-rich endogenous peptide (SCOOP) family members, including SCOOP12 and SCOOP18, which regulates development and immunity ^2,23–26^. Another group of phytocytokines, the rapid alkalinization factors (RALFs), also play central roles in immunity and CWI perception ^27,28^. Several pathogens, including Fo, secrete RALF-like mimics that hijack the FERONIA (FER) receptor to suppress host immunity ^29–31^. Various RALFs exemplify CW-peptide integration by binding de-methylesterified pectins via FER-based receptor complex, illustrating the nature of the CrRLK1L-RALF network as a flexible receptor-ligand system rather than simple one-to-one interactions ^32,33^. FER, like THE1, belongs to the CrRLK1L family and integrates CW-derived signals with developmental and immune pathways ^34,35^. Recent structural and mechanistic analyses highlight FER’s pleiotropy and its role in sensing CW tension ^36^. Additionally, the RK WALL ASSOCIATED KINASE (WAK)-LIKE 22/ RESISTANCE TO FUSARIUM OXYSPORUM 1 (WAKL22/RFO1) mediates responses to altered pectin methylesterification, contributes to resistance against root fungal pathogens like Fo5176, and modulates plant growth ^37–39^. These few examples evidence the need to identify additional PRRs involved in sensing CW perturbations during vascular pathogen infection.

RK-mediated signaling depends not only on ligand perception but also on the cell’s ability to dynamically adjust receptor abundance, localization, and activity. Because PRRs must be rapidly activated, attenuated, or recycled during infection, clathrin-mediated endocytosis (CME) should function as a key mechanism linking CW perception to downstream signaling. Through CME, receptors are internalized from the PM and directed to the multivesicular body/prevacuolar compartment (MVB/PVC) pathway for vacuolar trafficking, turnover, and signal attenuation, thereby maintaining immune homeostasis and PM organization ^40–42^. In plants, CME requires the TPLATE complex (TPC), an essential adaptor module that initiates endocytic vesicle formation ^43,44^. This regulatory system has been well documented for several RKs, including SOMATIC EMBRYOGENESIS RECEPTOR-LIKE KINASE 1 (SERK1), BRASSINOSTEROID INSENSITIVE 1 (BRI1), FLAGELLIN-SENSING 2 (FLS2), LysM-CONTAINING RECEPTOR-LIKE KINASE 5 (LYK5), PEP RECEPTOR 1 (PEPR1), and EF-TU RECEPTOR (EFR) ^45–51^. CME also maintains cellular homeostasis by recycling membrane components via the trans-Golgi network and endosomal system ^52,53^, and can mediate uptake of pathogen-derived extracellular vesicles and may facilitate entry of filamentous-pathogen effectors ^54,55^, highlighting its broader role in host-pathogen communication.

Therefore, we hypothesize that novel PRRs involved in sensing Fo-induced CW alterations can be identified among receptors undergoing CME. Using the Arabidopsis-Fo5176 model, we aim to elucidate how early CW perturbations are perceived and how these initial signaling events shape plant responses to vascular pathogen attack.

## RESULTS

### Clathrin-Mediated Endocytosis (CME) early adaptor complexes are required for plant defense against Fo5176

To assess the contribution of CME to the Arabidopsis-Fo5176 interaction, we performed fungal plate-infection assays ^56^ using the *ap2m-1* and *twd40-2-3* mutants, impaired in the early adaptor complexes AP-2 and TPLATE (TPC), respectively ^57–59^. Only *ap2m-1* showed significant reduced sensitivity to the previously reported Fo-induced root growth inhibition (Fig. 1A; ^14^). Both CME adaptor mutants exhibited increased fungal penetration into the vasculature compared to wild-type (WT; Fig. 1B). This increased susceptibility was confirmed in soil assays where Fo-induced wilting symptoms were tracked (Fig. 1C). Our findings show that impairment of CME early adaptors disrupts root defense mechanisms during Fo5176 colonization.

**Figure 1.**
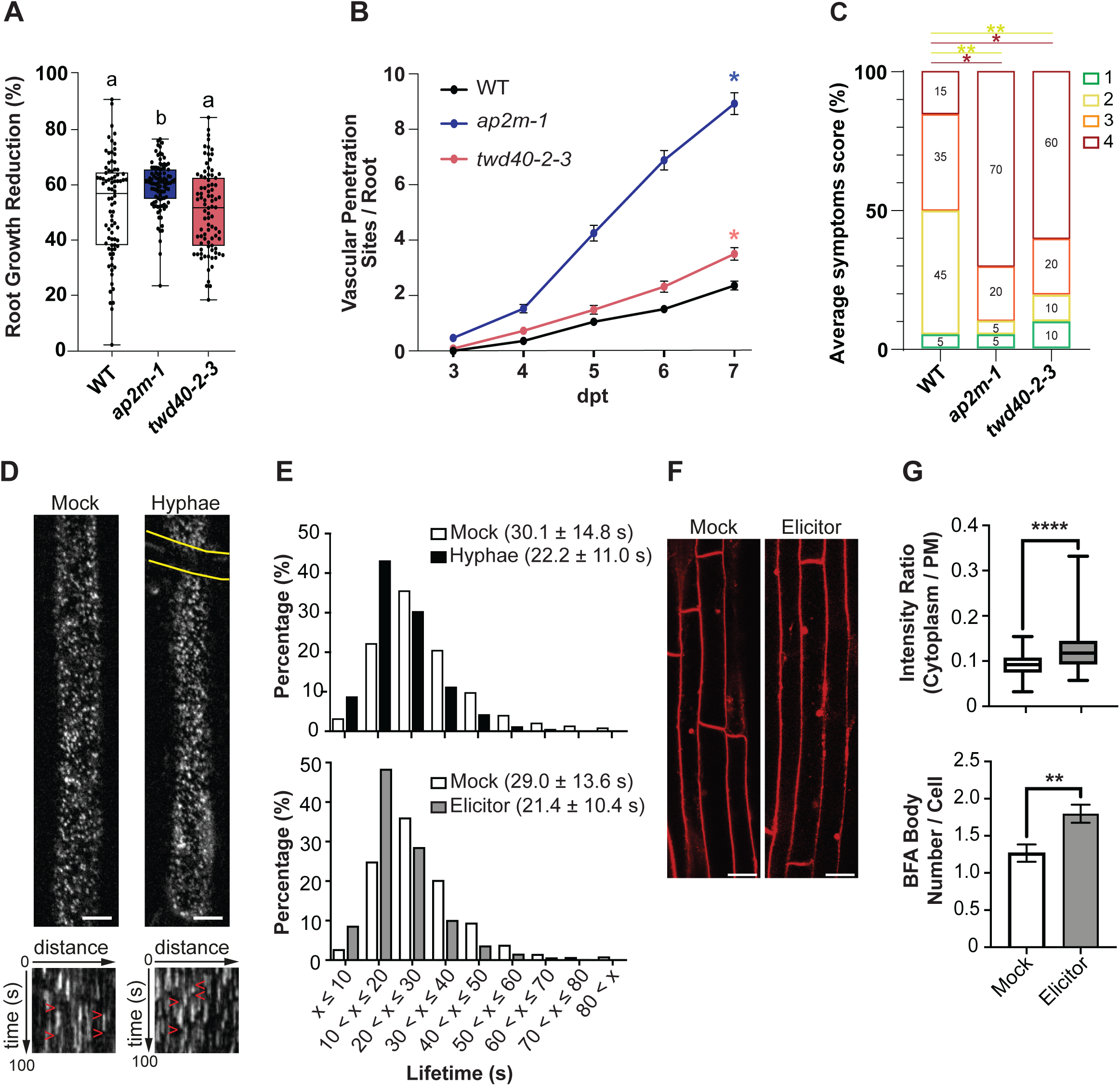
The CME early adaptor proteins are required for plant defense against Fo5176. **A.** Root growth reduction of Fo5176 *pSIX1:GFP*-infected wild-type (WT; Col-0), *ap2m-1,* and *twd40-2-3* seedlings relative to Mock-treated ones, at 6 days post treatment (dpt). The box plots are shown: centerlines show the medians; box limits indicate the 25th and 75th percentiles; whiskers extend to the minimum and maximum. *N* ≥ 82 seedling per genotype from 3 independent replicates (*N* of roots ≥ 20/experiment). Repeated measures (RM) one-way ANOVA with Dunn’s multiple comparisons test. Alphabet letters represent groups of significant differences with P ≤ 0.05. **B.** Cumulative number of Fo5176 *pSIX1:GFP* vascular penetrations per root in WT, *ap2m-1,* or *twd40-2-3* from 3 to 7 dpt. Av ± SEM; *N* ≥ 96 roots per genotype from 3 experiments (*N* of roots ≥ 26/experiment). RM two-way analysis of variance (ANOVA) with Tukey’s multiple comparisons test, *\**P ≤ 0.05, with respect to WT at 7 dpt. **C.** Scoring of Fo5176-induced disease symptoms in soil-infected WT, *ap2m-1*, and *twd40-2-3* plants at 14 dpt. Phenotypes: 1 (asymptomatic plants), 2 (plants with ≤ 50% of yellow or dry leaves), 3 (plants with > 50% of yellow or dry leaves), or 4 (dead plants). Data represent average in percentage (%); *N* = 20 plants per genotype from 1 experiment out of 3 with similar results. RM Chi^2^ analysis, *P ≤ 0.05, **P ≤ 0.005. The colors of the asterisks correspond to the colors assigned to the different symptom phenotypes. **D.** Representative spinning disc images (upper panels) and kymographs from the corresponding movies (n = 100 frames, t = 100 s; lower panels) of a root epidermal cell at the maturation zone, expressing TML-GFP treated for 5 min with Mock or Fo5176 young hyphae (yellow lines). Images and kymographs show the discrete and dynamic TML-GFP foci. Red arrowheads in the kymographs mark the start and the end of dwell time of foci at the plasma membrane. Scale bars = 5 μm. **E.** Lifetime distribution of TML-GFP foci at the plasma membrane of root epidermal cells upon live fungus (upper panel), or Fo5176 elicitor mix (lower panel), and the corresponding Mock treatments (1/2 MS or water, respectively). Average lifetime of the TML-GFP foci indicated in the bracket (Av ± SD), *N*≥ 1500 foci pretreatment: N ≥ 150 foci/cell; ≥ 8 cells/ treatment from 2-3 experiments (2-3 cells/seedling and 2-3 seedlings/experiment were imaged). **F.** Representative images of root epidermal cells at the maturation zone stained with FM4-64 (red) and treated with BFA and either Mock or Fo5176 elicitor mix for 10 min at room temperature. Scale bars = 20 μm. **G.** Quantification of endocytosis activity in cells as shown in (F). Upper panel: cytosolic signal intensity is normalized to the PM one to indicate the amount of internalized FM4-64. Lower panel: BFA body numbers per cell were counted. Box plots as described in (A). N of cells ≥ 61 cells from 3 experiments (3-5 cells/seedling and N of seedlings ≥ 3 /experiment were imaged). Data represent the Av ± SE. RM t-test, **P ≤ 0.01, ****P ≤ 0.0001.

To monitor the modulation of CME adaptors during Arabidopsis-Fo5176 interaction, we examined the dynamics of the GFP-tagged TPC subunit, TML ^43^, at the PM. We mimicked early contact events of pathogen infection (as 1 day post treatment (dpt) shown in ^17^), by exposing the root to Fo5176 young hyphae. In epidermal cells of the differentiated root zone, the TML-GFP signal appeared as discrete foci at the PM, which dynamically appear and disappear without lateral movement, as previously described in etiolated hypocotyls (^43^, Fig 1D). The PM dwell time (lifetime) of TML-GFP was reduced in cells directly contacted by a live Fo5176 hypha, relative to adjacent non-contacted cells (Fig. 1E, upper panel), as response to chemical and/or mechanical fungal signals. A similar reduction in dwell time was triggered by the Fo5176 elicitor mix (Fig. 1E, bottom panel), which has been shown to induce rapid cellulose synthesis alterations ^14^, indicating that the plant CME adaptor can respond to Fo5176 chemical signals. The reduced TML-GFP lifetimes may indicate elevated endocytic activity during microbe-associated molecular pattern (MAMP) perception, consistent with the role of TPC in CME initiation. To test this, we quantified the uptake of the membrane fluorescent marker FM4-64 in the presence of Brefeldin A (BFA), which generates trans-Golgi network (TGN)-derived BFA bodies (Fig. 1F and G). Fo5176 elicitor mix caused significant increases in both FM4-64 cytoplasm/PM intensity ratios and BFA body numbers relative to mock. These results show that Fo5176-molecules enhance CME, supporting the link between decreased TML lifetimes and CME activation.

### Identification of the uncharacterized Mal-LRR-RLK MEE39/ATHE involved in plant response to Fo5176

To identify PRRs internalized via CME and potentially involved in plant response to Fo-induced CW changes, we performed an organelle immunoprecipitation (IP) assay using seedlings expressing either the MVB/PVC marker ARA7-YFP ^60^ or free-GFP as negative control ^43^ (Fig S1A). We immuno-enroched MVB/PVCs from roots at 3 dpt with mock or Fo5176 spores under hydroponic conditions, a time point at which the hyphae have not yet reached the vasculature ^17^. Using LC-MS/MS analysis followed by MS2 spectra detection, we identified 79 and 46 proteins in mock and Fo5176-treated samples in ARA7-YFP and free-GFP IP profiles, respectively (Table S1). After discarding entities appearing in both free-GFP and ARA7-YFP samples, we had a list of 77 candidates (Fig 2A). Among them, there was a member of the LRR-RK-I family (AT3G46330), composed of an extracellular malectin-like domain (MLD) and three LRRs, followed by a transmembrane domain and a cytosolic kinase domain (Fig. S1B). MLD is named upon sequence homology to malectin that binds carbohydrates in animals ^61^. Indeed, recent studies show that the LRR-Malectin ectodomain can bind cellulose-derived oligomers derived from plant CW degradation by pathogens ^62,63^. Thus, we focused on this receptor for further characterization as a potential sensor of Fo-derived CW alterations. Originally, AT3G46330 was annotated as MEE39 (MATERNAL EFFECT EMBRYO ARREST 39) ^64^, which does not seem to correspond with its biological role, since the mutant is phenotypically similar to the WT under control conditions (Fig. 4F). Therefore, we also named it ATHENA (*<u>A</u>rabidopsis <u>th</u>aliana* <u>en</u>hanced resistance to v<u>a</u>scular pathogens, ATHE), as a potential receptor mediating plant defense against vascular pathogens, a name that reflects its biological function reported in this study.

**Figure 2.**
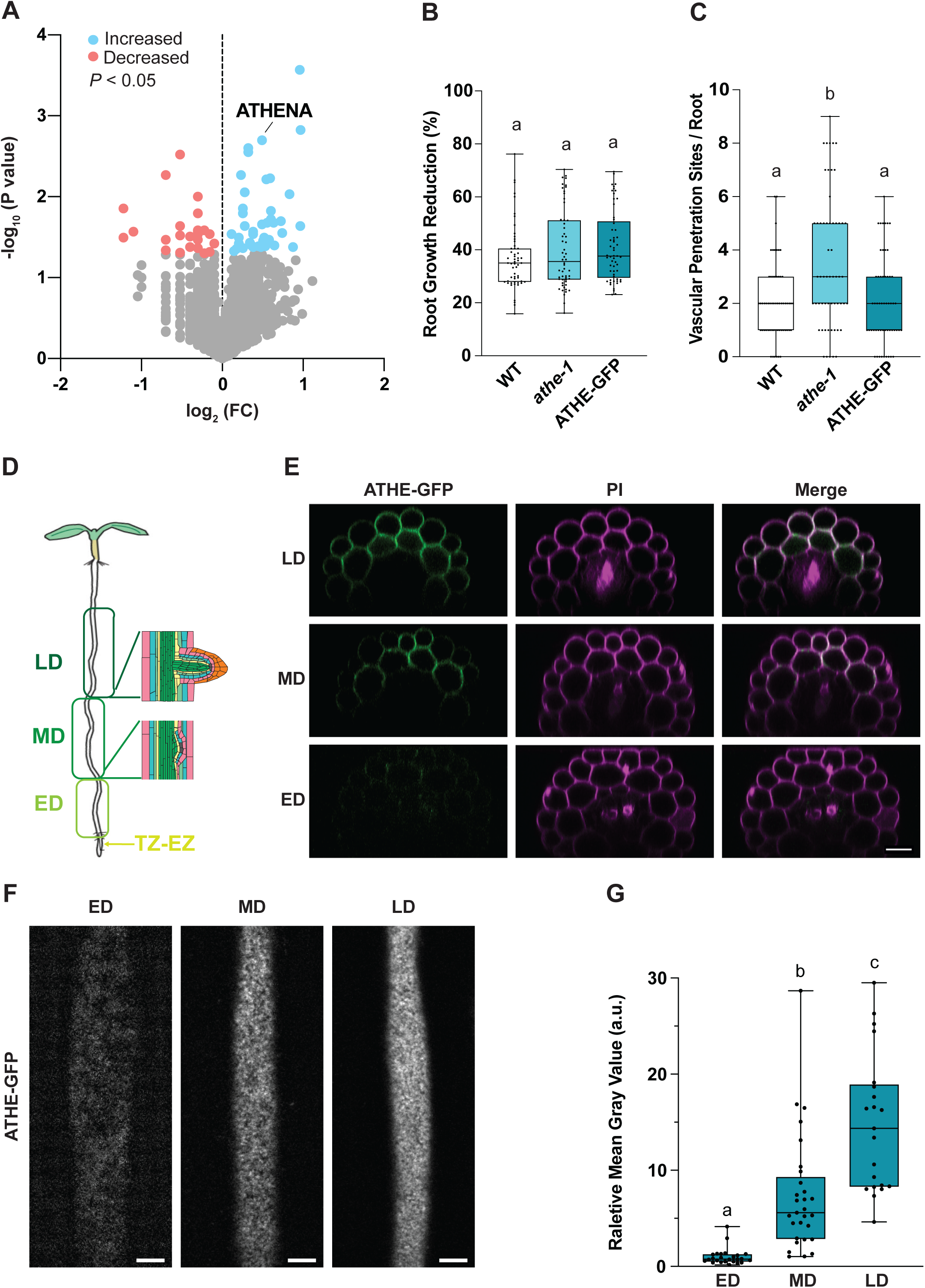
ATHENA is a PM-localized LRR-RK receptor protein contributing to plant defense against Fo infection. **A.** Volcano plot of differences in the abundance of proteins identified in the ARA7-YFP IP fraction upon Fo5176 hydroponic infection compared to Mock conditions. *N* = 4 independent replicates. **B.** Root growth reduction of Fo5176 *pSIX1:GFP*-infected wild type (WT; Col-0), *athe-1,* and ATHE-GFP seedlings relative to Mock-treated ones, after 6 days post treatment (dpt).The box plots are shown: centerlines show the medians; box limits indicate the 25th and 75th percentiles; whiskers extend to the minimum and maximum. *N* ≥ 55 roots from 3 experiments (*N* of roots ≥ 15/experiment). Repeated Measures (RM) one-way ANOVA with Kruskal-Wallis multiple comparison test. Alphabet letters represent groups of significant differences with P ≤ 0.05 **C.** Cumulative number of Fo5176 *pSIX1:GFP* vascular penetrations per root in WT, *athe-1*, and ATHE-GFP at 7 dpt as described in (B). Box plots as described in (B), *N* ≥ 50 roots from 3 experiments (*N* of roots ≥ 15/experiment). RM one-way ANOVA with Kruskal-Wallis multiple comparison test. Alphabet letters represent groups of significant differences with P ≤ 0.05. **D.** Root zones defined in this study: TZ-EZ, transition-elongation zone; ED, early differentiation zone; MD, middle differentiation zone; LD, late differentiation zone. **E.** Representative confocal microscope optical cross sections of ATHE-GFP (green) in 8-day-old roots stained with Propidium Iodide (PI; magenta) at different root differentiated zones, as indicated in (D). Scale bar = 20 μm. *N* = 15 roots observed with similar results. **F.** Representative spinning disk confocal images of ATHE-GFP in root epidermal cells at the root differentiated zones indicated in (D). Scale bars = 5 μm. **G.** Amount of ATHE-GFP at the plasma membrane of epidermal cells as shown in (F), quantified as mean gray value (arbitrary units, a.u.) normalized to the average in the ED. Box plots as described in (B), *N* ≥ 21 cells from 3 experiments (2-4 cells/root and ≥ 3 roots/experiments). RM one-way ANOVA with Kruskal-Wallis multiple comparison test. Alphabet letters indicate significant differences with P ≤ 0.05.

### ATHENA participates in plant defense against vascular pathogens

The expected role of ATHE in plant-Fo interaction was tested using a mutant containing a single T-DNA insertion, generating a knock-out allele of *ATHE*, *athe-1* (Fig. S1B). Plate infection assays using fluorescently labeled Fo5176 pSIX1:GFP ^56^ showed no difference in the fungus-induced root growth inhibition in *athe-1* compared to WT plants (Fig. 2B), but the mutant displayed a higher frequency of fungal vascular penetrations than the WT (Fig. 2C). The complemented line ATHE-GFP (*athe-1* pATHE::ATHE-GFP) was not distinguishable from WT plants regarding fungal-induced root growth inhibition and vascular penetrations (Fig. 2B and C). To correlate ATHE’s role in reducing Fo5176 colonization of the root vasculature with its capacity to proliferate here and cause wilting symptoms, we performed soil infection assays. According to the phenotypes observed in plates, *athe-1* mutant displayed more severe wilting symptoms and dead plants than WT, whereas ATHE-GFP was very similar to WT regarding Fo5176 disease symptoms (Fig. S1 C and D). In addition, another soil-borne vascular pathogen, the bacterium *Ralstonia solanacearum* (Rs), also caused more severe wilting symptoms and a higher disease score in *athe-1* plants compared to WT (Fig. S1E and F). Altogether, these results suggest that ATHE is involved in plant defense against two root vascular pathogens from different superkingdoms.

### ATHENA localizes at the PM of root cells with gradual distribution along the differentiated area of the root axis

According to its predicted structure, ATHE should localize at the PM, which we confirmed using confocal microscopy. The subcellular localization of ATHE was examined in different regions of the root axis: early differentiated zone (ED), middle-differentiated zone (MD), and late differentiated zone (LD), which are marked by the emergence of root hairs, the presence of the first lateral root primordium initiation, and the first emerged lateral root, respectively (Fig. 2D). ATHE-GFP was visible at the PM of epidermal and cortical cells (Fig. 2E) showing an asymmetrical distribution along the root axis. Thus, ATHE-GFP signal was detectable only in differentiated cells, increasing progressively from ED to LD, but was nearly absent in the ED (Fig. 2E). The PM localization of ATHE-GFP was confirmed by a plasmolysis assay using mannitol treatment and comparison with the PM marker Lti6B-GFP (^65^; Fig. S1G). A detailed subcellular localization analysis of ATHE-GFP in epidermal PM revealed a punctate pattern typical of PM receptors and confirmed its asymmetrical distribution along the root axis (Fig. F and G).

Our results confirmed the predicted PM localization of ATHE, which displayed a punctate, foci-like distribution. We also found that ATHE is predominantly enriched in the outer root cell layers of the MD and LD zones, where Fo5176 preferentially penetrates the epidermis, and that its accumulation gradually increases along the root axis.

### ATHENA protein localization, abundance, and gene expression are altered by Fo5176 infection

Consistent with the proposed internalization of ATHE from the PM as part of its potential role in the Fo5176-induced response, we observed a pronounced depletion of ATHE-GFP from the PM in most cells at 1dpt with Fo5176 young hyphae (Fig. 3A). This timing is biologically comparable to 2 dpt with Fo5176 spores in plate-based infections, when the hyphae have already colonized the apoplast of the epidermal root layer ^17^. In contrast, ATHE-GFP accumulation in the ED increased in response to the fungus, whereas under mock conditions it was barely detectable (Fig. 3A). The notable decrease in ATHE-GFP signal at the PM of MD cells upon Fo5176 treatment (Fig. 3A) was confirmed at the subcellular level in epidermal MD cells (Fig. 3B and C). Short-term interaction with Fo5176 young hyphae (15-30 min; mimicking early contact events of pathogen infection) did not alter the PM localization of ATHE-GFP or the PM-marker NSPN12-YFP (^60^; Fig. S2 A and B), although this treatment was previously shown to reduce the abundance of cellulose synthase complexes at the PM ^14^. This time-dependent relocalization of ATHE is consistent with its potential internalization triggered by the direct perception of, or a co-receptor-mediated response to Fo-derived signal.

**Figure 3.**
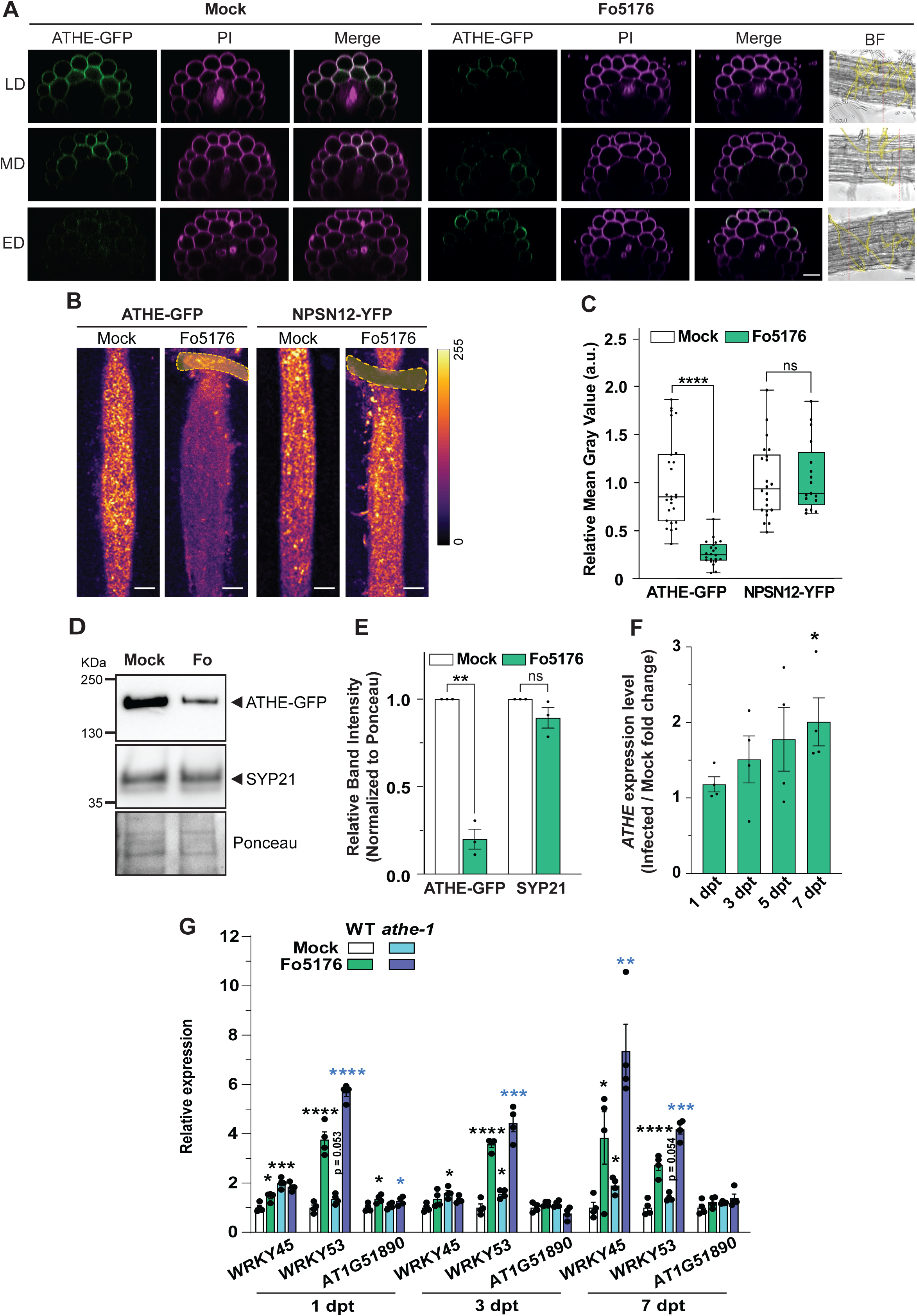
ATHE-GFP protein level and gene expression are altered by Fo5176 infection. **A.** Representative confocal microscope optical cross section of ATHE-GFP (green) in 8-day-old roots stained with Propidium Iodide (PI; magenta) at the root differentiated zones indicated in Fig 2D, after Mock (left) and 1-day of exposure to Fo5176 young hyphae. Bright-field (BF) images show the positions used to generate the orthogonal views (dashed red lines) and the hyphae (yellow). Scale bars = 20 μm. *N* = 30 roots/treatment observed with similar results **B.** Representative spinning disk confocal images of ATHE-GFP or NPSN12-YFP in 8-day-old root epidermal cells in the MD after 1-day of exposure to Fo5176 young hyphae (yellow). The scale represents the fluorescence signal intensity of the tagged proteins. Scale bars = 5 μm. **C.** Amount of ATHE-GFP and NPSN12-YFP at the plasma membranes of cells as described in (B), quantified as mean gray value (arbitrary units, a.u.) of fungal-exposed samples normalized to their corresponding Mock. Box plots: centerlines show the medians; box limits indicate the 25th and 75th percentiles; whiskers extend to the minimum and maximum. *N* ≥ 17 from 3 experiments (2-4 cells/root and ≥ 3 roots/experiments). Repeated measures (RM) *t*-test. ****P ≤ 0.0001, ns: not significant. **D.** Representative Western blot of total ATHE-GFP and SYP21 proteins in 8-day-old ATHE-GFP roots at 1 day post treatment (dpt) with Mock or Fo5176 young hyphae. Ponceau S staining provides loading control. **E.** ATHE-GFP and SYP21 signal quantification from immunoblots as in (D). For each protein, the intensity of the band was first normalized to its corresponding Ponceau signal. Then, the value for the fungal-exposed sample was normalized to that of the Mock. Av ± SEM, *N* = 3 independent experiments. RM *t*-test. ** *P* ≤ 0.01, ns: not significant. **F.** *ATHE* expression relative to *AtGAPDH* in 8-day-old WT roots at different dpt to Fo5176-containing plates, normalized to Mock-treated roots. Av ± SEM, *N* = 4 independent replicates (≥ 20 roots/experiment). RM *t*-test. *P ≤ 0.05. **G.** Expression of *WRKY45*, *WRKY53,* and *AT1G51890* immunity marker genes relative to *AtGAPDH* in WT (Col-0) and *athe-1* roots at 1, 3, and 7 dpt to Mock or Fo5176 *pSIX1:GFP*. Av ± SEM, *N* = 4 independent replicates from ≥ 20 roots. RM *t*-test. *P ≤ 0.05, **P ≤ 0.01, ***P ≤ 0.001, ****P ≤ 0.0001. Asterisks in black indicate the differences to WT Mock. Asterisks in blue indicate the differences of *athe-1* Fo5176 vs *athe-1* Mock. *<u>Of note</u>: optical cross sections shown in Fig 2E are also shown as Mock in Fig 3A*.

These cellular and subcellular results were confirmed at the whole root level in ATHE-GFP plants exposed to Fo5176 young hyphae for 1 day. Western blot analysis revealed a decrease in the total amount of ATHE-GFP in infected roots, while the total protein content of the MVB/PVC membrane localized SNARE protein SYP21 ^66^ remained unaltered (Fig. 3D and E). Furthermore, the ATHE-GFP protein level was partially recovered when the Fo5176 infection was performed in media supplemented with the clathrin-mediated endocytosis inhibitor ES9-17 ^67^ (Fig. S2C and D). These data confirm that the reduction of ATHE-GFP in response to the fungus is not a general response of membrane proteins and that its depletion from the PM is driven by clathrin-mediated endocytic processes triggered during Fo5176 infection.

In parallel, we assessed *ATHE* transcriptional regulation using a pATHE::mCherry-N7 line that targets mCherry to the nucleus via the N7 peptide to streamline signal quantification ^65^. Under mock conditions, *ATHE* expression was barely detectable in epidermal root cells of the transition-elongation and ED zones, but was clearly observed in the MD zone (Fig. S2 E and F), consistent with the gradient observed in ATHE protein levels (Fig. 2E, F, and G). Seedlings exposed to Fo5176 young hyphae showed a significant increase in *ATHE* expression in epidermal cells along all different root zones at 1 dpt that was notably enhanced at 3 dpt (Fig. S2 E and F). These data were further confirmed by qRT-PCR in whole roots at different times after fungal exposure (Fig. 3F), showing a 2-fold upregulation of relative *ATHE* expression at 7 dpt with Fo5176 spores. Our results showed that *ATHE* transcript levels gradually increased while the fungal infection progressed.

Taken together, our data revealed a reorganization of ATHE expression and protein localization in response to Fo5176 colonization, dependent on the infection stage. This reorganization may reflect increased protein turnover consistent with PM receptor dynamics upon signal perception.

### ATHENA participates in plant immune response to Fo5176 infection

As receptor-mediated signaling leads to transcriptional changes, we tested whether ATHE is involved in transcriptional changes of genes previously reported to be upregulated upon Fo5176 infection; *WRKY45*, *WRKY53* and *At1G51890* ^29,68,69^. Using plate-infection assays at 1, 3, and 7 dpt, we monitored transcriptional responses during hyphal surface contact, apoplast colonization, and vascular invasion ^17^. These assays confirmed previously reported transcriptional changes and provided higher temporal resolution. In WT roots, *WRKY45* was induced at 1 and 7 dpt, *WRKY53* increased at all time points, and *At1G51890* was upregulated only at 1 dpt (Fig. 3G). In *athe-1*, *WRKY45* and *WRKY53* showed elevated basal expression under all mock conditions. (Fig. 3G). ATHE loss also altered Fo-infection-induced transcription in a stage-dependent manner. At 1 dpt, *athe-1* failed to induce *WRKY45*. At 3 dpt, only *WRKY53* was induced, and its induction was weaker than in WT. At 7 dpt, all three genes responded similarly to WT, but the magnitude of induction remained markedly reduced, compared to the basal upregulation in *athe-1* in mock conditions. Together, these results show that ATHE modulates a subset of Fo5176-induced defense genes in a spatially and temporally dependent manner. Loss of ATHE elevates basal expression yet dampens or prevents proper activation of these genes during infection.

### ATHENA forms a complex with MIK2 in response to Fo5176 infection

Frequently, RKs exert their function interacting with other PM receptors to trigger the appropriate cellular responses ^70^. To identify possible ATHE-partners participating in its role in plant defense against Fo5176, we performed a pull-down assay followed by LC-MS/MS with ATHE-GFP roots upon mock and Fo5176 plate-infection conditions at 3 dpt. The Lti6B-GFP PM-marker was included as the background-binding control. As a result, 14 proteins were found to be specifically enriched in ATHE-GFP IP but not in Lti6B-GFP IP upon Fo5176 infection (Table S2). Of these 14 proteins, only 2 were enriched in Fo5176-infected compared to mock-treated roots (Table S2). Of these, we selected the LRR-RK MIK2 for further characterization, as it was previously shown to contribute to plant defense against Fo5176 and to responses to CW perturbations ^22,24^, functions that align with those we identify and propose for ATHE. The association between ATHE and MIK2 was confirmed by co-immunoprecipitation assays using whole mock- or Fo5176-treated roots of the dual marker line *athe-1* ATHE-GFP - *mik2-1* MIK2-mCh (*mik2-1* pMIK2:MIK2-mCherry from ^25^). As we previously showed, ATHE protein levels decreased upon Fo5176 infection (Fig. 3D, E, and 4A). Notably, despite the reduced amount of immunoprecipitated ATHE-GFP, more MIK2-mCh was recovered in association with ATHE-GFP during infection than under mock conditions (Fig. 4A and B). Importantly, MIK2-mCh protein levels in the input fraction were unaffected by Fo5176 infection (Fig. 4A, Input), indicating that increased binding was not due to changes in MIK2-mCh abundance. As a control, we performed the same co-immunoprecipitation assay to test the interaction between ATHE and the CrRLK1L FER, previously reported to participate in plant responses to Fo5176 and CW perturbations ^29,36,71^. Using a FER-specific antibody ^72^, we detected no FER protein in the ATHE-GFP-immunoprecipitated fraction (Fig. 4C and S3A). To further examine the ATHE-MIK2 interaction at the subcellular level, we imaged MD root epidermal cells in which ATHE-GFP foci remained visible at the PM during Fo5176 infection. In mock-treated plants, MIK2-mCherry localized to the PM in foci similar to those observed for ATHE-GFP (Fig. 4D). Notably, the basal colocalization of these receptors in mock-treated cells was significantly enhanced in cells contacting young Fo5176 hyphae for 1 day (Fig. 4D, 4E, S3B, and S3C). These microscopy and biochemical data indicate Fo5176 infection leads to an increased ATHE-MIK2 interaction.

**Figure 4.**
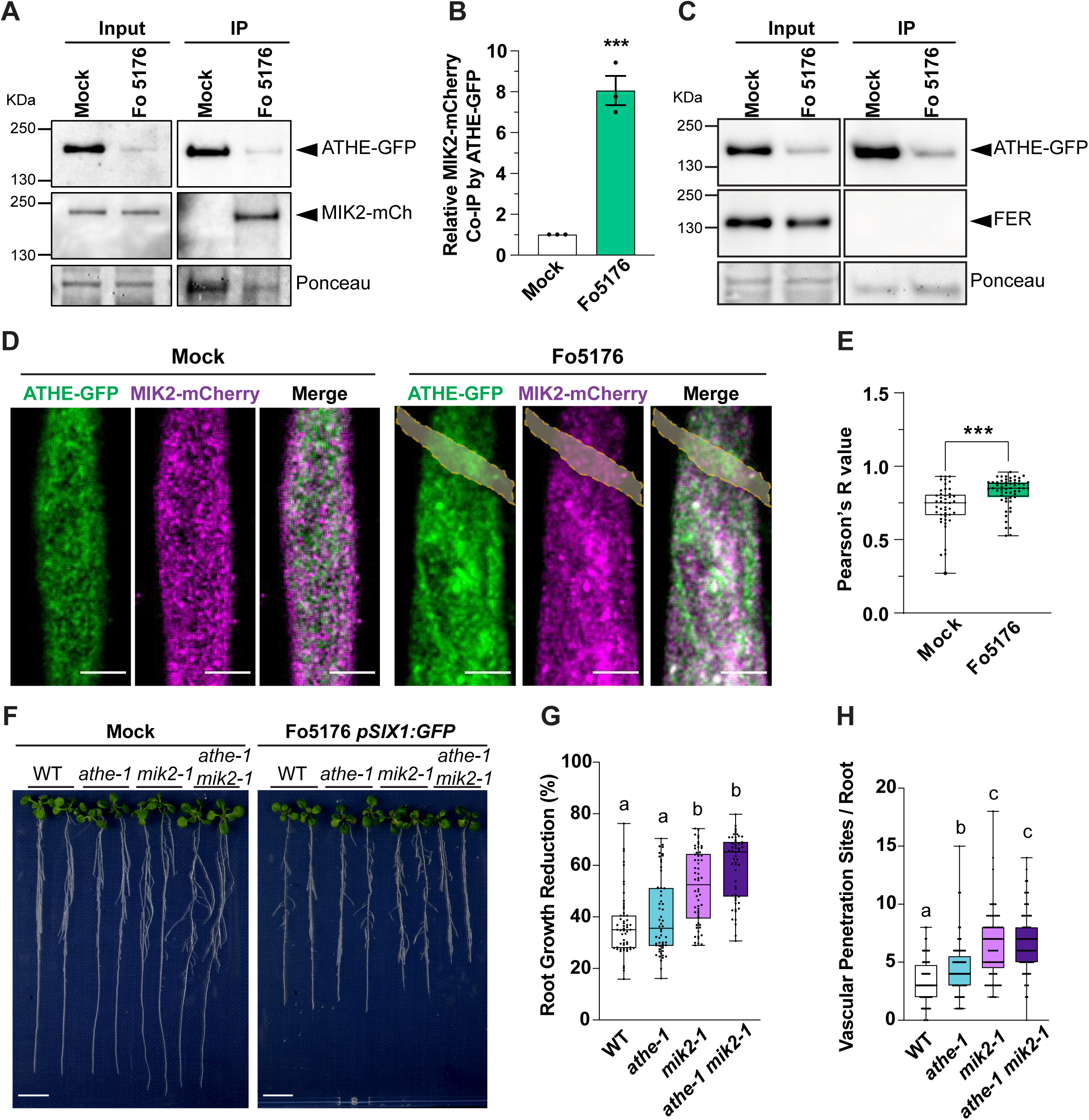
ATHE interacts with MIK2 during Fo5176 infection. **A.** Representative Western blot co-immunoprecipitation analysis of ATHE-GFP and MIK2-mCherry using GFP-trap beads, in Mock or Fo5176-infected roots at 1 day post treatment (dpt), using anti-GFP (ATHE-GFP) and anti-RFP (MIK2-mCherry) antibodies. Ponceau S staining provides loading control. **B.** ATHE-GFP and SYP21 co-immunoprecipitation quantification from immunoblots as in (A), normalizing the MIK2-mCherry intensity band to that of the ATHE-GFP on each condition. Av ± SEM, *N* = 3 independent experiments. Repeated Measures (RM) *t*-test. ***P ≤ 0.001. **C.** Representative Western blot co-immunoprecipitation analysis of ATHE-GFP and FERONIA (FER) using GFP-trap beads, in Mock or Fo5176-infected roots at 1 dpt, using anti-GFP (ATHE-GFP) and anti-FER antibodies. Ponceau S staining provides loading control. **D.** Representative spinning disk confocal images of ATHE-GFP (green) and MIK2-mCherry (magenta) in root epidermal cells at the MD zone at 1 dpt with Mock or Fo5176 young hyphae (yellow). Scale bars = 5 µm. **E.** ATHE-GFP and MIK2-mCherry colocalization in roots as in (D). Av ± SEM, *N* ≥ 44 cells of 10 roots/ treatment. Box plots: centerlines show the medians; box limits indicate the 25th and 75th percentiles; whiskers extend to the minimum and maximum. RM *t*-test. ***P ≤ 0.001. **F.** Representative images of WT, *athe-1*, *mik2-1*, and *athe-1 mik2-1* mutants at 6 dpt with Mock or Fo5176 *pSIX1:GFP*. Scale bars = 1 cm. **G.** Root growth reduction of Fo5176 *pSIX1:GFP*-infected WT, *athe-1*, *mik2-1*, and *athe-1 mik2-1* plants relative to Mock-treated ones at 6 dpt, as in (F). Box plots as described in (E). *N* ≥ 55 roots from 3 experiments (≥ 15 roots/experiment). RM one-way ANOVA with Kruskal-Wallis multiple comparison test. Alphabet letters indicate significant differences with P ≤ 0.05. **H.** Cumulative number of Fo5176 *pSIX1:GFP* vascular penetrations per root in WT, *athe-1*, *mik2-1*, and *athe-1 mik2-1* plants at 7 dpt. Box plots as in (E), *N* ≥ 72 roots from 3 experiments (≥ 17 roots/experiment). RM one-way ANOVA with Kruskal-Wallis multiple comparison test. Alphabet letters indicate significant differences with P ≤ 0.05.

To further understand the biological significance of the ATHE-MIK2 interaction at the PM, we assessed ATHE-GFP abundance at the PM in the *mik2-1* mutant. We found that the absence of MIK2 resulted in a significantly reduced ATHE-GFP signal at the PM under mock conditions, which remained unchanged upon Fo5176 contact (Fig. S3D and E). These observations indicate that MIK2 is required for proper ATHE localization at the PM in the absence of stress and appears to be necessary for its Fo5176-triggered internalization.

We then assessed the potential functional redundancy between ATHE and MIK2 during Fo5176 infection using plate-based assays of the *athe-1 mik2-1* double mutant and its parental lines. Our data show that *mik2-1* roots exhibited stronger growth inhibition and a higher frequency of fungal vascular penetrations compared with WT (Fig. 4F–H), consistent with and extending previous findings ^23^. The *athe-1 mik2-1* double mutant displayed a susceptibility phenotype indistinguishable from *mik2-1*, both in terms of root growth inhibition and vascular penetration events, and these responses were markedly stronger than those observed in *athe-1* alone (Fig. 4F–H).

Taken together, the biochemical, imaging, and genetic evidence identify MIK2 as an upstream regulator required for ATHE localization and function, with both receptors forming an infection-reinforced complex that operates in a shared pathway to restrict Fo5176 colonization

### ATHENA participates in plant response to cellulose alterations

As noted above, ATHE contains an extracellular domain containing MLD and LRR domains (Fig. S1B), also found in other RKs involved in responses to Fo5176 infection and CW perturbation, including FER and THE1 ^21,22,34,73–75^. During Fo5176 infection, inhibition of cellulose synthesis constitutes a key CW-remodeling response that helps counteract fungal colonization ^17^, and we show that this occurs across distinct developmental regions of the root (Fig. S4A). To assess whether ATHE contributes to this process, we used isoxaben (ISX), a cellulose-synthesis inhibitor commonly employed to mimic CW stress ^76–78^. ISX acts primarily in dividing and elongating cells where primary-wall cellulose synthases are enriched, which we also confirmed in our system (Fig. S4B, C). To determine ATHE’s contribution to the ISX response, we quantified root-growth inhibition after 5 dpt (Fig. 5A). ISX-induced root-growth inhibition was significantly attenuated in *athe-1*, whereas the ATHE-GFP complemented line responded similarly to WT. At the cellular level, 1 day of ISX treatment triggered an approximately three-fold increase in ATHE-GFP signal in ED cells, where the protein is normally undetectable (Fig. 5B and C). By contrast, ATHE-GFP abundance remained unchanged in MD cells, where ATHE is already present under mock conditions (Fig. 2E–G). These results are consistent with the weaker effects of ISX in this root region (Fig. S4B and C). Western blot analysis confirmed a global increase in total ATHE-GFP protein in roots following ISX treatment (Fig. S4F and G). We next examined whether this increase in ATHE protein was accompanied by transcriptional upregulation. Using the pATHE::mCherry-N7 reporter, we detected a strong induction of the activity of *ATHE* promoter in TZ-EZ and ED epidermal cells of ISX-treated roots compared to mock conditions, with MD cells also showing a more moderate increase (Fig. 5D). qRT-PCR further validated this transcriptional activation, revealing a 2.3-fold rise in *ATHE* transcript levels after 1 day of ISX exposure (Fig. 5E).

**Figure 5.**
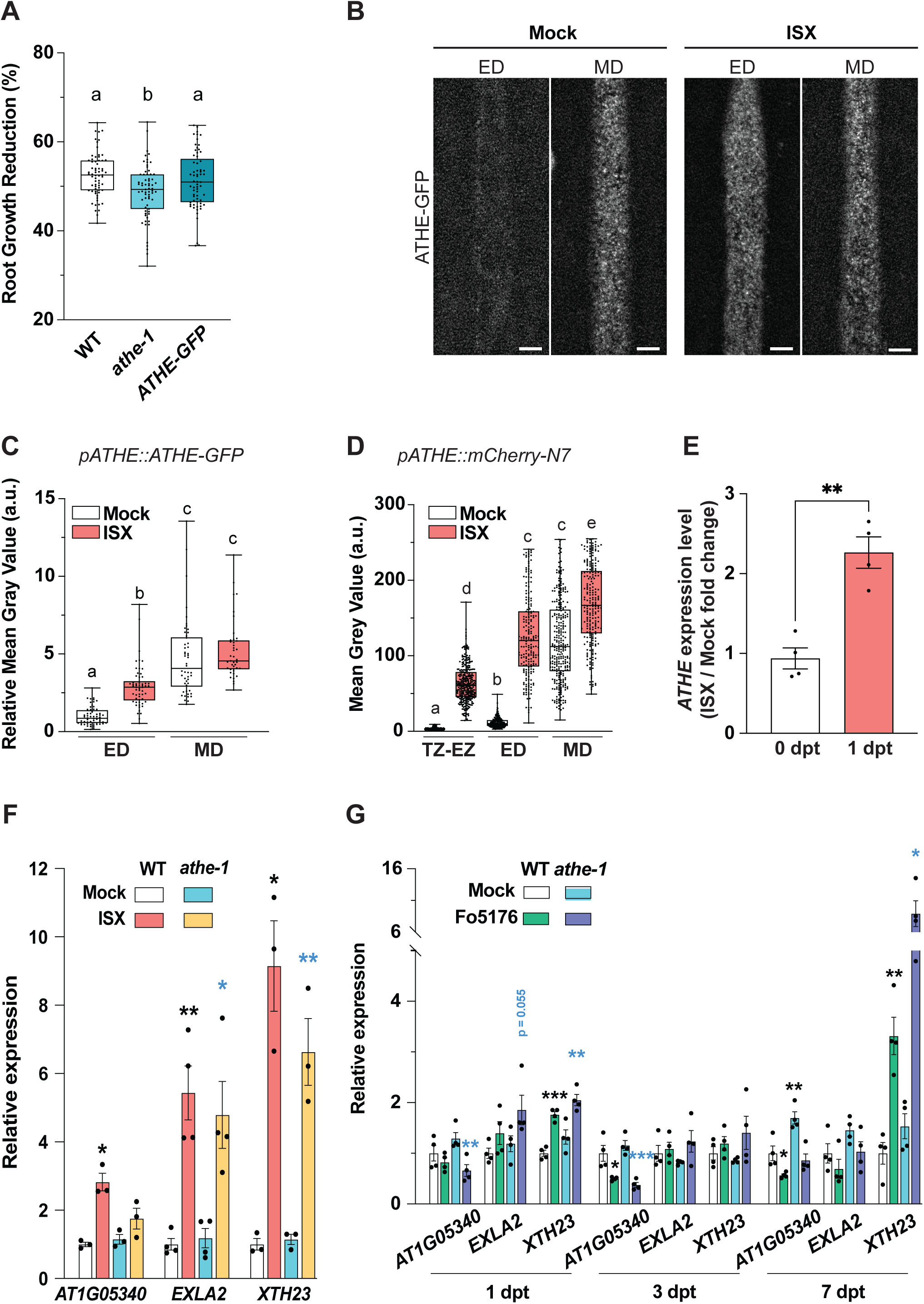
ATHENA participates in plant response to isoxaben. **A.** Root growth reduction of 2nM isoxaben (ISX)-treated wild type (WT; Col-0), *athe-1*, and ATHE-GFP seedlings relative to Mock-treated ones, after 5 days post treatment (dpt). The box plots are shown: centerlines show the medians; box limits indicate the 25th and 75th percentiles; whiskers extend to the minimum and maximum. *N* ≥ 65 roots from 4 experiments (≥ 15 roots/experiments). Repeated measures (RM) one-way ANOVA with Dunn’s multiple comparison test. Alphabet letters indicate significant differences with P ≤ 0.05. **B.** Representative spinning disk confocal images of ATHE-GFP in epidermal cells of the ED and MD in response to 300 nM ISX for 1 day. Scale bars = 5 μm. **C.** Amount of ATHE-GFP at the plasma membranes of cells as described in (B), quantified as mean gray value (arbitrary units, a.u.) of ISX-exposed samples normalized to their corresponding Mock-treated ED. Box plots as described in (A). *N* ≥ 41 cells were obtained from 3 experiments (3-5 cells/root and ≥ 3 roots/experiments). RM one-way ANOVA with Dunn’s multiple comparison test. Alphabet letters indicate significant differences with P ≤ 0.05. **D.** Amount of mCherry signal intensity of *pATHE:mCherry-N7* epidermal cells at the root differentiated zones indicated in Fig 2(D) in response to Mock or 300 nM ISX treatments for one day, quantified as mean gray value (arbitrary units; a.u.). Box plots as in (A), *N* ≥ 208 cells were obtained from 3 experiments (≥ 6 cells/root and 6 roots/experiment were imaged). RM one-way ANOVA with Dunn’s multiple comparison test. Alphabet letters indicate significant differences with P ≤ 0.05. **E.** *ATHE* expression relative to *AtGAPDH* in WT roots after 1 day post treatment of 300 nM ISX normalized to Mock-treated ones. Av ± SEM, *N* = 4 independent experiments (≥ 20 roots/experiment). RM *t*-test compared to the fold change at 1 dpt. **P ≤ 0.01. **F.** Expression of the ISX-response genes *AT1G05340*, *EXLA2,* and *XTH23* relative to *AtGAPDH* in WT and *athe-1* roots at 1dpt with Mock or 300 nM ISX **G.** Gene expression of the ISX-response genes *AT1G05340*, *EXLA2,* and *XTH23* relative to *AtGAPDH* in WT and *athe-1* roots in at 1, 3 and 7 dpt with Mock or Fo5176 spores. In (F) and (G) Av ± SEM, *N* = 4 independent replicates (≥ 20 roots/experiment). RM *t*-test. *P ≤ 0.05, **P ≤ 0.01, ***P ≤ 0.001, ****P ≤ 0.0001. Asterisks in black indicate the differences of the corresponding sample vs WT Mock. Asterisks in blue indicate the differences in *athe-1* Mock versus *athe-1* treatment (ISX or Fo5176).

By weakening cellulose deposition, ISX reduces CW resistance to turgor, generating mechanical stress ^71,79,80^, and it also reorganizes the cortical microtubule (MT) array in the root (Fig. S4D, E) ^81–83^. To determine whether ATHE participates in plant response to alterations in MT stability, we treated roots with the MT-depolymerizing drug oryzalin (Ory) ^84^. Ory disrupts MT organization and consequently lowers the density of cellulose synthase complexes at the PM, an effect comparable to that induced by ISX (Fig. S4B-E). Remarkably, *athe-1* roots were insensitive to Ory-induced growth inhibition, and both ATHE-GFP abundance at the PM and ATHE promoter activity increased in MD cells at 1 dpt (Fig. S4H–J). These results parallel those obtained with ISX, supporting a role for ATHE in sensing disturbances in the cellulose synthesis machinery that generate mechanical stress. Finally, to exclude the possibility that ATHE responds to general osmotic changes rather than CW-related perturbations, we examined ATHE protein and transcript levels under sorbitol-induced osmotic stress. No changes in *athe-1* root growth sensitivity, ATHE-GFP abundance at the PM, or *ATHE* expression patterns across root zones were detected (Fig. S5), ruling out a nonspecific osmotic response.

Because ISX and Ory elicited similar effects on ATHE-related processes, we focused on ISX to further characterize ATHE’s role in CW-derived changes. To investigate ATHE’s involvement in the transcriptional responses to cellulose synthesis inhibition, we quantified the expression of the ISX-responsive genes *At1G05340, EXPANSIN-LIKE PROTEIN 2 (EXLA2),* and *XYLOGLUCAN ENDOTRANSGLUCOSYLASEzHYDROLASE 23* (*XTH23),* ^85–88^ in WT and *athe-1* plants. Basal transcript levels were comparable between genotypes (Fig. 5F). As expected, all three genes were strongly induced by ISX in WT roots. In contrast, ISX-induced expression of *At1G05340* and *XTH23* in *athe-1* was attenuated in relative to WT (Fig. 5F). Because Fo5176 induces CW-related stress similar to ISX, we also monitored gene expression during Fo5176 plate infection. Reflecting the parallelism between Fo5176 and ISX responses, WT roots upregulated *XTH23* after Fo5176 exposure, while *athe-1* roots exhibited a reduced induction trend at 1 dpt (Fig. 5G). Moreover, *At1G05340* was significantly downregulated in Fo5176-infected *athe-1* roots (Fig. 5G).

MIK2 has also been implicated in the Arabidopsis response to ISX ^22,26,71^. We therefore examined the genetic interaction between ATHE and MIK2 in sensing this CW perturbation. *mik2-1* roots were less sensitive than *athe-1* to ISX treatment, and notably, the *athe-1 mik2-1* double mutant restored ISX sensitivity to WT levels (Fig. S6A). In addition, similar to what we observed for ATHE, MIK2 protein levels increased upon ISX exposure in whole-root extracts by western blot (Fig. S6C). ISX treatment also enhanced ATHE-MIK2 association detected by co-immunoprecipitation (Fig. S6C and D). Overall, these results suggest that while cellulose synthesis deficiency influences ATHE–MIK2 association, its impact appears more moderate compared to the strong enhancement observed during Fo5176 infection (Figs. 4 and S6D).

Fo5176 also perturbs plant cellulose by degrading it and releasing cellulose-derived oligosaccharides (COSs) such as cellobiose, cellotriose, and cellohexaose ^12^, which serve as DAMPs that may be sensed by ATHE. Root growth of *athe-1* was significantly less inhibited by cellobiose and cellotriose than that of WT, consistent with its reduced sensitivity to ISX and Ory (Figs. S7A, 5A, and S4H). In agreement with these observations, *mik2-1* roots were also less sensitive to cellobiose and cellotriose, and COS sensitivity was restored to WT levels in the *athe-1 mik2-1* double mutant (Fig. S7A), indicating a non-additive interaction with MIK2 in the perception of cellulose-derived perturbations. Given that COSs induce a rapid (∼25 min) but transient upregulation of *WRKY30* and *WRKY40* that returns to baseline within 1 day in WT plants ^69^, we investigated whether ATHE influences this early transcriptional response. As reported, COS treatment induced both genes in WT roots, whereas this induction was reduced in *athe-1* when compared with their respective mock controls (Fig. S7B). Notably, in *athe-1*, *WRKY30* expression remained elevated at 1 dpt with cellobiose or cellotriose exposure, whereas it returned to basal levels in WT roors (Fig. S7B), suggesting that ATHE attenuates prolonged *WRKY30* activation triggered by COS perception. Overall, the enhanced defense-gene activation in *athe-1*, reduced ISX- and Fo5176-responsive transcriptional changes, and prolonged COS-induced *WRKY30* expression support a role for ATHE as an integrator of cellulose-status cues, defense pathways, and mechanical-stress responses. However, external application of cellobiose or cellotriose did not significantly alter ATHE-GFP PM abundance or *ATHE* expression along the root axis (Fig. S7C–E), suggesting that ATHE is not directly stabilized or transcriptionally induced by COSs. Recent structural data revealed that the extracellular domain of the MD-LRR–containing RK IGP1/CORK1 binds cello-oligomers through a highly specific sugar-binding pocket within the LRR domain, whose structural integrity is reinforced by the MD ^63,89^. These structural insights confirm previous observations from ITC binding experiments ^62^. Using the same experimental approach, we assessed whether ATHE directly binds COSs *in vitro*. Titration of cellobiose or cellotriose into solutions containing the ATHE extracellular domain (ATHE-ECD) did, however, not reveal any detectable interaction (Fig. S7F and G).

Together, these findings show that ATHE contributes to root responses triggered by CW-perturbing treatments that impose both mechanical and chemical stress, such as ISX, Ory, cellobiose, and cellotriose.

### ATHENA participates in plant response to Fo-RALF

To further investigate the functional relationship between ATHE and MIK2 in plant immunity, we tested whether ATHE contributes to the plant response to SCOOP12, a potent SCOOP peptide perceived by MIK2 ^23,25^. As expected, the *mik2-1* mutant was insensitive to SCOOP12, whereas *athe-1* exhibited root growth inhibition comparable to the WT upon peptide treatment (Fig. S8). Consistently, SCOOP12-induced root growth inhibition in the *athe-1 mik2-1* double mutant was indistinguishable from that of the *mik2-1* single mutant (Fig. S8). Together, these results indicate that ATHE is not required for this MIK2-mediated response to SCOOP12.

Together with SCOOPs, RALF peptides have been shown to play key roles in modulating immune responses, cell expansion, and CW remodeling. In addition, Fo secretes a RALF-like (Fo-RALF) to suppress host immunity hijacking FER ^29,90^. Because FER and THE1, which binds RALF34 to regulate growth upon cellulose biosynthesis inhibition ^91^, are RKs containing a MLD in their ectodomains, as ATHE, we next examined whether ATHE is required for plant responses to RALF peptides and its potential genetic interaction with MIK2 in this process. In addition to Fo5165-RALF (Fo-RALF), we included in our analysis the root-specific peptides RALF1 and RALF23, representatives of the clades most closely related to Fo-RALF and perceived by FER. We also analyzed RALF34, which belongs to a more distantly related clade ^30^. We confirmed the sensitivity of WT roots to both plant and fungal RALFs ^90,92^. In contrast, *athe-1* retained sensitivity to endogenous RALFs but was almost insensitive to Fo-RALF (Fig. 6A), indicating that ATHE specifically mediates Fo-RALF perception. Both *mik2-1* and the *athe-1 mik2-1* double mutant showed reduced sensitivity to Fo-RALF relative to WT, and their responses were consistently weaker than those of *athe-1* (Fig 6A). These results indicate that ATHE contributes to the plant response to Fo-RALF, while MIK2 acts epistatically to ATHE and has a comparatively smaller impact on this response.

**Figure 6.**
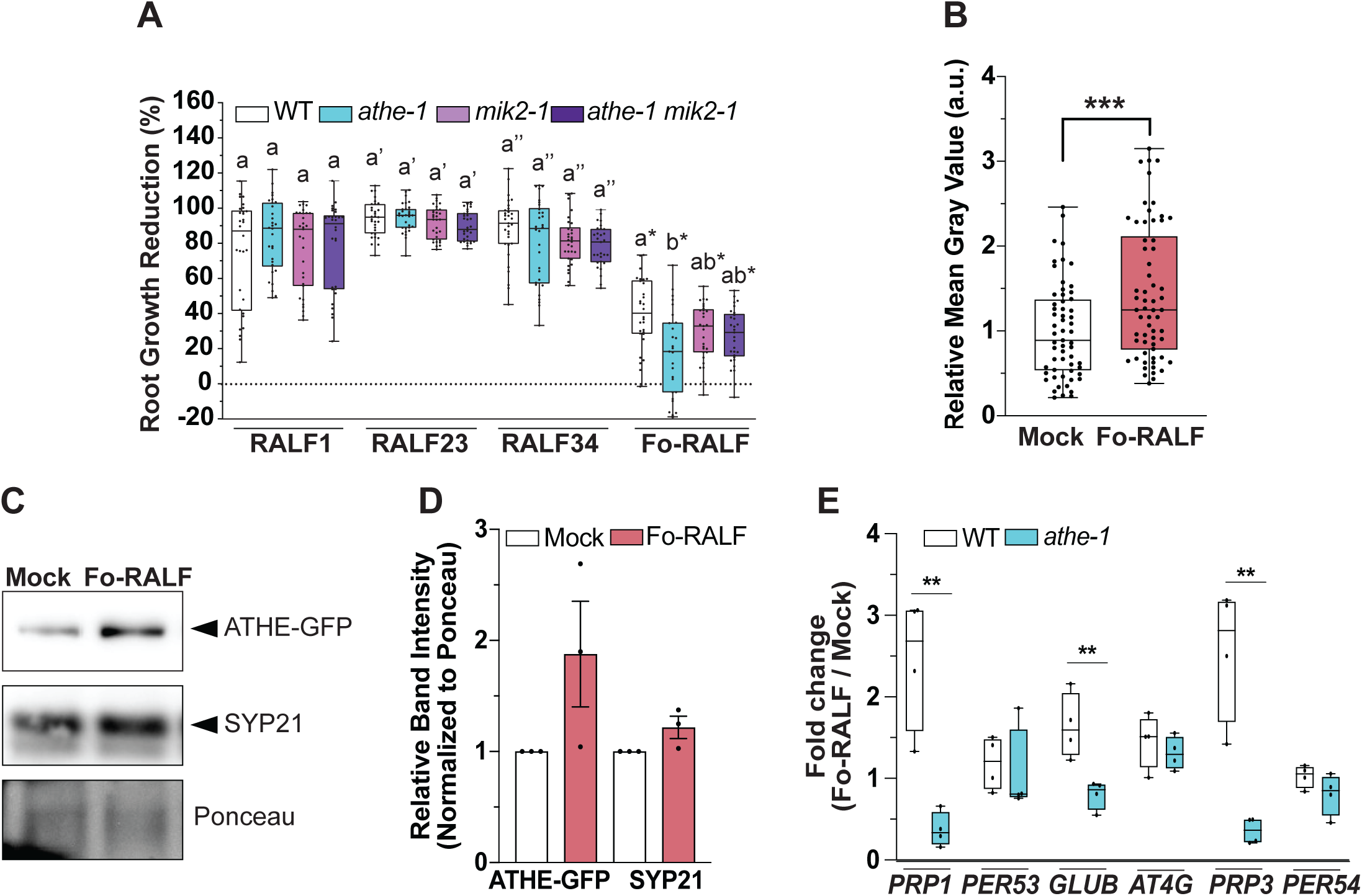
ATHENA participates in plant response to Fo-RALF. **A.** Root growth reduction of 2 µM RALF1, RALF23, RALF34, or Fo-RALF peptides-treated wild type (WT; Col-0), *athe-1, mik2-1*, and *athe-1 mik2-1* seedlings relative to Mock-treated ones, after 3 days post-treatment (3 dpt). The box plots are shown: centerlines show the medians; box limits indicate the 25th and 75th percentiles; whiskers extend to the minimum and maximum. *N* ≥ 30 roots from 3 independent experiments (*N* of roots = 10-15/experiment). Repeated Measures (RM) one-way ANOVA with Dunnett’s correction. Alphabet letters with different symbols (′, ″, or *) indicate significant differences within each treatment with P ≤ 0.05. **B.** Amount of ATHE-GFP at the plasma membrane of cells, quantified as mean gray value (arbitrary units, a.u.) of Fo-RALF-exposed samples normalized to their corresponding Mock. Box plots as in (A). *N* ≥ 58 cells from ≥ 3 experiments (5-6 cells per root, 3-4 roots per replicate, 3 replicates). RM t-test. ***P ≤ 0.001. **C.** Representative Western blot of total ATHE-GFP and SYP21 proteins in 8-day-old ATHE-GFP roots at 1dpt with Mock or Fo-RALF 2 µM. Ponceau S staining provides loading control. **D.** ATHE-GFP and SYP21 signal quantification from immunoblots as in (C). For each protein, the intensity of the band was first normalized to its corresponding Ponceau signal. Then, the value for the Fo-RALF-exposed sample was normalized to that of the Mock. Av ± SEM, *N* = 3 independent experiments. RM *t*-test. **E.** Expression of the Fo-RALF-responsive genes *PRP1, PER53, GLUB, AT4G16260, PRP3,* and *PER54* relative to *AtGAPDH* in WT and *athe-1* roots at 1dpt with Mock or 2 µM Fo-RALF. Box plots as in (A). *N* = 3 independent replicates (≥ 20 roots/replicate). RM t-test. **P ≤ 0.01.

To gain insight into the putative role of ATHE in sensing Fo-RALF, we imaged ATHE-GFP in plants exposed to this peptide for 1 day. Fo-RALF induced a clear increase in ATHE-GFP signal intensity in MD cells compared with the mock control (Fig. 6B). Accordingly, whole roots exposed to Fo-RALF for 1 day exhibited increased ATHE-GFP protein accumulation, whereas the abundance of the endomembrane marker SYP21 remained unchanged relative to mock-treated controls (Fig. 6C and D). To determine whether Fo-RALF modulates plant immunity, and whether ATHE contributes to this process, we analyzed the expression of defense-related genes known to be upregulated during Fo infection ^29,93^. We focused on genes associated with reinforcement of apoplastic barriers: *PRP1* and *PRP3* (encoding proline-rich proteins involved in CW assembly and reinforcement) ^94^, *GLUB* (encoding a germin-like protein contributing to apoplastic reactive oxygen species production) ^95^, *AT4G16260* (*AT4G*, encoding a stress-responsive gene previously linked to pathogen infection) ^96^, and *PER53* and *PER54* (encoding class III peroxidases associated with lignification and reactive oxygen species metabolism) ^97^. We detected that in WT plants, Fo-RALF triggered a significant induction of all tested genes except *PER54* (Fig. 6E). In contrast, Fo-RALF also failed to induce *PRP1*, *PRP3,* or *GLUB* in the *athe-1* mutant (Fig. 6E). Because these genes are associated with CW strengthening and oxidative metabolism, their reduced activation in *athe-1* suggests that ATHE is required for Fo-RALF-mediated reinforcement of apoplastic barriers and defence-related oxidative processes.

Overall, our data reveal that ATHE is dispensable for the MIK2-dependent response to SCOOP12 but is essential for response to Fo-RALF. In addition, we show that ATHE1 is required for Fo-RALF-induced activation of defence-related genes involved in CW reinforcement and apoplastic oxidative processes.

### Ectopic ATHENA expression in tomato leads to increased resistance to Fol

Our data show that ATHE is required for Arabidopsis defense against Fo5176 and Rs infection (Figs. 1C and S1C-F). Thus, we aimed to identify orthologs of this protein in other plant lineages. Protein sequences most similar to ATHE were identified, aligned, and subjected to phylogenetic analysis. The resulting tree generated using ATHE and its closest homologs from diverse species, showed that ATHE is conserved within *Brassicaceae*, while no orthologous sequences were detected in more distantly related taxa, including the *Solanaceae* plant species *Solanum lycopersicum* (Fig. S9). These results indicate that ATHE is a lineage-specific protein restricted to *Brassicaceae,* making it an interesting candidate for cross-family gene transfer in crop-disease management. To evaluate this potential, we generated independent *S. lycopersicum* transgenic lines constitutively expressing *AtATHE* in the Marmande cultivar (Fig. 7A). These lines exhibited significantly higher survival rate when infected by Fol007 than the WT plants (Fig. 7B-C). These results demonstrate that AtATHE enhances resistance to Fol infection in tomato, supporting its conserved role in Fo-related immune responses and underscoring its potential for biotechnological applications in crop protection.

**Figure 7.**
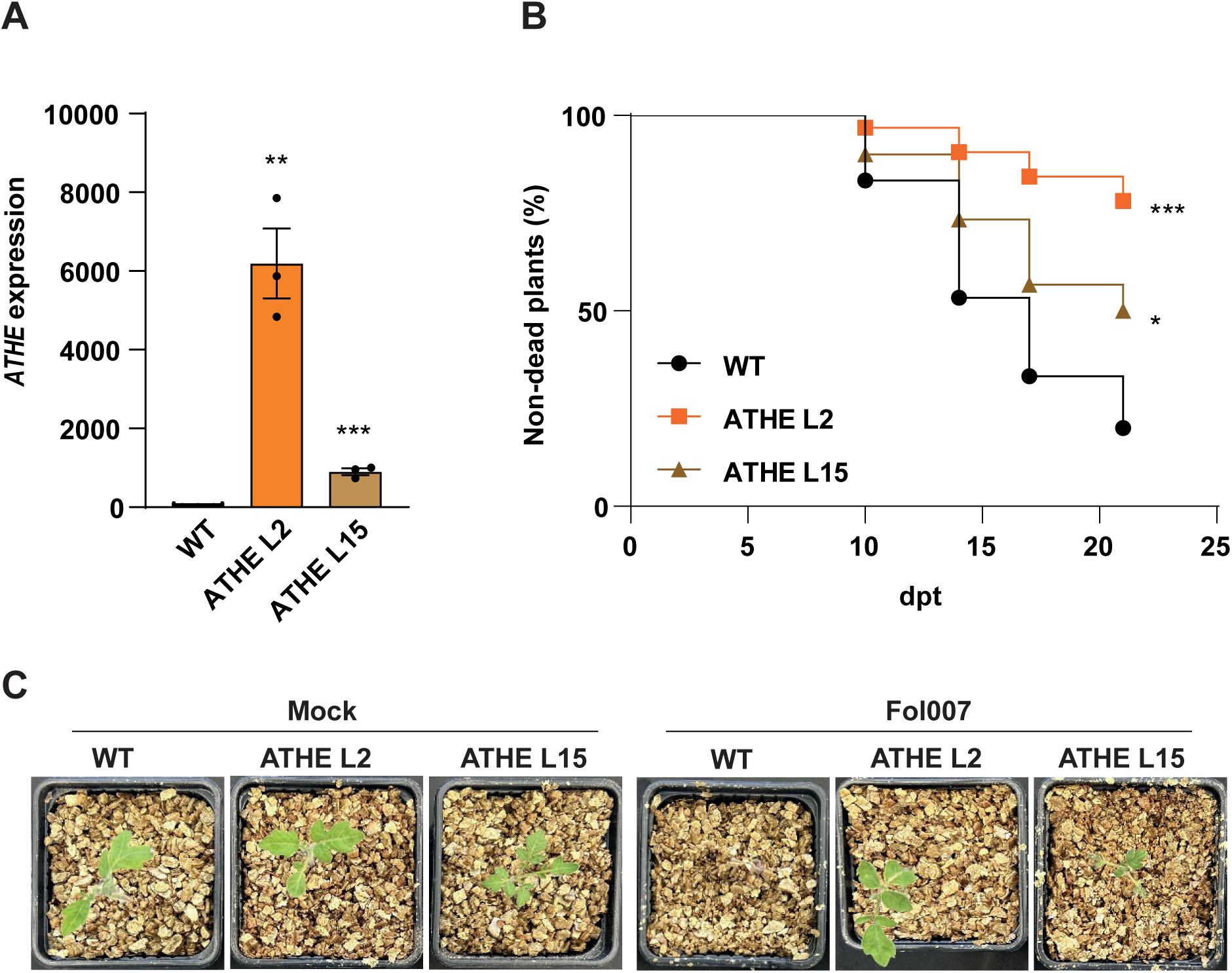
ATHE expression in tomatoes increases their resistance to Fo. **A.** *ATHE* expression in WT (Marmande) and two independent lines expressing *ATHE* (L2 and L15) relative to *ACTIN*. Av ± SEM, *N* = 3 independent replicates. Repeated Measures (RM) t-test. **P ≤ 0.005; ***P ≤ 0.0005. **B.** Representative Kaplan-Meier plot showing the non-dead tomato plants at different days post-treatment (dpt) with Fol007 microconidia. *N* = 30 plants/genotype. *P ≤ 0.05; ***P ≤ 0.0005 versus WT according to the log-rank test. **C.** Representative images of WT, ATHE L2, and ATHE L15 plants at 17 dpt with Fol007.

## DISCUSSION

Plant CWs constitute both a structural barrier and a dynamic sensory interface that enables plants to detect microbial invasion ^8^. Although CWI signaling is essential for coordinating immune activation during pathogen attack, the receptors that perceive CWI alterations during early infection, particularly by vascular pathogens, have remained poorly resolved. Here, we identify ATHE/MEE39 as a previously uncharacterized Mal-LRR-RK that functions as an infection-responsive CWI sensor and restricts colonization by both *Fusarium oxysporum* Fo5176 and *Ralstonia solanacearum*. Our data reveal that ATHE undergoes rapid remodeling of abundance, localization, and trafficking upon pathogen contact and that its function depends on a stress-reinforced partnership with the LRR-RK MIK2.

Our initial observation that mutants defective in CME early adaptor complexes are more susceptible to Fo5176 establishes CME as an essential regulator of root defense. Live-cell imaging of TPLATE-complex lifetimes and endocytosis tracking demonstrated that Fo-derived signals rapidly increase CME activity (Fig. 1). Because PRR internalization is required for its activation, attenuation, and endosomal signaling ^46^, these data underscore that CME is not merely a housekeeping process but an active component of plant immunity. CME responsiveness also provides a mechanistic rationale for isolating receptors internalized during early pathogen contact, which led to the identification of ATHE/MEE39 from the MVB/PVC proteome of Fo5176-infected roots (^64^; Fig. 2A and Table S1).

ATHE’s architecture is consistent with a PM receptor dedicated to monitoring CW-associated signals and structural perturbations. Consistent with this potential role, ATHE localizes to punctate PM foci in epidermal and cortical cells and exhibits a striking developmental gradient, with minimal abundance in early differentiating cells and progressive enrichment toward the late differentiation zone, precisely the region where Fo5176 typically initiates penetration (Fig. 2 D-G; ^17^). Fungal contact triggers rapid, CME-dependent depletion of ATHE from the PM, coupled with strong transcriptional upregulation across both early and late differentiation zones (Figs. 3 A-D, and S2). Such reciprocal regulation, internalization combined with transcriptional activation, is characteristic of ligand-activated PRRs and suggests that ATHE is actively engaged during the early stages of pathogen sensing. This spatial pattern echoes recent transcriptomic studies showing that MAMP exposure induces layered transcriptional reprogramming across root cell types in response to damage ^98–102^. Our findings extend these observations to the receptor level, revealing that ATHE localization is not homogenous nor static but undergoes dynamic redistribution during infection. The spatial and temporal specificity of this response implies that ATHE contributes to Fo5176 defense primarily in the outer root layers, where initial hyphal exploration occurs. Moreover, the requirement for apoplastic colonization to activate ATHE-dependent responses indicates that sufficient fungal-derived cues accumulate only after hyphae establish themselves in the apoplast. This timing argues against early PM-proximal pH shifts, reported immediately after Fo5176 contact ^103^, as the primary trigger of ATHE signaling, pointing instead to apoplastic signals associated with CW disturbance or pathogen growth.

Functionally, loss of ATHE compromised root defense at multiple levels. *athe-1* mutants exhibited significantly increased fungal entry into the vasculature in plate-based assays and displayed heightened wilting susceptibility in soil (Figs. 2B-C and S1B-D), indicating a failure to restrict early hyphal progression. This susceptibility also extended to the bacterial vascular pathogen *Ralstonia solanacearum* (Fig. S1E and F), demonstrating that ATHE contributes to immunity against two taxonomically unrelated wilt pathogens. Beyond limiting microbial ingress, ATHE also influenced Fo-induced transcriptional reprogramming. Thus*, athe-1* roots accumulated elevated basal levels of several defense-related transcripts yet failed to appropriately induce or sustain their activation during Fo infection (Fig. 3G). This dysregulated basal expression together with reduced pathogen-induced transcription indicates that ATHE functions early in immune signaling. Loss of ATHE disrupts the coordination between basal immune gene homeostasis and inducible defense activation, diminishing both preparedness and response amplitude. Similar signaling roles have been described for RFO1/WAKL22, another PM-RK that also detects CW modifications ^39^. Together, these findings support a model in which ATHE acts as a regulatory hub that calibrates root immunity, suppressing inappropriate basal activation while ensuring robust defense signaling during infection progression.

A central mechanistic finding is that ATHE forms an infection-reinforced complex with MIK2, an LRR-RK required for plant response to SCOOP peptides, CWI stress, and Fo5176 ^26^. Although both receptors colocalize at the PM under basal conditions, their association increases markedly during infection (Fig. 4A-E). Importantly, MIK2 is required for proper ATHE accumulation at the PM even without pathogen challenge, and *athe-1 mik2-1* double mutant shows susceptibility indistinguishable from *mik2-1* alone (Figs. 4 F-H and S3D and E). These results position MIK2 upstream of ATHE as a stabilizing or activating partner, and indicate that MIK2 acts epistatically to ATHE to limit Fo colonization. The strengthening of the ATHE-MIK2 complex during infection also suggests that the complex responds to pathogen-induced changes in CW composition, tension, or apoplastic environment.

Among the apoplastic cues generated during Fo infection, signals arising from pathogen-induced cellulose disruption ^12,17^ emerged as strong candidates for activation of the ATHE-MIK2 module. Consistent with this idea, perturbations that impair the cellulose-synthesis machinery, either through ISX-mediated inhibition or microtubule destabilization by Ory, (both of which alter wall mechanics in elongating and early-differentiating root cells (^71,76–80^; Fig. S4A-E), revealed a clear dependence on ATHE. In *athe-1*, these treatments caused reduced growth inhibition and diminished activation of CW-damage–associated genes, whereas WT roots displayed a pronounced increase in ATHE transcript and protein levels (Figs. 5 and S4H-J). Fo5176 itself induced the ISX-responsive gene *XTH23*, and ATHE contributed to this transcriptional activation, further linking ATHE to the perception of CW-altering cues. Importantly, osmotic treatments failed to trigger these responses (Fig. S5), indicating that ATHE monitors structural perturbations rather than general stress, an important distinction given that reduced cellulose content can indirectly affect cellular turgor ^86,104^. Together, these observations position ATHE as a key component of the machinery that interprets pathogen- and mechanically-induced modifications to the CW synthesis apparatus and channels these cues into appropriate immune outputs.

ATHE also contributed to the plant response to COSs, which function as DAMPs during Fo infection and activate PTI signaling ^12,62,69,105,106^. *athe-1* roots showed reduced sensitivity to cellobiose and cellotriose. A similar phenotype was observed in *mik2-1* roots, and COS sensitivity was restored to WT levels in the *athe-1 mik2-1* double mutant (Fig. S7A). Comparable responses to ISX were also detected in *mik2-1* and the double mutant (Figs. S6 and S7A), suggesting a non-additive relationship with MIK2 in perceiving CW-derived perturbations. This pattern is consistent with antagonistic functions within the same signaling pathway or with activation of a compensatory mechanism in the double mutant that reinstates the WT response. In addition, *athe-1* roots showed early COS-triggered gene activation and prolonged induction of *WRKY30* (Fig. S7B), suggesting that ATHE modulates DAMP-induced transcription rather than serving as a COS receptor. *WRKY30* overexpression has been linked to enhanced stress resilience during germination and leaf development ^107–109^, and our findings indicate that this transcription factor also supports root defense.

Unlike COSs, which alter downstream transcription without affecting ATHE protein abundance or localization (Fig. S7B-E), mechanical perturbations (induced by ISX or Ory) strongly induced ATHE and modified its subcellular distribution. Consistent with this distinction, ATHE showed no detectable binding to cellobiose or cellotriose *in vitro* (Fig. S7F and G). While the MLD-LRR architecture of ATHE shows no detectable binding activity, the LRR-MLD ectodomain of IGP1/CORK1 binds cellotriose ^63^. This difference suggests that the relative positioning of the MLD and LRR domains may influence ligand specificity, a possibility that warrants further experimental testing. Together, these results indicate that ATHE integrates mechanical and biochemical inputs arising from cellulose perturbation. Overall, ATHE contributes to root responses triggered by both mechanical stress and chemically derived wall fragments generated by ISX, oryzalin, and COSs, supporting the emerging view that malectin-like RKs function as versatile cell-wall-integrity sensors ^73,110^.

Notably, neither cellulose perturbation nor COS treatment fully recapitulated ATHE-MIK2 interaction during infection, indicating the involvement of additional fungal signals. We excluded the MIK2-dependent SCOOP12 peptide as contributor and instead identified fungal RALFs (Fo-RALFs), but not endogenous ones, as key ATHE-dependent elicitors (Figs. 6 and S8). Other MIK2-dependent SCOOPs recently implicated in CWI sensing and immunity ^26^ may also function within the ATHE-MIK2 relationship during CWI responses, a possibility that remains to be tested together with their potential roles in plant defense against vascular pathogens. The absence of a role for ATHE in plant RALF-induced responses is consistent with its lack of interaction with FER, the main RK implicated in RALF perception (Figs. 4C and S3A). Fo-RALF exposure triggered the expression of genes associated with reinforcement of apoplastic barriers, previously shown to be induced during Fo infection ^29,93^. Meanwhile, *athe-1* roots failed to activate certain Fo-RALF-induced CW-reinforcement and apoplastic oxidative programs (Fig. 6E). MIK2 contributed partially to Fo-RALF responsiveness, and the *athe-1 mik2-1* double mutant phenocopied *mik2-1* (Fig. 6A). This genetic relationship indicates that MIK2 acts epistatically to ATHE within the same signaling pathway. Because diverse pathogens manipulate RALF signaling to suppress host immunity ^29–31^, ATHE may function as a *Brassicaceae*-specific surveillance receptor that detects pathogen interference with endogenous CWI signaling. ATHE and MIK2’s involvement in fungal RALF detection underscores the multifaceted nature of the ATHE-MIK2 module as an integrator of both pathogen-derived and CWI cues. RALFs have been implicated in immunity and CWI perception ^27,28^, and our findings extend this role to microbe-derived RALFs. Together, these results link microbial immunity responses to chemical and mechanical, but not osmotic, CWI alterations and identify Fo-RALFs as an additional class of ATHE-dependent signals modulating the ATHE-MIK2 module. Given that pectin methylation has been reported to be required for RALF perception via FER ^32,111^, these findings further raise the possibility that Fo-RALFs are detected through interactions with alternative CW components, with the ATHE–MIK2 module contributing to this recognition process, a possibility that requires further investigation.

Furthermore, despite ATHE being *Brassicaceae*-specific (Fig. S9), heterologous expression of Arabidopsis ATHE in tomato significantly improved resistance to Fol007 (Fig. 7). This demonstration of cross-family functionality suggests that the downstream pathways engaged by ATHE are conserved in non-*Brassicaceae* species. Notably, the recent identification of a MIK2-clade protein in tomato with properties similar to the Arabidopsis RK ^112^ supports the possibility of a conserved ATHE-MIK2 interdependency across plant lineages. ATHE therefore represents a promising candidate for engineering vascular pathogen resistance in major crops.

In summary, our study identifies ATHE/MEE39 as a previously uncharacterized receptor that integrates cellulose status, CW mechanical stress, and fungal peptide cues to activate protective responses during root infection. By forming an infection-reinforced complex with MIK2 and undergoing pathogen-triggered CME, ATHE connects early CWI perturbation to downstream immune output. These findings expand the CWI signaling network and provide new molecular tools for improving resistance to devastating vascular pathogens.

## ACKNOWLEDGMENTS

We are particularly grateful to Montserrat Capellade (CRAG) for her excellent technical assistance. We also extend our thanks to all members of the former Plant Cell Biology laboratory at ETH Zürich and the Cell Biology of Plant Resilience laboratory at CBGP for their technical support and stimulating scientific discussions. We thank Niko Geldner (University of Lausanne) for generously providing pUBQ10:NPSN12-YFP used in this study. We further acknowledge the Functional Genomics Center Zürich (FGCZ) for their support during the generation and analysis of the proteomic data and the ScopeM (ETH Zürich) and the CBGP for their confocal microscopy services.

## Funding

The work described in this manuscript was supported by the Swiss Federal Institute of Technology (ETH) Zurich, the Swiss National Science Foundation grant 310030_184769, the ETHZ Foundation grant 0-20172-16, and the MICIU/AEI/10.13039/501100011033 and European Union NextGenerationEU/PRTR grant CNS2023-144211 to C.S.-R.; the MCIN/AEI/ 10.13039/501100011033 grant PID2019-108595RB-I00, and the “Severo Ochoa Programme for Centres of Excellence in R&D” CEX2019-000902-S 10.13039/501100011033 to N.S.C.; the Netaji Subhas – Indian Council of Agricultural Research International Fellowship to A.K.; the European Research Council grant 773153, the University of Zurich, and the Swiss National Science Foundation grant 31003A_182625 to C.Z; the University of Lausanne, the European Research Council grant 716358, and the Swiss National Science Foundation grant 310030_204526 to J.S.

## Author contributions

C.S.R. conceived the study; J.C.M. and C.S.R. designed experiments; J.C.M., G.S.A., H.Y.H., M.M.D., F.R., L.C., F.M.G.-A., A.K., H.Y., and C.B. performed the experiments, J.C.M., H.Y.H, G.S.A., M.M.D., F.R., L.C., F.M.G.-A, A.K., H.Y., and J.S. analyzed the data; N.S.C., C.Z., J.S., and C.S.-R. supervised the work; J.C.M., M.M.D., and C.S.R. wrote the manuscript with comments from all authors.

## Declaration of Interests

The authors declare no competing interests.

## Data and materials availability

All data needed to evaluate the conclusions in the paper are present in the paper and/or the Supplementary Materials. The mass spectrometry proteomics data have been deposited to the ProteomeXchange Consortium via the PRIDE ^113^ partner repository with the dataset identifier PXD040278 and PXD040293.

## MATERIAL AND METHODS

### Plant material and growth conditions

*Arabidopsis thaliana* wild type (Col-0) and previously described mutants and marker lines in this background were used in this study: *ap2m-1* (SALK_083693) and *twd40-2-3* (SALK_112497) ^57–59^, *mik2-1* (SALK_061769) ^22^, *pTML:TML-YFP* ^43^, *pUBQ10:ARA7-YFP*, *pUBQ10:NPSN12-YFP* ^60^, *p35S:free-GFP* ^43^, *p35S:Lti6B-GFP* ^43,65^, *mik2-1 pMIK2:MIK2-mCherry* ^25^, YFP-CesA6 ^114^ and mCherry-TUA5 ^115^. *athe-1* (SALK_108641) T-DNA mutant was ordered through NASC. Transformation of *pATHE::ATHE-GFP* into *athe-1* mutants and *pATHE:mCherry-N7* into Col-0 plants were performed with floral dip ^116,117^. Transgenic lines were selected on soil by spraying with 0.1% BASTA for T1 generation. T2 heterozygous *pATHE::ATHE-GFP* and *pATHE::mCherry-N7* seedlings were screened for GFP and RFP fluorescence, respectively, to select for homozygous lines. *athe-1* and *mik2-1* mutants were crossed to generate the *athe-1 mik2-1* double mutant.

Arabidopsis was grown either vertically on 0.9% bactoagar supplemented with non-buffered half strength MS media (pH 5.7) or on soil (Substrat 2, Klasmann) if not specified. The growth conditions followed a 16-h light (24°C)/8-h dark cycle at 21°C. *Solanum lycopersicum* cv Marmander was also used in this work. Transformation of *p35S::ATHE-HA* into the WT was performed as described before ^118^. Transgenic lines were selected on MS media supplemented with kanamycin.

Tomato seeds were sterilized with 50% bleach for 15 min and 5 washes with sterile water. Then, they were placed in humid filter paper at 28°C in darkness for 3 days. Then, they were transferred to pots with coarse vermiculite and grown under 16-h light/8-h dark cycle and 28°C. After two weeks, each plant was watered weekly with 1 mL of non buffered ½ MS.

### Generation of constructs

All primers used to generate the constructs are listed in Table S3. To obtain *pATHE::ATHE-GFP,* 1773 bp upstream of the ATHE’s ORF (AT3G46330) were amplified using pNewEcoRI_pATHE_Fw and pNEW_pATHE_Rev primers; ATHE gDNA was cloned using gATHE_Fw and gATHE_Ala_Rev primers and the GFP was amplified preceded by a flexible alanine linker (GAAAAA) with the Ala_GFP_Fw and GFP_XbaI_Rev primers. *pATHE::mCherry-N7* was generated by amplifying ATHE promoter (1773pb) with the pNEW_EcoRI_pATHE_Fw and pATHE_mCh_Rev primers; mCherry, with mCh_Fw and mCh_Ala_Rev primers, and the N7 nuclear targeting sequence (AT4G19150) ^65^, preceded by a flexible alanine linker (GAAAAA), with Ala_Fw and N7_XbaI_Rev primers. In both cases, the amplified fragments were cloned by one-step ligation into a pNEW vector predigested with EcoRI and XbaI, and were assembled using NEBuilder® HiFi DNA Assembly Cloning Kit (BioConcept) following the manufacturer’s protocol. The final constructs were transformed into DH5ɑ *E. coli* cells, which were selected by kanamycin resistance. Electrocompetent GV3101 *Agrobacterium tumefaciens* co-transformed with the plasmids pSOUP ^119^ and that extracted from the *E.coli* indicated above. Resistant Agrobacterium cells selected with kanamycin, gentamicin and rifampin were used for floral dip to generate Arabidopsis transformants.

To obtain *p35S:ATHE-HA* for tomato transformation, the ORF of ATHE was cloned into the pART27 vector ^120^ followed at the 3’end by the HA epitope and under the control of the 35S promoter. The construct was transformed into DH5ɑ *E. coli* cells, which were selected by spectinomycin resistance. Finally, thermocompetent C58C1 *A. tumefaciens* were transformed with the construction and Agrobacterium cells selected with rifampicin, tetracycline and spectinomycin were used to transform tomato cv Marmande plants.

### Fungal material and growth conditions

*Fusarium oxysporum* 5176 (Fo5176) WT and the line expressing *pSIX1:GFP* were used for Arabidopsis infections. *Fusarium oxysporum* 007 (Fol007) WT was used for tomato infections. The cultivation and storage of the strains were performed as described earlier ^14,56,121^. In brief, spores germinated and grew in liquid potato dextrose broth (PDB) at 27°C in the dark for 5 days, and the culture was filtered through miracloth to harvest the spores.

Young Fo5176 hyphae were obtained for imaging ATHE-GFP and *pATHE:mCherry-N7* by confocal microscopy and determining ATHE-GFP protein levels using Western blot. 1 mL of 1 x 10^6^ spores were germinated in half MS (pH 5.7) + 1% sucrose liquid media by shaking overnight at 600 rpm in a 2 mL Eppendorf tube. The culture was centrifuged at 2,000 g for 5 min, the supernatant was discarded, and the pellet containing the young hyphae was washed 3 times with half MS liquid media to remove sucrose excess. These hyphae were then re-suspended to 10^7^ hyphae/mL with half MS.

### Fo5176 hydroponic infection assay

The hydroponic infection assay was performed as previously described ^17^. After 7 days of germination (dag) in 16-h light/8-h dark cycle 21°C, 80 rpm, the media was changed to ½ MS without sucrose where the seedlings grew for another 3 days. At 10 dag, the Fo-infected samples were exposed to a final concentration of 1 x 10^3^ Fo5176 spores / mL and shaken at 100 rpm for 30 min. The culture was changed to fresh ½ MS media and the original 80 rpm were restored. The mock-samples were treated in the same way but using ½ MS media.

### Fo5176 infection assays

Plate infections assays were performed as previously described ^14,56^. Briefly, 8 day-old seedlings germinated on top of Whatman filter paper strips in ½ MS media supplemented with 0.9% bactoagar were transferred to a mock or infection plate. The infection plates were prepared by spreading 100 µL 1 x 10^7^ *pSIX1:GFP* spores / mL on the media. Vascular penetration sites were recorded from 3 to 7 days post transfer (dpt), determined as strong and linear signals visible under a Leica M205 FCA fluorescent stereomicroscope equipped with a long pass GFP filter (ET GFP LP; Excitation nm: ET480/40x; Emission nm: ET510 LP). The primary root length was also measured in mock- and Fo-treated plants over time and the individual root length reduction was expressed as percentage from the average mock root length of the same genotype and time.

For Arabidopsis soil infection assays, 14-day-old plants grown on plates as described above, were immersed in either water (mock treatment) or a Fo5176 spore suspension (5 × 10⁶ spores/mL) with intermittent manual agitation for 15 minutes, then gently placed in soil pots containing a peat:vermiculite mixture (3:1). Pots were placed in a growth chamber with the same conditions as plants were previously grown but raising the temperature to 28°C. After 5 dpt, dead plants, whose root systems were mechanically damaged during the previous process were removed and not used for the analysis. Symptoms were recorded at 7, 10, 14, 17, and 21 dpt. All mock-treated plants (100%) across all genotypes remained asymptomatic.

### Fol007 soil infection assays

*S.lycopersicum* seeds were sterilized and grown in vermiculite as explained before. For infection experiments, roots were incubated in 20 mL water (Mock) or 20 mL of 5x10^6^ spores/mL (treatment) for 15 minutes. After the incubation, each plant was transferred into an individual vermiculite pot, to avoid competition between plants. The pots were covered with transparent film to maintain the humidity for 2 days. After 5 dpt, dead plants, whose root systems were mechanically damaged during the previous process were removed and not used for the analysis. Ten different plants of each genotype were used in the experiments. The mortality rates were recorded after 7, 10, 14, 17, 21 and 24 days. Plants that did not show any visible disease symptoms (asymptomatic) were removed from the experiment.

### Ralstonia solanacearum soil infection assay

Plants were grown on Jiffy pots with 8-h light/16-h dark cycles and 60% humidity at 22°C for 4-5 weeks. The infection assay of *Ralstonia solanacearum* bacteria was performed with the soil-drench method as previously described ^122^. In brief, 3 vertical holes were made in Jiffy pots, and the pots were submerged in a solution of overnight-grown *R. solanacearum* GMI1000 for 30 min. The solution was adjusted to an OD600 of 0.01 with distilled water, and 30 mL solution was used for each plant. Inoculated plants were then transferred to trays containing a thin layer of soil drenched with the same *R. solanacearum* solution, and the later growth condition followed 12-h light/12-h dark cycles and 60% humidity at 27°C during 22 days. Plant wilting symptoms were recorded and scored. The scoring measured symptoms on a 0 to 4 scale (0: no wilting, 1: 25% wilted leaves, 2: 50% wilted leaves, 3: 75% wilted leaves, and 4 = death).

### Subcellular fractionation and organelle proteome immunoisolation

Protocol was adapted from the one described in ^123^. Root tissue (≥500 mg per sample) from 3 dpt hydroponically infected or mock-treated plants (as described above) was ground with mortar and pestle on ice in ice-cold homogenization buffer (HB, 0.17 M 8% sucrose; 1 mM EDTA; 20 mM HEPES pH 7.5; 20 mM KCl; 1 mM DTT; 0.2% protease inhibitor cocktail (Sigma)) (5 mL/g root FW). Debris was pelleted by centrifugation at 2,500 rpm and 4 °C for 5 min, and the supernatant was further centrifuged at 4,500 rpm, 4 °C for 5 min. The following supernatant was again centrifuged at 2,500 rpm and 4 °C for 5 min to obtain the post-nuclear supernatant (PNS). Up to 4.5 mL of PNS was loaded on top of a 7 mL 42% sucrose cushion and ultra-centrifuged at 40,000 rpm and 4 °C for 30 min with an SW-40 rotor (Beckman). Membrane samples collected from the 8% / 42% sucrose interface were used for immunoisolation using µMACS GFP Isolation Kit (Miltenyi Biotec). One mL of each sample was incubated with 50 µL anti-GFP beads and mixed by inverting them every 20 min. Microcolumns were prepared by washing once with 200 µL elution buffer (50 mM Tris-HCl pH 8; 150 mM NaCl; 1% Triton X-100) and twice with 200 µL HB buffer. The samples were then loaded into the microcolumns, and washed 5 times with 200 µL HB buffer. 20 µL Laemmli buffer ^124^ preheated to 80 °C were added to each microcolumn and 5 min later another 80 µL preheated Laemmli buffer was added to the microcolumns. The eluate was collected and stored at -20 °C until further preparation for MS analysis.

### TCA precipitation, trypsin FASP digestion, and Stage Tip C18 clean-up

Proteins from pull-down samples (80 μL) were precipitated by addition of 9 μL ice-cold 100% trichloroacetic acid (TCA), followed by vortexing and incubation at 4°C for 30-45 min. Samples were centrifuged at 16,000 g and 4°C for 30 min, and the supernatant was discarded. Pellets were washed three times with 200 μL ice-cold acetone, with vortexing and centrifugation at 16,000 g and 4°C for 10 min each. The final pellet was air-dried and subjected to Filter-Aided Sample Preparation (FASP) digestion.

The FASP digestion was performed as previously described ^125^. 30 µL of SDS-lysis buffer (4%(w/v) SDS; 100 mM Tris-HCl pH 8.2; 0.1 M DTT) was added to resuspend the pellet, which was incubated at 800 rpm and 95 °C for 5 min, vortexed and spun down briefly. The mixture was then sonicated with 100% amplitude, 70% cycle for 1 min in a VialTweeter powered by UP200St ultrasonic processor (Hielscher Ultrasonics GmbH). Then, the sample was centrifuged at 16,000 g for 10 min, and 1 µL supernatant was taken to quantify protein amount with a Qubit (ThermoFisher), which was around 40 µg/sample. 200 µL UA buffer (8 M urea in 100 mM Tris-HCl pH 8.2) were added to the left 20 µL of sample and loaded to a filter unit, which was centrifuged at 14,000 g and RT for 25 min until the entire sample was loaded. Another 200 µL UA buffer was added to the filter unit and centrifuged at 14,000 g and RT for 20 min, and the flow-through was discarded. 100 µL IAA solution (0.05 M iodoacetamide in UA buffer) was added to the filter unit before mixing at 600 rpm on a thermoblock for 1 min. The sample was incubated for 5 min on the bench and centrifuged at 14,000 g and RT for 15 min. The filter unit was washed 3 times by adding 100 µL UA buffer and centrifuging at 14,000 g and RT for 12-15 min, followed by 2x washing by adding 100 µL 0.5 M NaCl and centrifuging at 14,000 g and RT for 12-15 min. 120 µL Trypsin (Promega) prepared in a 1:50 ratio in TEAB (0.05 M Triethylammoniumbicarbonate pH 8.5) was added to the filter unit that was then mixed at 600 rpm on a thermoblock for 1 min. The digestion was incubated on the bench overnight in a wet cell. After incubation, the filter unit was centrifuged at 14,000 g and RT for 15-20 min. 100 µL TEAB was added to the filter unit that was centrifuged again at 14,000 g and RT for 5 min. Finally, the eluate was acidified with 5% trifluoroacetic acid (TFA) to a final concentration of 0.5% TFA in the sample.

The peptide sample was further cleaned up with in-house made Stage Tip C18 ^126^ The Stage Tip was pre-equilibrated by loading 150 µL 100% acetonitrile (ACN) in a 2 mL tube and centrifuged at 2,000 g and RT for 1 min, and the flow-through was discarded. Next, the Stage Tip was equilibrated with 150 µL 60% ACN + 0.1% TFA and centrifuged at 2,000 g and RT for 1 min, and the flow-through was discarded. The Stage Tip was further conditioned by repeating twice of loading 150 µL 3% ACN + 0.1% TFA and centrifuging at 2,000 g and RT for 1 min. After conditioning, the trypsin-digested peptide sample was loaded into the Stage Tip, and the tip was centrifuged at 2,000 g and RT for 2 min. The loading was repeated until the entire sample was loaded. The Stage tip was then washed for 2 – 3 times by adding 150 µL 3% ACN + 0.1% TFA and centrifuging at 2,000 g and RT for 1 min. After the washing, the Stage Tip was placed in a 1.5 mL tube, loaded with 150 µL 60% ACN + 0.1% TFA, and centrifuged at 2,000 g and RT for 2 min. Finally, the eluted sample was completely dried by speed-vacuum and dissolved in 20 µL LC-MS (3% CAN; 0.1% FA) solution for MS analysis.

### LC-MS/MS analysis

Mass spectrometry (MS) analysis was performed with the support of Functional Genomics Center Zurich (FGCZ). Dissolved samples were injected by a Waters M-class UPLC system (Waters AG) operating in trap/elute mode. A Symmetry C18 trap column (5 µm, 180 µm X 20 mm, Waters AG) and a HSS T3 C18 reverse-phase column (1.8 µm, 75 µm X 250 mm, Waters AG) as separation columns were used. The columns were equilibrated with 99% solvent A (0.1% formic acid (FA) in water) and 1% solvent B (0.1% FA in ACN). Trapping of peptides was performed at 15 µl/min for 30 sec and afterwards the peptides were eluted using the gradient 1-40% B in 120 min. The flow rate was constant 0.3 µl/min and the temperature was controlled at 50°C. High accuracy mass spectra were acquired with an Q-Exactive HF mass spectrometer (Thermo Scientific) that was operated in data-dependent acquisition mode. A survey scan was followed by up to 12 MS2 scans. The survey scan was recorded using quadrupole transmission in the mass range of 350-1500 m/z with an AGC (Automatic Gain Control) target of 3E6, a resolution of 120’000 at 200 m/z, and a maximum injection time of 50 ms. All fragment mass spectra were recorded with a resolution of 30’000 at 200 m/z, an AGC target value of 1E5 and a maximum injection time of 50 ms. The normalized collision energy was set to 28%. Dynamic exclusion was activated and set to 30 s. All recorded data were automatically transferred to a data management system (B-Fabric) ^127^ and the peak lists were extracted automatically using FCC ^128^. Peak lists were searched on Mascot server v.2.4.1 (Matrix Science) against the Swiss-Prot (all species) database. Tryptic peptides with up to 1 possible miscleavages and charge states 2+, 3+, +4 were allowed in the search. Carbamidomethylated cysteine was specified as fixed modification, and oxidized methionine and acetylation at protein N-terminus were specified as variable modifications. Data were searched with a monoisotopic precursor and fragment ions mass tolerance of 10 ppm and 0.05 Da, respectively. The search results were then imported into Scaffold v.4 for visualization. A t-test analysis on MS2 spectra number was performed to compare the mock- and the Fo-treated samples of each marker line. We selected the entities with a minimum of 5 total MS2 spectra across the 4 replicates of each treatment, and a predicted PM localization (UniProt).

### Root growth inhibition assays

To test the sensibility to ISX, oryzalin, cellobiose, cellotriose or cellohexaose, plants were grown vertically on 0.9% bactoagar supplemented with ½ MS media and 1 % sucrose (pH 5.7) at 16-h light/8-h dark cycle at 21°C. At 5 dag, the seedlings were transferred to new ½ MS + 1% sucrose plates containing either 2 nM ISX, 0.1 µM oryzalin, 100 µM cellobiose/cellotriose/cellohexaose, using same amount of ethanol (ISX, oryzalin), or water (cellobiose, cellotriose, cellohexaose). At 5 dpt (ISX, oryzalin) or 3 dpt (cellobiose, cellotriose, cellohexaose), the primary root length was measured and the individual root length reduction was expressed as percentage from the average mock root length of the same genotype.

To test the sensibility to SCOOP12 peptides, plants were grown vertically on 0.9% bactoagar supplemented with ½ MS media and 1 % sucrose (pH 5.7) at 16-h light/8-h dark cycle at 21°C. At 3 dag, the seedlings were transferred to new ½ MS media plates containing 0, 10, 100, or 1000 nM SCOOP12 peptides. At 7 dpt, the primary root length was measured. The root length was normalized to the average mock root length within each genotype to show the root growth inhibition phenotypes. For the RALF root growth inhibition assay, synthetic RALFs (Table S4; from PhtdPeptides) were used. Col-0 and *athe-1* plants were grown on ½MS+S plates at 21 °C in the phytotron for five days before being transferred to 12-well plates for hydroponic growth containing the different RALFs at 2 µM final concentration. Three days later, seedlings were transferred to ½ MS + S plates, scanned, and their primary root lengths were quantified digitally using ImageJ.

### Confocal microscopy and data processing

Plasma membrane protein dynamics of the epidermal cells were imaged with a CSU-W1 Yokogawa spinning disk head fitted to a Nikon Eclipse Ti-E-inverted microscope with a CFI PlanApo × 100 N.A. 1.40 oil immersion objective, an EM-CCD ImageEM 1K (C9100-14) (Hamamatsu Photonics, Japan), and a ×1.2 lens between the spinning disk and camera. GFP was imaged using a 488 nm solid-state diode laser and a 525/50 nm emission filter; RFP was imaged with a 561 nm solid-state diode laser and a 609/54 nm emission filter. Time-lapse images were taken with 1 s intervals for 3 min if not specified. These images were processed and analyzed with Fiji ^129^. Backgrounds were subtracted by the “Subtract Background” tool (rolling ball radius, 20 pixels). To measure TML lifetimes, four frames were averaged by “Walking Average” and kymograph analysis was performed with the kymograph tool of FIESTA ^130^. Maximum projection from each image was created, and lines with line width = 2 pi were drawn along the discrete foci. The software later generated kymographs, and the kymographs were measured manually to report for the lifetimes of the foci. For live fungus treatment, young hyphae described above were used. For Fo5176 elicitor mix treatment, the full concentration (30 mg/mL) was used when no co-treatment was applied.

Endomembrane trafficking was analyzed in 5-day-old Col-0 wild type Arabidopsis seedlings grown on plates. They were incubated in ½ MS medium containing 50 μM brefeldin A (BFA) for 1 h and stained with 5 μM FM4-64 in the presence of 50 μM BFA on ice for 10 min. Subsequent mock or Fo5176 elicitor mix treatments were performed as described for spinning disk imaging and carried out on coverslips. Imaging was performed using a Zeiss LSM 780 or a Leica TCS SP8 STED/FLIM/FCS confocal laser scanning microscope, each equipped with a 63×/1.4 NA oil immersion objective. GFP was excited at 488 nm and detected between 500-545 nm, and RFP was excited at 561 nm and detected between 592-754 nm. For quantification of endocytosis, epidermal cells with uniform plasma membrane signal were selected and analyzed in Fiji. Plasma membrane fluorescence was measured along a line selection (3-pixel width) generated from a polygonal cell outline. The corresponding cytoplasmic signal was measured after mask generation and three iterations of erosion to exclude the plasma membrane. Plasma membrane-to-cytoplasm fluorescence intensity ratios were calculated, and cell outlines and areas were stored for further analysis. For BFA body quantification, the Z-plane containing the highest number of BFA bodies was selected from Z-stacks, and BFA bodies were counted per cell.

To test the response of ATHE-GFP, MIK2-mCherry, NPSN12-YFP, ATHE promoter activity, and ATHE localization in different root layers, 8-day-old roots were treated with Fo5176 WT young hyphae (prepared as described above) on ½ MS bactoagar plates or with 300 nM ISX (Isoxaben Sigma 36138-100MG dissolved in EtOH), 1 µM oryzalin (Fluka 36182-100MG dissolved in EtOH), 300 mM sorbitol (D-Sorbitol Roth 6213.1-1KG; dissolved in water), 100 µM cellobiose (D-(+)-Cellobiose Sigma 22150-10G; dissolved in water), 100 µM cellotriose (Cellotriose Megazyme O-CTR-50MG; dissolved in water),100 µM cellohexaose (Cellohexaose Megazyme O-CHE-10MG; dissolved in water), or 2 µM Fo-RALF in ½ MS liquid media supplemented with 1 % sucrose. Media containing the same volume concentration of ethanol or water were used as the mock treatment.

Root epidermal cells expressing *pATHE:ATHE-GFP, pMIK2:MIK2-mCherry* or *pATHE:ATHE-GFP pMIK2:MIK2-mCherry* were imaged with a CSU-W1 Yokogawa spinning disk head fitted to a Nikon Eclipse Ti-E-inverted microscope with a CFI PlanApo × 100 N.A. 1.40 oil immersion objective, two iXon Ultra EM-CCD cameras (Andor, GB), and a ×1.2 lens between the spinning disk and camera were used. GFP was detected using a 488 nm solid-state diode laser and a 525/50 nm emission filter, mCherry was detected with a 561 nm solid-state diode laser and a 630/75 nm emission filter. Time-lapse images were taken with 1 s intervals for 3 min if not specified. Z-stack analysis was performed in 30 x 0.25 μm sections. The images were processed and analyzed with Fiji ^129^. Backgrounds were subtracted by the “Subtract Background” tool (rolling ball radius, 20 pixels). For colocalization analysis, only images focused at the PM focal plane of the cell were used. The R’s Pearson value was calculated using the “Coloc 2” plug-in of Fiji.

Root epidermal cells expressing *pATHE:mCherry-N7* and root cells expressing ATHE-GFP (for ATHE localization in different root layers) were imaged with a Zeiss 780 or Zeiss 880 confocal laser scanning microscope equipped. mCherry was visualized under a 20x 0.5 NA objective. RFP was visualized using 561 nm laser excitation and 592-754 nm spectral detection. Z-stack images were taken, and image tiling and stitching was performed with the ZEN software (Zeiss). The images were further processed and analyzed with Fiji. Frames that had the strongest signal of each cell were selected from the Z-stack and the mean gray values were measured with the polygon tool. For ATHE localization in different root layers, images were acquired using a 40x water immersion objective and acquisition settings included a digital gain of 800 for GFP and 700 for Propidium Iodide (PI), scan speed of 5, line averaging of 2-4, bidirectional scanning in the X-direction enabled, and a pinhole size set to 1 Airy unit. Fluorescence signals from reporter genes and stains were visualized using the following excitation/spectral detection settings: GFP, 488 nm/495-537 nm and PI, 561 nm/592-660 nm. Orthogonal views were generated using Fiji ^129^.

### Membrane protein isolation and Western blot analysis

To analyze ATHE-GFP protein levels, membrane proteins were extracted by adapting from a previous method ^131^. For each experiment, 8-day-old roots were harvested and ground in liquid nitrogen. To homogenize the samples, 20 mg root tissue was added with 120 µL extraction buffer 1 (50 mM Tris-HCl pH 7.6, 150 mM NaCl, Protease Inhibitor Cocktail pill (PiC 10mg tablet - cOmplete, Roche), 0.5 mM EDTA) on ice. The samples were then centrifuged at 16,250 g and 4°C for 15 min, and the supernatant containing the cytosolic proteins was removed. The pellet, which contained plant debris, unbroken cells, cell wall components, orgenelles, nuclei, and microsomal fractions, was re-suspended with 60 µL extraction buffer 2 (50 mM Tris-HCl pH 7.6, 150 mM NaCl, Protease Inhibitor Cocktail pill, 0.5 mM EDTA, 0.5% Triton-X, 0.5% NP-40) without pipetting. The re-suspended samples were centrifuged at 12,000 g and 4°C for 20 min. The supernatant was collected and quantified for protein concentration with Quick Start Bradford protein assay following the manufacturer’s protocol. The samples were then heated in 1x Laemmli buffer at 95 °C for 10 min. SDS-PAGE was performed by loading the same amount of protein (30 μg/sample) in precast SDS 4-12% acrylamide gradient gels (Expedeon, GB). Protein transfer to nitrocellulose membranes was performed in wet conditions (Bio-Rad system). For western blot analysis, nitrocellulose membranes were blocked with 3% non-fat dry milk / 0.1 % Tween-20, incubated with the primary antibodies at RT for 2h (or overnight at 4 °C), washed, and incubated with secondary antibodies at RT for 1h. Primary antibodies were anti-GFP (3H9, 1:1000 dilution, Chromotek) and anti-SYP21 (1:1000 dilution; ^66^), which were recognized by the respective secondary antibodies, goat anti-rat HRP (1:5000 dilution; Axon Lab) and goat anti-rabbit HRP (1:5000 dilution; Axon Lab), respectively. Protein detection was performed with the Westar Supernova kit (Cyanagen), which provides sensitive chemiluminescence detection. After the analysis with the antibodies, the amount of transferred protein was evaluated by incubating the membrane with Ponceau S staining solution (0.1% (w/v) Ponceau S in 5% (v/v) acetic acid) for 10 min and washing with distilled water to remove the background.

### Co-immunoprecipitation analysis

To detect the possible interactors of ATHE, a co-immunoprecipitation (co-IP) targeting ATHE-GFP analysis was performed including Lti6B-GFP co-IP as a negative control. 500 mg root tissue from Fo- or mock-hydroponically treated seedlings (as described above) at 3 dpt was homogenized with a mortar and a pestle on ice in 1 ml extraction buffer (50 mM Tris-HCl pH 8.0, 150 mM NaCl, 1% Triton-X, 1% PVPP, Protease Inhibitor Cocktail pill (PiC 10mg tablet - cOmplete, Roche), 1mM DTT). The samples were centrifuged at 5,000 rpm and 4°C for 5 min, and the supernatant was transferred to a new tube where they were centrifuged again at 5,000 rpm and 4°C for 20 min, to obtain the membrane enriched fractions. The supernatant was transferred to a new tube containing 50 μL of anti-GFP μMACS beads (Miltenyi Biotec) and incubated at 4°C for 1 hr with rotation. Next, anti-GFP μMACS beads were processed using the μMACS column purification protocol and washed with the wash buffer (50 mM Tris-HCl pH 8.0, 300 mM NaCl, 1% Triton-X, Protease Inhibitor Cocktail pill, 1 mM DTT). Final elution was performed using 80 μl of the elution buffer (50 mM Tris-HCl pH 7.5, 2% SDS, 1 mM DTT) at 80°C. After elution, samples were stored at -20°C before being analyzed with MS.

The isolated proteins were subjected to TCA precipitation, trypsin FASP digestion, Stage Tip C18 clean-up, and LC-MS/MS analysis as described above. MS spectra were further analyzed using MaxQuant v.1.1.1.14 software with a peptide FDR of 1% with LFQ intensity quantification ^132^. The TAIR 10 (www.arabidopsis.org) and SUBA 5 (www.suba.live) databases were used for peptide identification and cellular localization annotation, respectively. LFQ intensities were then analyzed using Perseus software v.1.6.5.0 ^133^.

Protein extracts from membrane enriched fractions were also used for co-immunoprecipitation experiments using GFP-trap magnetic beads (Chromotek, http://www.chromotek.com/), and analyzed by SDS-PAGE and western blotting. Antibodies against GFP (3H9, 1:1000 dilution, rat, Chromotek) and RFP (6G6, 1:1000 dilution, mouse, Chromotek) were used for detecting ATHE-GFP and MIK2-mCherry, respectively. As secondary antibodies, goat anti-rat HRP (1:5000 dilution; Axon Lab) and goat anti-mouse HRP (1:5000 dilution; Axon Lab) were used, respectively. The protein detection was performed with the Westar Supernova kit (Cyanagen).

### RNA extraction and qRT-PCR analysis

100 mg root material was harvested and frozen immediately with liquid nitrogen. RNA was extracted with the RNeasy Plant Mini Kit (Qiagen) following the manufacturer’s protocol. Later, cDNA was synthesized using the Thermo Scientific MaximaTM H Minus cDNA Synthesis Master Mix with dsDNase (ThermoFisher) following the manufacturer’s protocol. The qRT-PCR was performed under the following PCR conditions: 95°C for 3 min, 40 cycles of 94°C for 10 s, 58°C for 15 s, and 72°C for 10 s using Fast SYBR Green Master Mix (ThermoFisher) in a 10 µL reaction. The reference gene GAPDH 600b (^134^ was amplified in parallel on each plate for normalization. The primers used to amplify target genes are listed in Table EV3. The 2 ΔCT method was used to quantify the relative expression of each gene ^135^.

### Protein expression and purification

*Spodoptera frugiperda* codon-optimized synthetic genes (Invitrogen GeneArt), coding for *Arabidopsis thaliana* ATHE (residues 24-514; AT3G46330) was cloned into a modified pFastBAC vector (Geneva Biotech) providing a 30K signal peptide ^136^ and a C-terminal TEV (tobacco etch virus protease) cleavage site, and a StrepII-9xHis affinity tag. Baculovirus generation was carried out using DH10 cells and virus production and amplification was done in Sf9 cells. Trichoplusia ni Tnao38 cells ^137^ were used for protein expression, infected with ATHE virus with a multiplicity of infection (MOI) of 3 and incubated 1 day at 28°C and 2 days at 22°C at 110 r.p.m. The secreted protein was purified on a Ni^2+^ (HisTrap excel, Cytiva, equilibrated in 25 mM KP_i_ pH 7.8 and 500 mM NaCl) followed by Strep (Strep-Tactin Superflow high-capacity, IBA Lifesciences, equilibrated in 25 mM Tris pH 8.0, 250 mM NaCl, 1 mM EDTA) affinity chromatography. The protein was incubated with TEV protease to remove the tags. The protein was further purified by SEC on a Superdex 200 Increase 10/300 GL column (Cytiva) equilibrated in 20 mM citrate pH5.0, 150mM NaCl. The protein was concentrated using Amicon Ultra concentrators (Millipore, molecular weight cut-off 30,000), and SDS-PAGE was used to assess its purity and structural integrity.

### Isothermal titration calorimetry (ITC)

Experiments were performed at 25 °C using a MicroCal PEAQ-ITC (Malvern Instruments) with a 200 µL standard cell and a 40 μL titration syringe. The protein was dialyzed against the corresponding ITC buffer (20 mM citrate pH5.0, 100 mM NaCl) and the ligands dissolved in the same buffer. The ATHE vs. cellobiose (Merk, C7252, purity > 99%) and cellotriose (Megazyme, O-CTR-50mg; purity > 95%) oligo experiments were performed using a 30 µM protein solution in the cell and 500 µM of the ligand in the syringe. A typical experiment consisted of injecting 3 µl of ligand solution into the cell at 150 s intervals and 500 r.p.m stirring speed. ITC data were corrected for the heat of dilution by subtracting the mixing enthalpies for titrant solution injections into protein free ITC buffer. Experiments were done in duplicates, and data were analyzed using the MicroCal PEAQ-ITC Analysis Software provided by the manufacturer.

### Quantification of CSC density and MT coverage at cortex

Both methods were described earlier ^14,138^. In brief, Gaussian kernel (sigma = 1.33 µm) and Otsu algorithm ^139^ was applied to every picture to detect cell boundaries. To detect particles (to quantify CSC density), a Laplacian image was generated from pictures that were smoothed using FeatureJ (Erik Meijering, Biomedical Imaging Group, EPFL Lausanne) with a Gaussian kernel with sigma = 0.2 µm. “Find Maxima” command was used to detect peaks with a noise tolerance of 800. Wider smoothing kernel, Sigma = 0.8 µm, and noise tolerance of 120 was employed to detect Golgi particles that were subtracted from the total number of particles. The CSC density was estimated by dividing the particle count by the cell area. For the microtubule coverage, various thresholds, based on signal to noise ratio of each individual picture, were applied to detect microtubules with low pixel noise. The microtubule signal and total cell area ratio was used to calculate microtubule cell coverage. All image analysis was performed using Fiji software.

### Phylogenetic analysis

The phylogeny was inferred using the Maximum likelihood method and ^140^ model of amino acid substitutions and the tree with the highest log likelihood (-4.737.34) is shown. The percentage of replicate trees in which the associated taxa clustered together (1000 replicates) is shown next to the branches ^141^. The initial tree for the heuristic search was selected by choosing the tree with the superior log-likelihood between a Neighbor-Joining (NJ) tree ^142^ and a Maximum Parsimony (MP) tree. The NJ tree was generated using a matrix of pairwise distances computed using the ^140^ model. The MP tree had the shortest length among 10 MP tree searches each performed with a randomly generated starting tree. The evolutionary rate differences among sites were modeled using a discrete Gamma distribution across 5 categories (+G, parameter = 1.4703). The analytical procedure encompassed 43 amino acid sequences. The partial deletion option was applied to eliminate all positions with less than 95% site coverage resulting in a final data set comprising 142 positions. Evolutionary analyses were conducted in Mega 12 ^143^ utilizing up to 7 parallel computing threads.

## Notes

### Competing Interest Statement

The authors have declared no competing interest.

