## Supplementary material for "The Receptor Kinase MEE39/ATHE Mediates Cell Wall Integrity Surveillance During Root Vascular Pathogen Infection": Suppl Figs and Tables

Juan Carlos Montesinos *et al.*

### SUPPLEMENTARY FIGURES AND TABLES

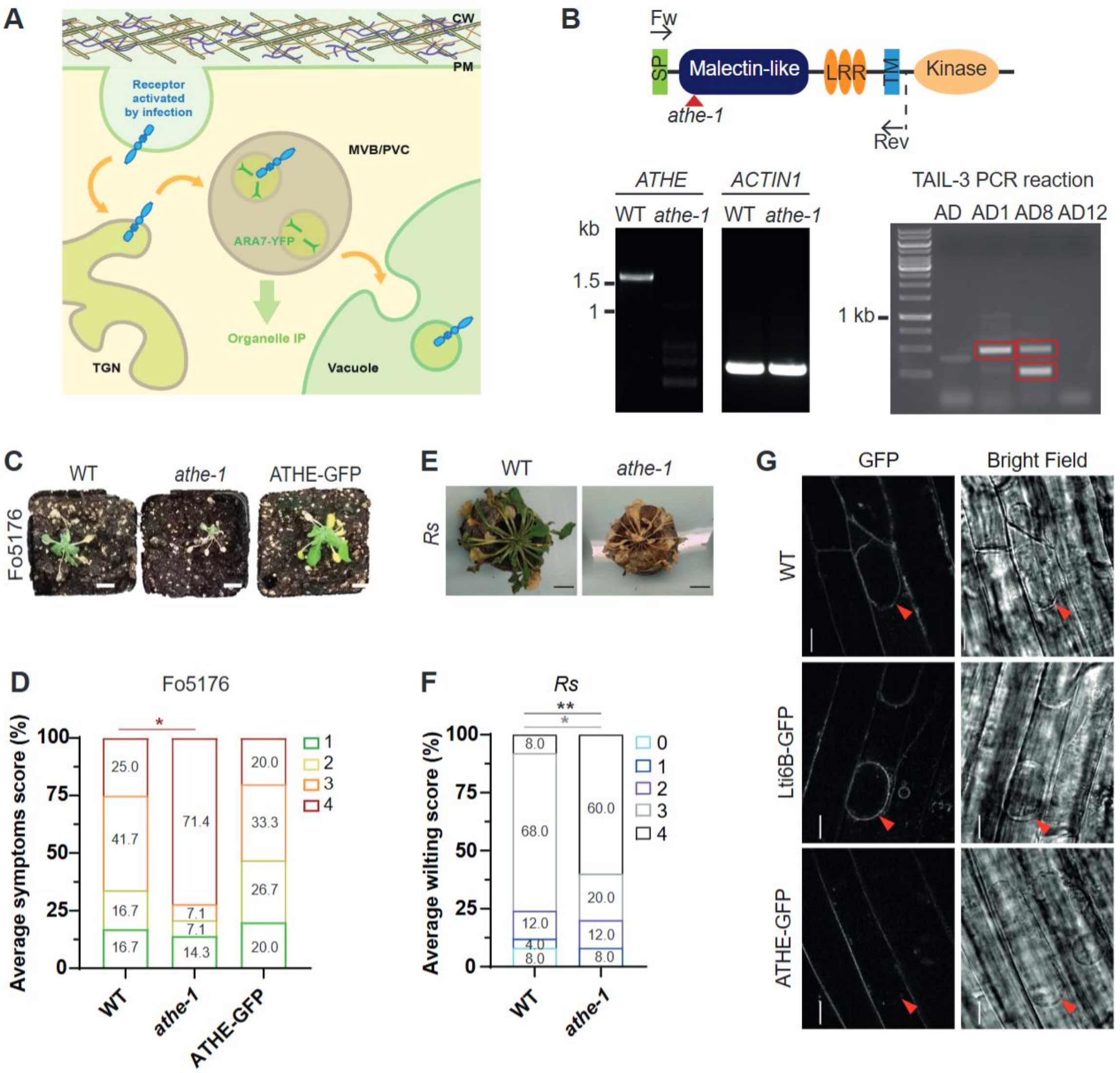

**Figure S1. ATHE is a plasma membrane-localized receptor required for plant defense against vascular pathogens. A.** Illustration of the strategy to identify new plant receptors involved in defense to Fo5176. After ligand perception, plasma membrane (PM) receptors are internalized by endocytosis and transported through the trans-Golgi network (TGN) and the Multivesicular body/Prevacuolar compartment (MVB/PVC) for vacuolar degradation. PM proteins involved in Arabidopsis-Fo interaction should be enriched in the MVB/PVC when plants are infected by the fungus.

The MVB/PVC marker line ARA7-YFP was utilized to immunoprecipitate proteins located in the MVB/PVC. The orange arrows indicate the canonical endocytic pathway towards the vacuole. **B.** Upper panel: localization of the T-DNA insertion in *athe-1* (red arrowhead) and primer pair used to detect *ATHE* by RT-PCR (black arrows). Left bottom panel: *ATHE* and *ACTIN1* RT-PCR products in 14-day-old WT (Col-0) and *athe-1* seedlings. Three independent replicates were performed with the same results. Right bottom panel: TAIL-3 PCR reaction to identify the number of T-DNA insertions in *athe-1* using 4 different AD primer combinations flanking regions of *ATHE*/*MEE39* gene (AD, AD1, AD8, and AD12). The bands marked with red rectangles were purified and sequenced. **C.** Representative images of WT, *athe-1,* and ATHE-GFP plants at 14 days post-treatment (dpt) with Fo5176 spores. **D.** Scoring of Fo5176-induced disease symptoms in soil-infected plants as described in (C). Phenotypes: 1 (asymptomatic plants), 2 (plants with ≤ 50% of yellow or dry leaves ), 3 (plants with > 50% of yellow or dry leaves), or 4 (dead plants). Data represent average in percentage (%); *N* = 15 plants/genotype from 1 experiment out of 3 with similar results. Repeated measures (RM) $Chi^2$ test, *P ≤ 0.05. The colors of the asterisks correspond to the colors assigned to the different symptom phenotypes **E.** Representative images WT and *athe-1* plants at 18 dpt with *Ralstonia solanacearum* (Rs) . Scale bars = 1 cm. **F.** Scoring of Rs-induced wilting symptoms in soil-infected plants at 18dpt as shown in (E). Wilting score 0: asymptomatic, 1: 25% wilted leaves, 2: 50% wilted leaves, 3: 75% wilted leaves, 4: 100% wilted plants. Data represent average in percentage (%); *N* = 25 plants/genotype from 1 experiment out of 3 with similar results. Repeated measures (RM) $Chi^2$ test, *P ≤ 0.05; **P ≤ 0.005. The colors of the asterisks correspond to the colors assigned to the different wilting scores. **G.** Representative spinning disk confocal images of ATHE-GFP or Lti6B-GFP in 10-days old root epidermal cells of MD plasmolyzed with 0.8 M mannitol for 10 min. WT (Col-0) roots were included as control for cell wall autofluorescence. Red arrowheads indicate PM detached from the cell wall. Scale bar = 10 μm.

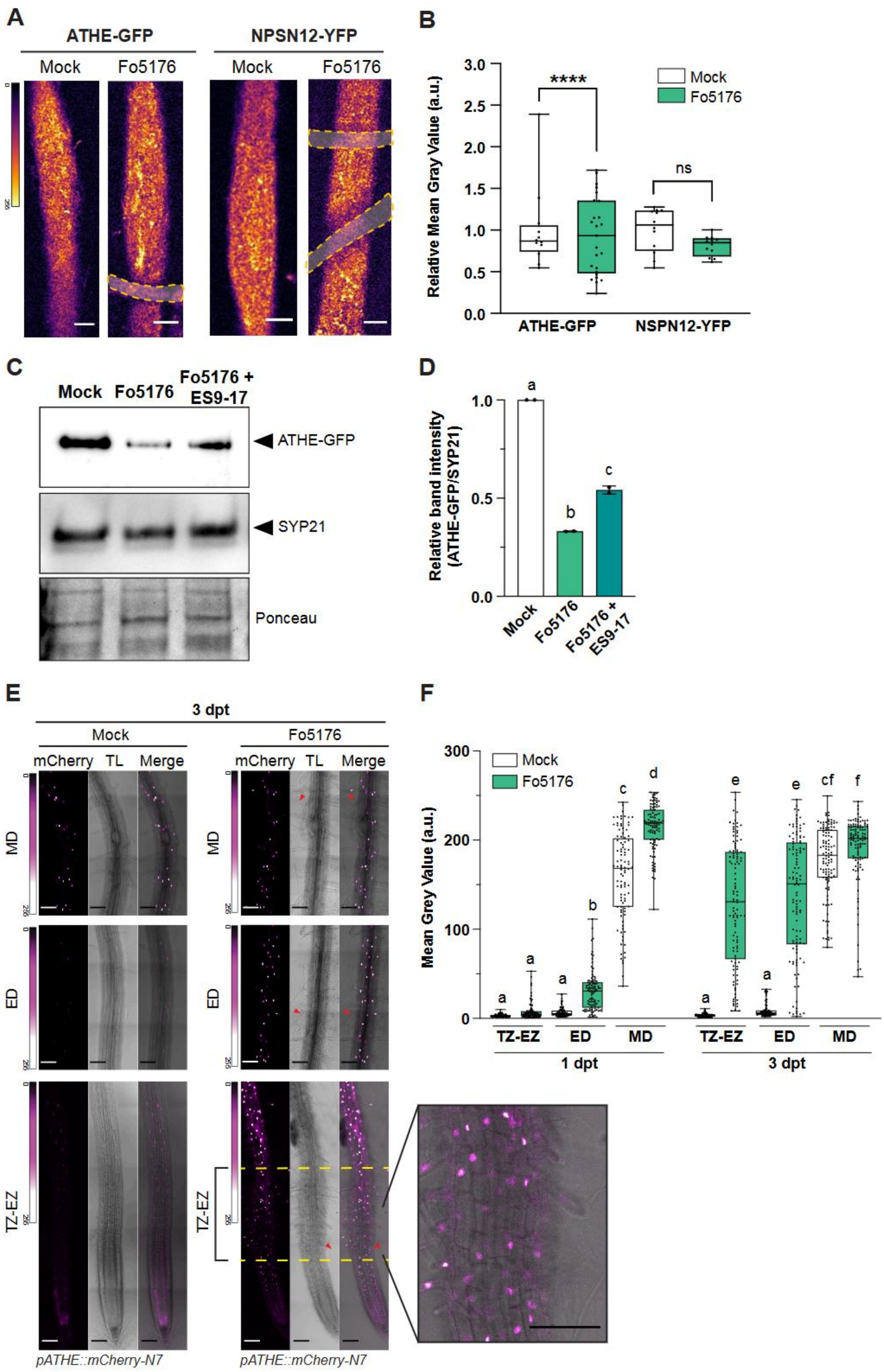

**Figure S2. ATHE expression under stresses, internalization and gene expression are dependent on the Fo5176 root kinetics colonization. A.** Representative spinning disk confocal images of ATHE-GFP or NPSN12-YFP in 8-

day-old root epidermal cells in the MD after 15-30 minutes exposure to Fo5176 young hyphae (yellow). The scale on the right represents the fluorescence signal intensity of the tagged proteins. Scale bars = 5 µm. **B.** Amount of ATHE-GFP and NPSN12-YFP at the plasma membranes of cells as described in (A), quantified as mean gray value (arbitrary units, a.u.) of fungal-exposed samples normalized to their corresponding Mock. . Areas-of-interest were selected manually to avoid the hyphae. Box plots: centerlines show the medians; box limits indicate the 25th and 75th percentiles; whiskers extend to the minimum and maximum. $N \geq 12$ cells (2-4 cells/root and ≥ 3 roots/experiments were imaged). Repeated measures (RM) *t*-test. ns, not significant. **C.**Representative Western blot of total ATHE-GFP and SYP21 proteins in 8-day-old ATHE-GFP roots at 1 dpt with Mock, Fo5176, or Fo5176 + 10 µM ES9-17. Ponceau S staining provides loading control. **D.** ATHE-GFP signal quantification from immunoblots as in (C) .The intensity of ATHE-GFP band in every condition was normalized to the intensity of the SYP21 band in the same condition. Av ± SEM, $N = 2$ independent experiments. RM one-way ANOVA with Tukey's multiple comparisons test. Alphabet letters indicate significant differences with $P \leq 0.05$. **E.** Representative confocal images of mCherry-N7 in *pATHE:mCherry-N7* 8-day-old root epidermal cells at the root differentiated zones indicated in Fig 2(D) at 3 days post treatment (dpt) with Fo5176 young hyphae (indicated by red arrowheads). The scale on the left of each image represents the fluorescence signal intensity of mCherry. TL, transmitted light. . Scale bars = 100 µm. **F.** Amount of mCherry signal intensity as depicted in (E) at 1dpt (left) and 3 dpt (right) with Fo5176 young hyphae, quantified as mean gray value (arbitrary units, a.u.). Box plots as described in (B). $N \geq 100$ cells from 3 experiments (≥ 6 cells/root and 3 roots/experiment were imaged). RM one-way ANOVA with Kruskal-Wallis multiple comparison test. Alphabet letters indicate significant differences with $P \leq 0.05$.

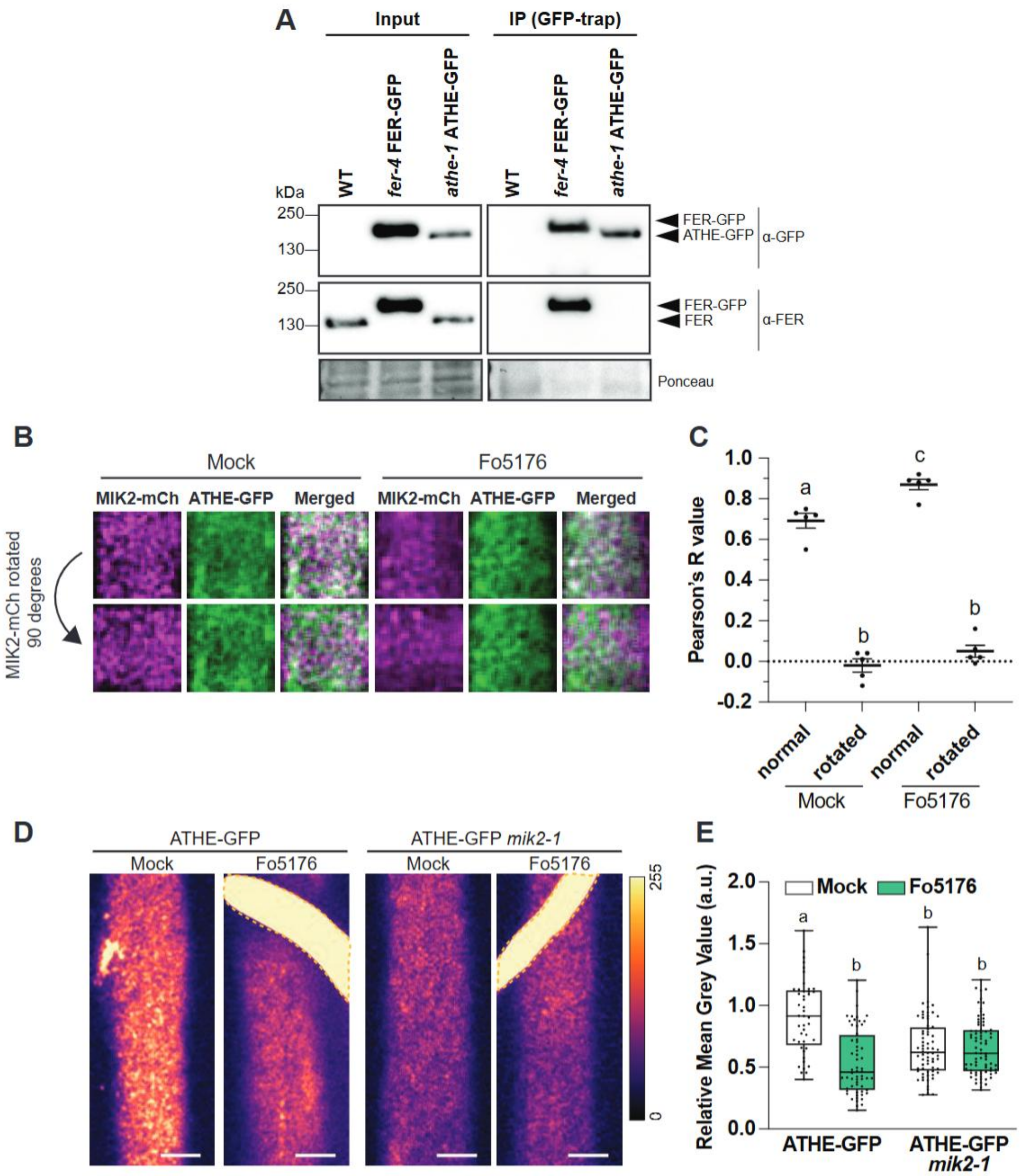

**Figure S3. Control for the MIK2-ATHE colocalization and ATHE-FER co-immunoprecipitation**. **A.** Representative Western blot co-immunoprecipitation analysis of ATHE-GFP and FERONIA (FER) using GFP-trap beads, in 8-day-old WT, *fer-4* FER-GFP and *athe-1* ATHE-GFP roots, using anti-GFP antibody (ATHE-GFP/FER-GFP) and a specific anti-FER antibody (FER and FER-GFP). Ponceau S staining provides loading control. **B.** Colocalization of ATHE-GFP (green) and MIK2-mCherry (magenta) as depicted in Fig 4(D). **C.** Pearson's R value was quantified in 5 of the images used for the Fig 4(D-E), raw (normal) and rotating 90 degrees the picture of the mCherry channel (rotated). Box plots: centerlines show the medians; box limits indicate the 25th and 75th percentiles; whiskers extend to the minimum and maximum.

Repeated measures (RM) one-way ANOVA with Tukey's multiple comparisons test. Alphabet letters indicate significant differences with $P \leq 0.05$. **D.** Representative spinning disk confocal images of ATHE-GFP in *athe-1* (ATHE-GFP) or *athe-1 mik2-1* (ATHE-GFP *mik2-1*) background in 8-day-old root epidermal cells in the MD after 1-day of exposure to Mock or Fo5176 young hyphae (yellow). The scale on the right represents the fluorescence signal intensity. Scale bars = 5 µm. **E.** Amount of ATHE-GFP at the plasma membranes of cells as described in (D), quantified as mean gray value (arbitrary units, a.u.) of fungal-exposed samples normalized to ATHE-GFP Mock. Box plots as described in (C). $N \geq 48$ cells per root area/treatment were quantified from 3 experiments (2-4 cells/root and ≥ 3 roots/experiments). RM one-way ANOVA with Tukey's multiple comparisons test. Alphabet letters indicate significant differences with $P \leq 0.05$.

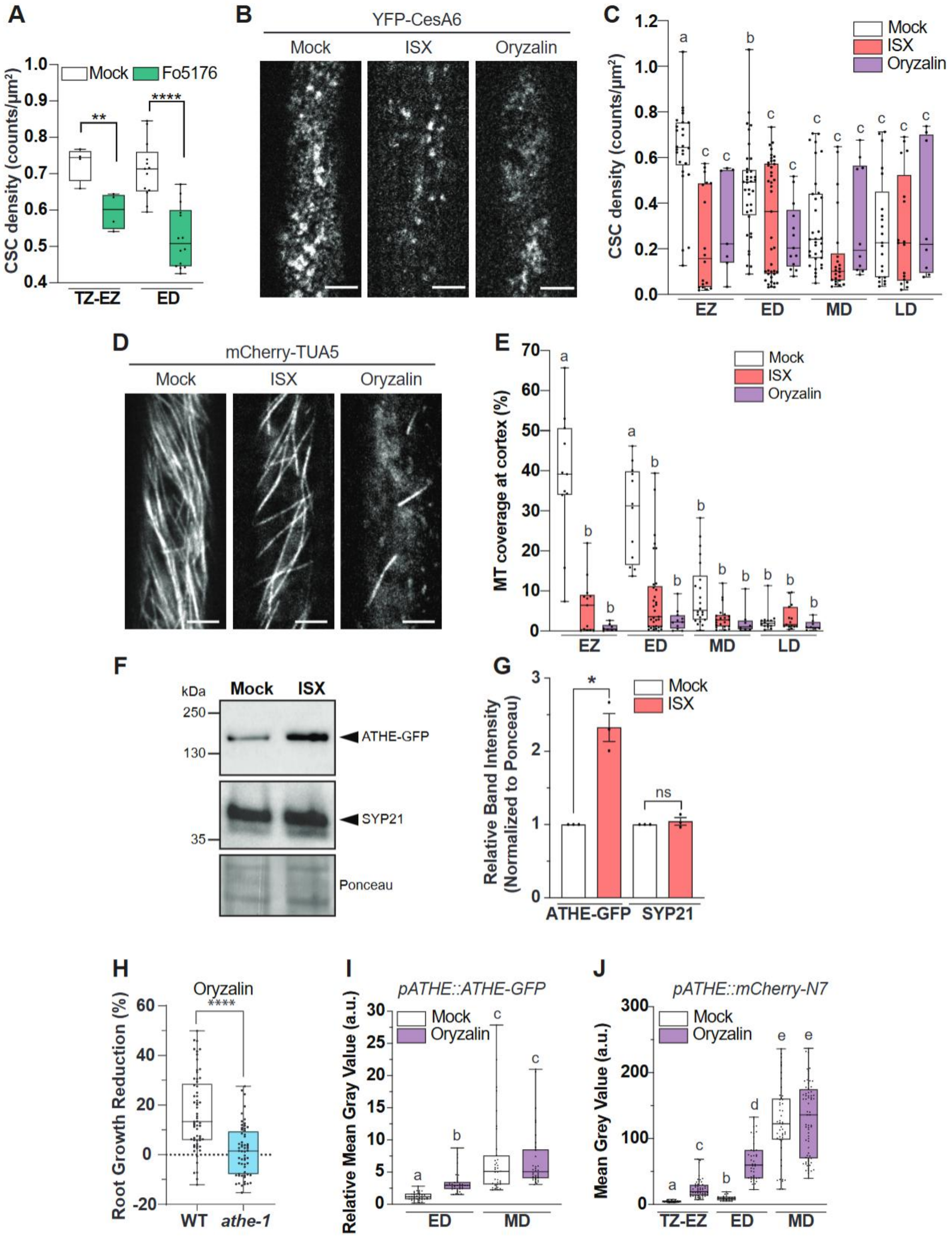

**Figure S4. Isoxaben induces disruption of cellulose synthesis complexes and microtubules depolymerization in early root developmental areas that the microtubule depolymerizing drug oryzalin mimics. A.** Density of cellulose synthesis complexes (CSC) based on YFP-CesA6 signal at the PM of TZ-EZ and ED epidermal cells after 1-day of exposure to Fo5176 young hyphae. Box plots:

centerlines show the medians; box limits indicate the 25th and 75th percentiles; whiskers extend to the minimum and maximum. *N* ≥ 4 cells/root using ≥ 6 roots/condition. Repeated measures (RM) *t*-test. **P ≤ 0.01, ****P ≤ 0.0001. **B.** Representative spinning disk confocal images of YFP-CesA6 in 8-day-old root epidermal cells of ED after 1 day exposure to Mock, 300 nM isoxaben (ISX) or 1 µM oryzalin. Scale bars = 5 µm. **C.** Density of cellulose synthesis complexes (CSC) based on YFP-CesA6 signal as depicted in (B) at the root differentiated zones indicated in Fig 2(D). Box plots as described in (A). *N* ≥ 14 cells from 3 experiments (2-4 cells/root and ≥ 3 roots/experiments). RM one-way ANOVA with Tukey's multiple comparisons test. Alphabet letters indicate significant differences with P ≤ 0.05. **D.** Representative spinning disk confocal images of mCherry-TUA5 in 8-day-old root epidermal cells of ED after 1 day exposure to Mock, 300 nM ISX or 1 µM oryzalin. Scale bars = 5 µm. **E.** Microtubules (MT) coverage based on mCherry-TUA5 signal in epidermal cells as depicted in (D) at the root differentiated zones indicated in Fig 2(D). Box plots as described in (A). *N* ≥ 12 cells from 3 experiments (2-4 cells/root and ≥ 3 roots/experiments). RM one-way ANOVA with Tukey's multiple comparisons test. Alphabet letters indicate significant differences with P ≤ 0.05. **F.** Representative Western blot of total ATHE-GFP and SYP21 proteins in 8-day-old ATHE-GFP roots treated either with Mock or 300 nM ISX for 1 day. Ponceau S staining provides loading control**. G.** ATHE-GFP and SYP21 signal quantification from immunoblots as in (F). For each protein, the intensity of the ISX-treated sample was normalized to that of the Mock. Av ± SEM, *N* = 3 experiments. RM *t*-test. *P ≤ 0.05. ns, not significant. **H.** Root growth reduction of 0.1 µM oryzalin-treated wild-type (WT; Col-0) and *athe-1* seedlings relative to Mock-treated ones, after 5 days post treatment (dpt). Box plots as described in (A). *N* ≥ 60 roots were from 4 experiments (≥ 15 roots/experiments). RM *t*-test. ****P ≤ 0.0001. **I.** Amount of ATHE-GFP at the plasma membranes of epidermal cells of the ED and MD in response to Oryzalin 1 µM or Mock treatment for 1 day, quantified as mean gray value (arbitrary units; a.u.). Box plots as described in (A). *N* ≥ 34 cells (3-5 cells/root) per root area/treatment. RM One-way ANOVA with Kruskal-Wallis multiple comparison test. Alphabet letters indicate significant differences with P ≤ 0.05 **J.** Amount of the mCherry signal intensity of *pATHE::mCherry-N7* epidermal cells at the root differentiated zones indicated in Fig 2(D) in response to Oryzalin 1 µM or Mock treatment for 1 day, quantified as mean gray value (arbitrary units; a.u.). Box plots as described in (A). *N* ≥ 21 cells (8-15 cells/root) per root area/treatment. RM one-way ANOVA with Kruskal-Wallis multiple comparison test. Alphabet letters indicate significant differences with $P \leq 0.05$.

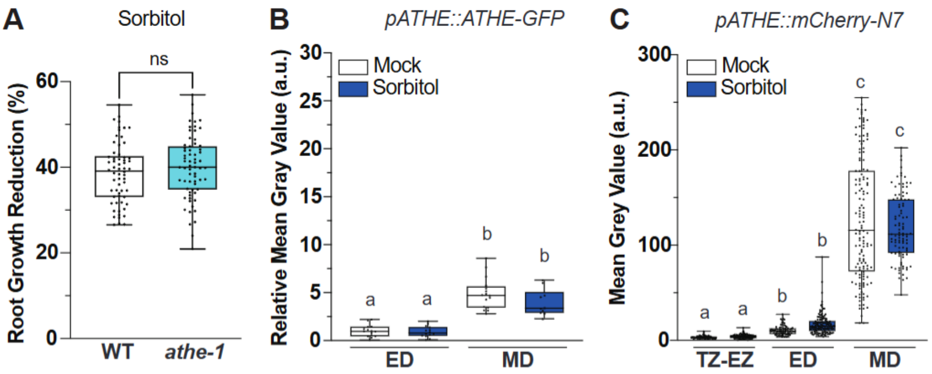

**Figure S5. Root growth rate, the differential regulation of ATHE protein level and gene expression is not altered by osmotic stress. A.** Root growth reduction of Mock or 200 mM Sorbitol-treated wild-type (WT; Col-0) and *athe-1* seedlings relative to Mock-treated ones, after 5 days post treatments (dpt). Box plots: centerlines show the medians; box limits indicate the 25th and 75th percentiles; whiskers extend to the minimum and maximum. $N \geq 62$ roots from 4 experiments (≥ 12 roots/experiments). Repeated measures (RM) *t*-test. ns, not significant. **B.** Amount of ATHE-GFP at the plasma membranes of epidermal cells of the ED and MD in response to Mock or Sorbitol 300 mM treatment for 1 day, quantified as mean gray value (arbitrary units; a.u.). Box plots as described in (A). $N \geq 14$ cells (3-5 cells/root) per root area/treatment. RM one-way ANOVA with Kruskal-Wallis multiple comparison test. Alphabet letters indicate significant differences with $P \leq 0.05$. **C.** Amount of the mCherry signal intensity of *pATHE::mCherry-N7* epidermal cells at the root differentiated zones indicated in Fig 2(D) in response to Mock or Sorbitol 300 nM treatment for one day, quantified as mean gray value (arbitrary units; a.u.). Box plots as described in (A). $N \geq 51$ cells (8-15 cells/root) per root area/treatment. RM one-way ANOVA with Kruskal-Wallis multiple comparison test. Alphabet letters indicate significant differences with $P \leq 0.05$.

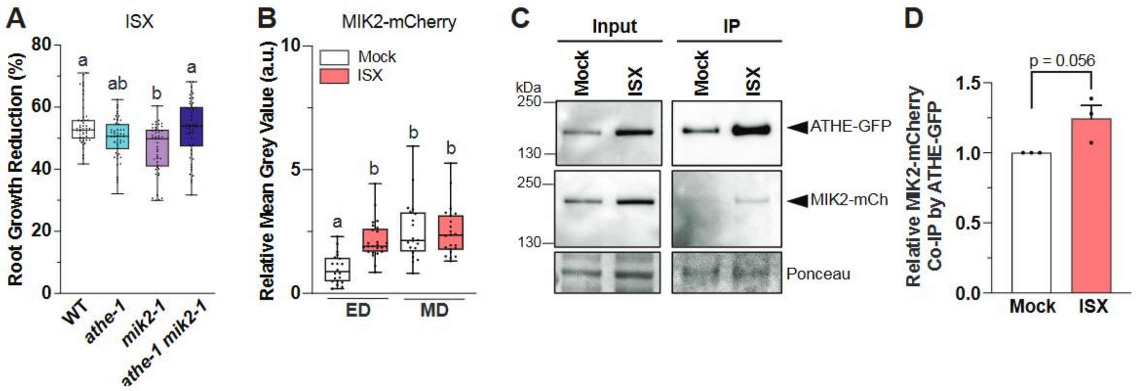

**Figure S6. ATHE interaction with MIK2 is not mediated by ISX. A.** Root growth reduction of Mock or 2 nM Isoxaben (ISX)-treated wild-type (WT; Col-0), *athe-1*, *mik2-1*, and *athe-1 mik2-1* seedlings relative to Mock-treated ones, after 5 days post treatment (dpt). Box plots: centerlines show the medians; box limits indicate the 25th and 75th percentiles; whiskers extend to the minimum and maximum. $N \geq 45$ roots from 3 experiments (≥ 15 roots/experiments). Repeated measures (RM) one-way ANOVA with Kruskal-Wallis multiple comparison test. **B.** Amount of MIK2-mCherry at the plasma membranes of epidermal cells of ED and MD in response to Mock or ISX 300 nM treatment for 1 day, quantified as mean gray value (arbitrary units; a.u.). Box plots as described in (A). $N \geq 20$ cells (3-5 cells/root) per root area/treatment. RM one-way ANOVA with Kruskal-Wallis multiple comparison test. Alphabet letters indicate significant differences with $P \leq 0.05$. **C.** Representative western blot co-immunoprecipitation analysis of ATHE-GFP and MIK2-mCherry using GFP-trap beads, in Mock- or ISX (300nM)-exposed roots at 1dpt, using anti-GFP antibody (ATHE-GFP) and anti-RFP antibody (MIK2-mCherry). Ponceau S staining provides loading control. **D.** ATHE-GFP and MIK2-Cherry signal quantification from co-immunoprecipitation immunoblots as in (C). The intensity of the MIK2-mCherry band in every condition was normalized to the intensity of ATHE-GFP band in the same condition (Mock or ISX). Av ± SEM, $N = 3$ independent experiments. RM *t*-test.

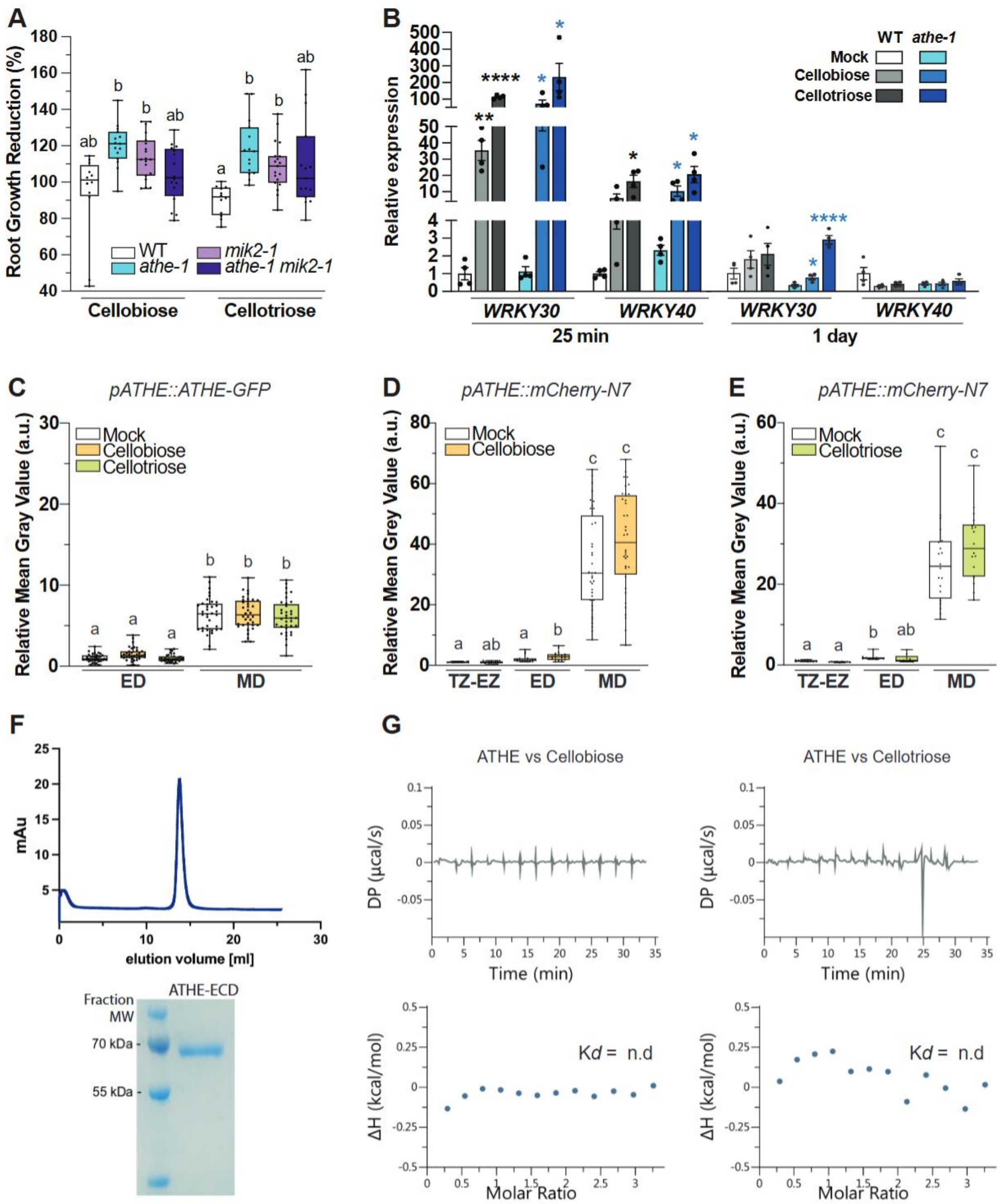

**Figure S7. ATHENA participates in the sensing of cellulose oligomers.**

**A.** Root growth reduction of Mock, Cellobiose 100 µM or Cellotriose 100 µM-treated WT, *athe-1*, *mik2-1*, and *athe-1 mik2-1* seedlings relative to Mock-treated ones, after 3 dpt. The box plots are shown: centerlines show the medians; box limits indicate the 25th and 75th percentiles; whiskers extend to the minimum and maximum.. $N \geq 12$ roots. RM one-way ANOVA *(Brown-Forsythe and Welch)* test. Alphabet letters indicate significant differences with $P \leq 0.05$. **B.** Gene expression of the Cellooligomers-responsive genes *WRKY30* and *WRKY40* relative to *AtGAPDH* in WT and *athe-1* roots in Mock or upon cellobiose 100 μM or cellotriose 100 μM at 25 min and 1 day post treatment. Av ± SEM, *N* = 4 independent replicates from ≥ 20 roots. RM *t*-test. *P ≤ 0.05, **P ≤ 0.01, ***P ≤ 0.001, ****P ≤ 0.0001. Asterisks in black indicate the difference with WT Mock. Asterisks in blue indicate the differences in *athe-1* Mock versus *athe-1* treatment. **C.** Amount of ATHE-GFP at the plasma membranes of epidermal cells of ED and MD in response to Mock, Cellobiose 100 μM, or Cellotriose 100 μM treatment for 1 day, quantified as mean gray value (arbitrary units; a.u.). Box plots as described in (A). *N* ≥ 31 cells per root area/treatment (>9 cells/root and ≥ 3 roots/experiments). RM one-way ANOVA with Tukey's multiple comparisons test. Alphabet letters indicate significant differences with P ≤ 0.05. **D and E.** Amount of the mCherry signal intensity of *pATHE::mCherry-N7* epidermal cells at the root differentiated zones indicated in Fig 2(D) in response to Cellobiose 100 μM (E) or Cellotriose 100 μM (F) treatment for 1 day. Box plots as described in (A). *N* ≥ 32 cells (6-14 cells/root and ≥ 3 roots/experiments) per root area/treatment were quantified. RM one-way ANOVA with Kruskal-Wallis multiple comparison test. Alphabet letters indicate significant differences with P ≤ 0.05. **F.** Size exclusion chromatography (upper panel) and SDS-PAGE analysis (lower panel) of ATHE-ECD expressed in insect cells. **G.** Isothermal titration calorimetry (ITC) experiments of ATHE-ECD vs Cellobiose (left) and Cellotriose (right); n.d = no detected binding

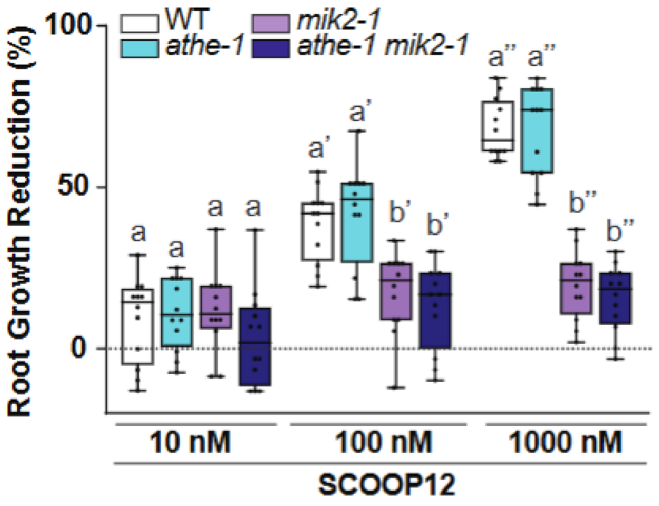

**Figure S8. ATHE interaction with MIK2 is not mediated by SCOOP12.**

Root Growth reduction of Wild-Type (WT; Col-0), *athe-1*, *mik2-1*, and *athe-1 mik2-1* seedlings exposed 7 days to 10, 100 and 1000 nM SCOOP12 peptides-treated relative to Mock-treated ones. Box plots: centerlines show the medians; box limits indicate the 25th and 75th percentiles; whiskers extend to the minimum and maximum. *N* = 12 roots/replicate. Repeated measures (RM) one-way ANOVA (Brown-Forsythe and Welch) test compared to the WT under each condition.Alphabet letters with different symbols (′ or ″) indicate significant differences within each treatment with $P \leq 0.05$.

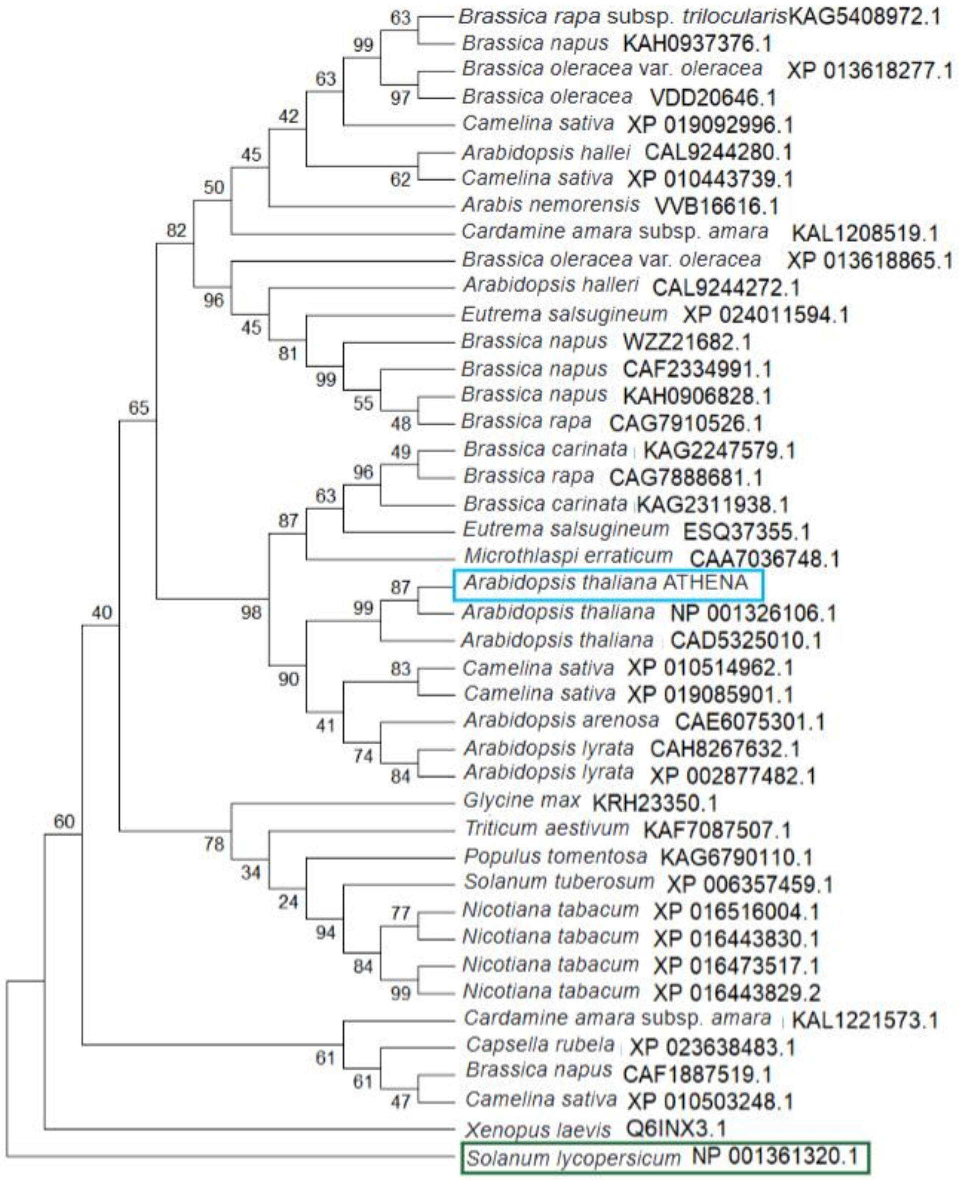

**Figure S9. Phylogenetic tree of ATHENA-like proteins in different plant species.** Phylogenetic analysis of ATHENA-like proteins in selected plant species by doing a blast The outgroup *Xenopus laevis* malectin protein Q6INX3 was included in the tree as root tree. The percentage of replicate trees in which the associated taxa clustered together (1000 replicates) is shown next to the branches. In blue is marked ATHENA from *Arabidopsis thaliana* and in green the phylogenetically closest protein from *Solanum lycopersicum.*

**Movie S1. ATHE-GFP localizes in foci structures in the PM.** Representative spinning disk confocal movie at the PM focal plane of an epidermal root cell at the MD of the ATHE-GFP line. Scale bar = 5 µm.

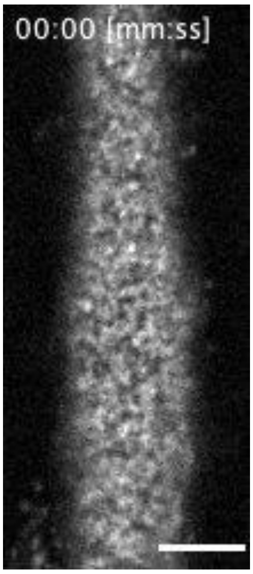

**Table S1. Differential MS2 spectra detection in organelle-IP upon Fo5176 infection.** MS2 spectra detected from four (1-4) independent experiments of Mock (m) and Fo-infected (i) root samples. The MEE39/ATHE protein is highlighted in yellow.

**Table S2. Label-free quantification of ATHE-GFP IP upon Fo5176 infection.** MS-2 spectra from three (1-3) independent experiments of Mock (m) and Fo- infected (i) root samples. ATHE and MIK2 are highlighted in yellow and green, respectively.

**Table S3. List of primers used in this work.**

**Table S4. Sequences of RALF peptides used in this work**

**Table S1.** Differential MS2 spectra detection in organelle-IP upon Fo5176 infection.  MS2 spectra detected from four (1-4) independent experiments of mock (m) and Fo-infected (i) root samples. The MEE39/ATHE protein is highlighted in yellow.

| ARA7-YFP IP profile | | | | | | | | | | | |
|---|---|---|---|---|---|---|---|---|---|---|---|
| | | | | MS2 spectra count | | | | | | | |
| Description | | | | ARA7-YFP_m | | | | ARA7-YFP_i | | | |
| | Accession Number | t-test (p-value) | Change | Rep1 | Rep2 | Rep3 | Rep4 | Rep1 | Rep2 | Rep3 | Rep4 |
| PE=2 SV=1 | sp\|Q9M1P7\|BOR2_ARATH | 0,00054 | 0 | 1 | 1 | 2 | 1 | 0 | 0 | 0 | 0 |
| thaliana GN=PIN3 PE=1 SV=1 (sp\|Q9S7Z8\|PIN3_ARATH) | sp\|Q9S7Z8\|PIN3_ARATH [3] | 0,026 | 0 | 3 | 1 | 5 | 1 | 0 | 0 | 0 | 0 |
| GN=FLA9 PE=1 SV=1 | sp\|Q9ZWA8\|FLA9_ARATH | 0,041 | 0 | 1 | 0 | 2 | 1 | 0 | 0 | 0 | 0 |
| GN=PVA22 PE=1 SV=1 | sp\|B9DHD7\|VAP22_ARATH | 0,014 | 0,06 | 3 | 1 | 5 | 2 | 0 | 1 | 0 | 0 |
| SV=1 | sp\|Q9FJV8\|PEN5_ARATH | 0,032 | 0,06 | 11 | 10 | 11 | 0 | 0 | 1 | 1 | 0 |
| GN=At4g11320 PE=2 SV=1 | sp\|Q9SUS9\|CPR4_ARATH | 0,027 | 0,08 | 16 | 12 | 16 | 1 | 0 | 1 | 1 | 1 |
| thaliana GN=SMT3 PE=2 SV=1 | sp\|Q94JS4\|SMT3B_ARATH | 0,0054 | 0,2 | 11 | 7 | 11 | 5 | 3 | 1 | 3 | 0 |
| GN=CYP71A16 PE=2 SV=1 | sp\|Q9FH66\|C71AG_ARATH | 0,034 | 0,2 | 24 | 10 | 21 | 6 | 4 | 6 | 0 | 1 |
| GN=ABCG14 PE=2 SV=1 | sp\|Q9C6W5\|AB14G_ARATH | 0,046 | 0,2 | 3 | 1 | 4 | 1 | 0 | 1 | 0 | 0 |
| thaliana GN=At2g21160 PE=2 SV=3 | sp\|P45434\|SSRA_ARATH | 0,003 | 0,3 | 4 | 5 | 5 | 5 | 3 | 2 | 1 | 0 |
| GN=B3GALT7 PE=2 SV=1 | sp\|Q6NQB7\|B3GT7_ARATH | 0,023 | 0,3 | 2 | 2 | 1 | 2 | 1 | 0 | 0 | 1 |
| thaliana GN=SMT2 PE=1 SV=2 | sp\|Q39227\|SMT2_ARATH | 0,03 | 0,3 | 10 | 6 | 17 | 6 | 3 | 3 | 2 | 2 |
| SV=1 | sp\|Q9FKT5\|THOC3_ARATH | 0,049 | 0,3 | 1 | 1 | 1 | 2 | 1 | 1 | 0 | 0 |
| GN=ATR1 PE=1 SV=1 | sp\|Q9SB48\|NCPR1_ARATH | 0,031 | 0,4 | 16 | 8 | 22 | 14 | 9 | 8 | 6 | 2 |
| PE=1 SV=1 (sp\|Q38890\|GUN25_ARATH) | sp\|Q38890\|GUN25_ARATH [2] | 0,042 | 0,4 | 6 | 5 | 4 | 3 | 2 | 4 | 2 | 0 |
| Syntaxin-22 OS=Arabidopsis thaliana GN=SYP22 PE=1 SV=1 | sp\|P93654\|SYP22_ARATH | 0,042 | 0,4 | 3 | 2 | 4 | 2 | 2 | 2 | 1 | 0 |
| dehydratase PASTICCINO 2 OS=Arabidopsis thaliana GN=PAS2 | sp\|Q8VZB2\|PAS2_ARATH | 0,01 | 0,5 | 4 | 6 | 4 | 3 | 2 | 1 | 2 | 2 |
| Apyrase 2 OS=Arabidopsis thaliana GN=APY2 PE=1 SV=1 | sp\|Q9SPM5\|APY2_ARATH | 0,016 | 0,5 | 7 | 5 | 9 | 7 | 5 | 4 | 2 | 2 |
| GN=ERD3 PE=2 SV=1 | sp\|Q94II3\|PMTL_ARATH | 0,026 | 0,5 | 17 | 13 | 15 | 7 | 4 | 5 | 10 | 6 |
| OS=Arabidopsis thaliana GN=CESA1 PE=1 SV=1 | sp\|O48946\|CESA1_ARATH [3] | 0,027 | 0,5 | 20 | 14 | 24 | 15 | 6 | 6 | 13 | 13 |
| GN=At1g29470 PE=2 SV=1 | sp\|Q6NPR7\|PMTO_ARATH | 0,029 | 0,5 | 23 | 19 | 33 | 17 | 15 | 13 | 12 | 11 |
| GN=NPF6.3 PE=1 SV=1 | sp\|Q05085\|PTR7_ARATH | 0,029 | 0,5 | 3 | 5 | 4 | 2 | 2 | 1 | 1 | 2 |
| OS=Arabidopsis thaliana GN=CESA3 PE=1 SV=2 | sp\|Q941L0\|CESA3_ARATH [3] | 0,034 | 0,5 | 29 | 36 | 32 | 16 | 12 | 10 | 15 | 21 |
| GN=MSBP2 PE=1 SV=1 | sp\|Q9M2Z4\|MSBP2_ARATH | 0,043 | 0,5 | 4 | 7 | 4 | 5 | 3 | 1 | 4 | 1 |
| Synaptotagmin-5 OS=Arabidopsis thaliana GN=SYT5 PE=2 SV=1 | sp\|Q8L706\|SYT5_ARATH | 0,026 | 0,6 | 7 | 13 | 10 | 9 | 6 | 6 | 6 | 6 |
| type OS=Arabidopsis thaliana GN=ECA1 PE=1 SV=2 | sp\|P92939\|ECA1_ARATH [2] | 0,028 | 0,6 | 58 | 67 | 45 | 47 | 34 | 24 | 44 | 39 |
| GN=ASNAP2 PE=1 SV=1 | sp\|Q9SPE6\|SNAA2_ARATH | 0,05 | 0,6 | 10 | 7 | 10 | 6 | 4 | 6 | 7 | 4 |
| Calreticulin-1 OS=Arabidopsis thaliana GN=CRT1 PE=1 SV=1 | sp\|O04151\|CALR1_ARATH | 0,029 | 0,7 | 58 | 62 | 45 | 43 | 37 | 44 | 31 | 31 |

| | | | | | | | | | | | |
|---|---|---|---|---|---|---|---|---|---|---|---|
| GN=RPL30B PE=3 SV=1 (sp\|Q8VZ19\|RL302_ARATH) | sp\|Q8VZ19\|RL302_ARATH [2] | 0,048 | 0,7 | 8 | 8 | 10 | 10 | 6 | 5 | 4 | 9 |
| GN=RABG3D PE=2 SV=1 (sp\|Q9C820\|RAG3D_ARATH) | sp\|Q9C820\|RAG3D_ARATH [2] | 0,038 | 0,8 | 26 | 31 | 22 | 22 | 20 | 21 | 17 | 19 |
| Patellin-3 OS=Arabidopsis thaliana GN=PATL3 PE=1 SV=2 | sp\|Q56Z59\|PATL3_ARATH | 0,029 | 1,3 | 29 | 39 | 29 | 26 | 39 | 36 | 44 | 45 |
| OS=Arabidopsis thaliana GN=ACA8 PE=1 SV=1 | sp\|Q9LF79\|ACA8_ARATH [2] | 0,047 | 1,4 | 14 | 13 | 20 | 13 | 24 | 17 | 19 | 24 |
| GEM-like protein 1 OS=Arabidopsis thaliana GN=FIP1 PE=1 SV=1 | sp\|Q9SE96\|GEML1_ARATH | 0,044 | 1,5 | 6 | 4 | 6 | 4 | 9 | 9 | 6 | 6 |
| PE=2 SV=1 | sp\|O65902\|ACAP1_ARATH | 0,032 | 1,6 | 3 | 4 | 4 | 4 | 4 | 6 | 7 | 6 |
| OS=Arabidopsis thaliana GN=PAP3 PE=2 SV=1 | sp\|O82291\|PAP3_ARATH | 0,038 | 1,6 | 4 | 4 | 3 | 3 | 4 | 6 | 6 | 5 |
| alpha, chloroplastic OS=Arabidopsis thaliana GN=CAC3 PE=1 | sp\|Q9LD43\|ACCA_ARATH | 0,0054 | 1,7 | 11 | 14 | 10 | 11 | 16 | 23 | 17 | 21 |
| GN=psbA PE=3 SV=1 (sp\|A0ZZ15\|PSBA_GOSBA) | sp\|A0ZZ15\|PSBA_GOSBA [52] | 0,014 | 1,8 | 2 | 1 | 3 | 1 | 3 | 4 | 3 | 4 |
| thaliana GN=PECT1 PE=1 SV=1 | sp\|Q9ZVI9\|PECT1_ARATH | 0,016 | 1,8 | 2 | 4 | 6 | 5 | 6 | 8 | 9 | 8 |
| GN=RPS10C PE=2 SV=2 (sp\|Q9LTF2\|RS103_ARATH) | sp\|Q9LTF2\|RS103_ARATH [2] | 0,043 | 1,8 | 5 | 10 | 4 | 5 | 9 | 14 | 13 | 8 |
| OS=Arabidopsis thaliana GN=PAP2 PE=2 SV=1 | sp\|O49629\|PAP2_ARATH | 0,0088 | 1,9 | 3 | 4 | 4 | 1 | 5 | 6 | 5 | 6 |
| PE=2 SV=1 | sp\|Q9CAG3\|LOX6_ARATH | 0,023 | 1,9 | 5 | 6 | 7 | 4 | 10 | 8 | 15 | 11 |
| (sp\|P0CG89\|H4_SOYBN) | sp\|P0CG89\|H4_SOYBN [14] | 0,0025 | 2,1 | 2 | 1 | 3 | 2 | 5 | 5 | 4 | 4 |
| SV=1 | sp\|Q9FGY1\|BXL1_ARATH | 0,0028 | 2,1 | 7 | 4 | 7 | 9 | 12 | 12 | 15 | 16 |
| GN=ALATS PE=1 SV=3 (sp\|P36428\|SYA_ARATH) | sp\|P36428\|SYA_ARATH [4] | 0,019 | 2,2 | 8 | 8 | 9 | 6 | 13 | 18 | 27 | 14 |
| GN=At5g53140 PE=2 SV=1 | sp\|Q94AT1\|P2C76_ARATH | 0,037 | 2,2 | 1 | 0 | 1 | 1 | 2 | 1 | 2 | 1 |
| PE=1 SV=1 | sp\|Q9FFT4\|PDC2_ARATH | 0,037 | 2,2 | 1 | 0 | 1 | 1 | 2 | 1 | 2 | 1 |
| thaliana GN=GAMMACA1 PE=1 SV=1 | sp\|Q9FWR5\|GCA1_ARATH | 0,04 | 2,3 | 2 | 0 | 2 | 2 | 4 | 2 | 3 | 5 |
| PE=3 SV=2 | sp\|Q680P8\|RS29_ARATH | 0,027 | 2,4 | 1 | 0 | 1 | 1 | 2 | 2 | 1 | 2 |
| OS=Arabidopsis thaliana GN=PAP1 PE=1 SV=1 | sp\|O81439\|PAP1_ARATH | 0,029 | 2,4 | 3 | 2 | 2 | 1 | 3 | 6 | 4 | 7 |
| OS=Arabidopsis thaliana GN=RH2 PE=2 SV=2 | sp\|Q94A52\|RH2_ARATH [2] | 0,042 | 2,4 | 3 | 2 | 5 | 3 | 5 | 9 | 13 | 6 |
| thaliana GN=ASN2 PE=1 SV=1 | sp\|Q9LV77\|ASNS2_ARATH | 0,041 | 2,8 | 0 | 0 | 1 | 1 | 2 | 1 | 1 | 1 |
| PE=1 SV=1 | sp\|Q9ZUB3\|SPD1_ARATH | 0,042 | 3 | 1 | 0 | 0 | 1 | 2 | 2 | 1 | 1 |
| OS=Arabidopsis thaliana GN=MEE39 PE=2 SV=1 | sp\|C0LGP2\|MEE39_ARATH | 0,002 | 3,1 | 1 | 0 | 1 | 1 | 3 | 3 | 2 | 2 |
| prephenate aminotransferase OS=Arabidopsis thaliana GN=PAT | sp\|Q9SIE1\|PAT_ARATH | 0,022 | 3,1 | 1 | 0 | 1 | 2 | 4 | 4 | 2 | 2 |
| SV=2 | sp\|O80690\|BGL46_ARATH | 0,035 | 3,2 | 0 | 0 | 1 | 1 | 2 | 2 | 1 | 1 |
| OS=Arabidopsis thaliana GN=XTH24 PE=1 SV=2 | sp\|P24806\|XTH24_ARATH | 0,032 | 3,3 | 2 | 0 | 1 | 7 | 9 | 6 | 13 | 7 |
| PE=2 SV=4 | sp\|P93733\|PLDB1_ARATH | 0,041 | 3,4 | 1 | 1 | 2 | 0 | 4 | 4 | 5 | 1 |
| GN=ACC1 PE=1 SV=1 (sp\|Q38970\|ACC1_ARATH) | sp\|Q38970\|ACC1_ARATH [2] | 0,0061 | 3,5 | 2 | 0 | 2 | 2 | 4 | 4 | 6 | 7 |
| (Fragment) OS=Lavandula lanata GN=rbcL PE=3 SV=1 | sp\|Q33600\|RBL_LAVLA [12] | 0,021 | 3,6 | 3 | 0 | 1 | 1 | 3 | 4 | 6 | 6 |
| thaliana GN=ASP1 PE=2 SV=1 | sp\|P46643\|AAT1_ARATH | 0,034 | 3,8 | 3 | 1 | 3 | 1 | 8 | 14 | 5 | 5 |
| OS=Arabidopsis thaliana GN=PAP9 PE=2 SV=1 | sp\|Q9M2P7\|PAP9_ARATH | 0,0059 | 3,9 | 0 | 0 | 1 | 1 | 2 | 1 | 2 | 2 |
| OS=Arabidopsis thaliana GN=BOU PE=2 SV=1 | sp\|Q93XM7\|MCAT_ARATH | 0,015 | 4 | 0 | 0 | 1 | 1 | 2 | 3 | 2 | 1 |
| chloroplastic OS=Arabidopsis thaliana GN=RCA PE=1 SV=2 | sp\|P10896\|RCA_ARATH [3] | 0,019 | 4 | 1 | 0 | 0 | 2 | 4 | 4 | 2 | 2 |

| | | | | | | | | | | | |
|---|---|---|---|---|---|---|---|---|---|---|---|
| OS=Arabidopsis thaliana GN=At2g02050 PE=3 SV=1 | sp\|Q9SKC9\|NDUB7_ARATH | 0,021 | 4,3 | 0 | 0 | 0 | 1 | 1 | 1 | 1 | 1 |
| OS=Arabidopsis thaliana GN=At1g62640 PE=2 SV=2 | sp\|P49243\|FABH_ARATH | 0,021 | 4,3 | 0 | 0 | 0 | 1 | 1 | 1 | 1 | 1 |
| SV=1 | sp\|Q9XEX2\|PRX2B_ARATH | 0,04 | 4,4 | 0 | 0 | 2 | 0 | 1 | 3 | 2 | 2 |
| PE=2 SV=2 | sp\|Q8LC69\|ATL8_ARATH | 0,02 | 5,3 | 0 | 0 | 0 | 1 | 1 | 1 | 2 | 1 |
| thaliana GN=GRIMP PE=1 SV=1 | sp\|Q9LX99\|GRIMP_ARATH | 0,028 | 5,5 | 1 | 0 | 0 | 0 | 1 | 1 | 1 | 2 |
| thaliana GN=At2g43090 PE=1 SV=1 | sp\|Q9ZW85\|LEUD3_ARATH | 0,0093 | 6,8 | 0 | 0 | 1 | 0 | 2 | 2 | 1 | 1 |
| SV=1 | sp\|Q9FMR9\|RIN1_ARATH | 0,042 | 7,5 | 0 | 0 | 1 | 0 | 2 | 1 | 3 | 1 |
| OS=Arabidopsis thaliana GN=PME3 PE=2 SV=2 | sp\|O49006\|PME3_ARATH [4] | 0,00027 | 9,1 | 0 | 0 | 1 | 0 | 2 | 2 | 2 | 2 |
| Phospholipase A I OS=Arabidopsis thaliana GN=PLA1 PE=2 SV=1 | sp\|F4HX15\|LPAI_ARATH | 0,023 | 9,2 | 1 | 0 | 0 | 0 | 4 | 1 | 3 | 1 |
| OS=Arabidopsis thaliana GN=RFS6 PE=2 SV=2 | sp\|Q8RX87\|RFS6_ARATH | 0,0015 | 9,3 | 1 | 0 | 4 | 3 | 20 | 17 | 26 | 12 |
| Patellin-5 OS=Arabidopsis thaliana GN=PATL5 PE=1 SV=2 | sp\|Q9M0R2\|PATL5_ARATH | 0,0002 | ND | 0 | 0 | 0 | 0 | 2 | 2 | 2 | 1 |
| GN=GSH2 PE=2 SV=3 | sp\|P46416\|GSHB_ARATH | 0,0029 | ND | 0 | 0 | 0 | 0 | 2 | 1 | 1 | 1 |
| GN=TIC20-II PE=2 SV=1 | sp\|O82251\|TI202_ARATH | 0,027 | ND | 0 | 0 | 0 | 0 | 0 | 1 | 1 | 1 |
| GN=APT2 PE=2 SV=1 | sp\|Q42563\|APT2_ARATH | 0,029 | ND | 0 | 0 | 0 | 0 | 2 | 1 | 2 | 0 |
| At5g05200, chloroplastic OS=Arabidopsis thaliana GN=At5g05200 | sp\|Q9ASX5\|Y5520_ARATH | 0,03 | ND | 0 | 0 | 0 | 0 | 1 | 1 | 1 | 0 |
| B OS=Arabidopsis thaliana GN=At3g18410 PE=2 SV=1 | sp\|Q94C12\|NDBAB_ARATH | 0,03 | ND | 0 | 0 | 0 | 0 | 1 | 1 | 1 | 0 |

| free-GFP IP profile | | | | | | | | | | | |
|---|---|---|---|---|---|---|---|---|---|---|---|
| | | | | MS2 spectra count | | | | | | | |
| Description | | | | freeGFP_m | | | | freeGFP_i | | | |
| | Accession Number | t-test (p-value) | Change | Rep1 | Rep2 | Rep3 | Rep4 | Rep1 | Rep2 | Rep3 | Rep4 |
| subunit STT3A OS=Arabidopsis thaliana GN=STT3A PE=2 SV=1 | sp\|Q93ZY3\|STT3A_ARATH | 0,029 | 0 | 1 | 2 | 2 | 0 | 0 | 0 | 0 | 0 |
| membrane-type OS=Arabidopsis thaliana GN=ACA11 PE=1 SV=1 | sp\|Q9M2L4\|ACA11_ARATH [2] | 0,046 | 0 | 3 | 1 | 0 | 1 | 0 | 0 | 0 | 0 |
| GN=DRG3 PE=1 SV=1 | sp\|Q9SVA6\|DRG3_ARATH | 0,013 | 0 | 1 | 1 | 1 | 2 | 0 | 0 | 0 | 0 |
| glutamate/aspartate-prephenate aminotransferase | sp\|Q9SIE1\|PAT_ARATH [2] | 0,038 | 0 | 3 | 0 | 2 | 1 | 0 | 0 | 0 | 0 |
| Myosin-17 OS=Arabidopsis thaliana GN=XI-K PE=1 SV=2 | sp\|F4K5J1\|MYO17_ARATH | 0,021 | 0 | 3 | 1 | 1 | 1 | 0 | 0 | 0 | 0 |
| thaliana GN=PPA2 PE=2 SV=2 (sp\|P21216\|IPYR2_ARATH) | sp\|P21216\|IPYR2_ARATH [3] | 0,022 | 0 | 3 | 1 | 1 | 1 | 0 | 0 | 0 | 0 |
| thaliana GN=CBF5 PE=1 SV=1 | sp\|Q9LD90\|CBF5_ARATH | 0,044 | 0 | 3 | 1 | 1 | 1 | 0 | 0 | 0 | 0 |
| isoform 2 OS=Arabidopsis thaliana GN=ACG12 PE=2 SV=1 | sp\|Q9FJI5\|G6PD6_ARATH [2] | 0,022 | 0 | 3 | 1 | 2 | 1 | 0 | 0 | 0 | 0 |
| Protein RER1A OS=Arabidopsis thaliana GN=RER1A PE=1 SV=1 | sp\|O48670\|RER1A_ARATH | 0,016 | 0 | 3 | 2 | 1 | 1 | 0 | 0 | 0 | 0 |
| PE=2 SV=2 | (+1) | 0,02 | 0 | 3 | 1 | 1 | 2 | 0 | 0 | 0 | 0 |
| Annexin D4 OS=Arabidopsis thaliana GN=ANN4 PE=2 SV=1 | sp\|Q9ZVJ6\|ANXD4_ARATH | 0,044 | 0 | 3 | 1 | 0 | 3 | 0 | 0 | 0 | 0 |
| type OS=Arabidopsis thaliana GN=ECA1 PE=1 SV=2 | sp\|P92939\|ECA1_ARATH [2] | 0,00053 | 0 | 3 | 3 | 2 | 1 | 0 | 0 | 0 | 0 |
| GN=NPC3 PE=2 SV=1 | sp\|Q9SRQ6\|NPC3_ARATH | 0,0057 | 0 | 3 | 3 | 2 | 1 | 0 | 0 | 0 | 0 |
| GN=PDIL1-4 PE=1 SV=1 | sp\|Q9FF55\|PDI14_ARATH | 0,036 | 0 | 3 | 4 | 0 | 2 | 0 | 0 | 0 | 0 |

| | | | | | | | | | | | |
|---|---|---|---|---|---|---|---|---|---|---|---|
| GN=RBP45B PE=1 SV=1 | sp\|Q9SAB3\|RB45B_ARATH | 0,014 | 0 | 4 | 2 | 1 | 2 | 0 | 0 | 0 | 0 |
| thaliana GN=GSL-OH PE=2 SV=1 | sp\|Q9SKK4\|GSL_ARATH | 0,019 | 0 | 3 | 1 | 1 | 4 | 0 | 0 | 0 | 0 |
| OS=Arabidopsis thaliana GN=CESA1 PE=1 SV=1 | sp\|O48946\|CESA1_ARATH [6] | 0,0034 | 0 | 4 | 2 | 2 | 2 | 0 | 0 | 0 | 0 |
| OS=Arabidopsis thaliana GN=P5CSB PE=2 SV=1 | sp\|P54888\|P5CS2_ARATH [2] | 0,0097 | 0 | 4 | 1 | 2 | 5 | 0 | 0 | 0 | 0 |
| GN=ATJ3 PE=1 SV=2 (sp\|Q94AW8\|DNAJ3_ARATH) | sp\|Q94AW8\|DNAJ3_ARATH [3] | 0,035 | 0 | 8 | 4 | 1 | 3 | 0 | 0 | 0 | 0 |
| thaliana GN=RH51 PE=2 SV=1 | sp\|Q9LIH9\|RH51_ARATH | 0,0071 | 0 | 6 | 4 | 2 | 6 | 0 | 0 | 0 | 0 |
| SV=3 (sp\|Q38858\|CALR2_ARATH) | sp\|Q38858\|CALR2_ARATH [3] | 0,0058 | 0,04 | 5 | 9 | 4 | 4 | 0 | 0 | 0 | 1 |
| OS=Arabidopsis thaliana GN=RHM1 PE=1 SV=1 | sp\|Q9SYM5\|RHM1_ARATH [2] | 0,026 | 0,05 | 4 | 6 | 1 | 7 | 0 | 0 | 0 | 1 |
| PE=2 SV=3 | sp\|P49690\|RL23_ARATH | 0,016 | 0,05 | 3 | 6 | 3 | 9 | 0 | 1 | 0 | 0 |
| PE=1 SV=1 | sp\|Q9C942\|CSE_ARATH | 0,0067 | 0,07 | 5 | 3 | 2 | 4 | 0 | 0 | 0 | 1 |
| GN=BETAC-AD PE=1 SV=2 (sp\|O81742\|APBLC_ARATH) | sp\|O81742\|APBLC_ARATH [2] | 0,012 | 0,1 | 3 | 2 | 1 | 2 | 0 | 1 | 0 | 0 |
| SV=3 | sp\|P36428\|SYA_ARATH | 0,0072 | 0,1 | 3 | 1 | 2 | 3 | 0 | 0 | 0 | 1 |
| thaliana GN=UXS2 PE=1 SV=1 (sp\|Q9LZI2\|UXS2_ARATH) | sp\|Q9LZI2\|UXS2_ARATH [2] | 0,038 | 0,1 | 3 | 6 | 3 | 1 | 0 | 1 | 0 | 1 |
| OS=Arabidopsis thaliana GN=At3g22845 PE=2 SV=1 | sp\|Q9LIL4\|P24B3_ARATH | 0,042 | 0,1 | 4 | 8 | 6 | 1 | 0 | 0 | 0 | 3 |
| thaliana GN=OEP37 PE=1 SV=2 | sp\|O80565\|OEP37_ARATH | 0,034 | 0,2 | 1 | 1 | 1 | 1 | 0 | 0 | 0 | 1 |
| GN=LACS4 PE=2 SV=1 | sp\|Q9T0A0\|LACS4_ARATH | 0,0011 | 0,2 | 3 | 3 | 2 | 2 | 0 | 1 | 0 | 1 |
| GN=RPS15AA PE=2 SV=2 (sp\|P42798\|R15A1_ARATH) | sp\|P42798\|R15A1_ARATH [2] | 0,034 | 0,2 | 5 | 6 | 2 | 9 | 0 | 1 | 0 | 3 |
| GN=RABH1B PE=1 SV=1 (sp\|O80501\|RAH1B_ARATH) | sp\|O80501\|RAH1B_ARATH [2] | 0,013 | 0,2 | 10 | 8 | 3 | 7 | 0 | 2 | 0 | 3 |
| OS=Arabidopsis thaliana GN=BIG5 PE=1 SV=2 | sp\|F4IXW2\|BIG5_ARATH | 0,038 | 0,3 | 4 | 2 | 2 | 4 | 0 | 2 | 0 | 1 |
| OS=Arabidopsis thaliana GN=At1g26630 PE=1 SV=1 | sp\|Q93VP3\|IF5A2_ARATH [2] | 0,018 | 0,3 | 5 | 4 | 2 | 7 | 2 | 1 | 0 | 2 |
| homolog A OS=Arabidopsis thaliana GN=RPN3A PE=1 SV=3 | sp\|Q9LNU4\|PSD3A_ARATH [2] | 0,033 | 0,3 | 8 | 4 | 9 | 12 | 0 | 6 | 0 | 4 |
| GN=RPL26A PE=2 SV=2 | sp\|P51414\|RL261_ARATH | 0,022 | 0,3 | 8 | 10 | 9 | 7 | 0 | 3 | 0 | 8 |
| GN=RABE1C PE=1 SV=1 (sp\|P28186\|RAE1C_ARATH) | sp\|P28186\|RAE1C_ARATH [12] | 0,012 | 0,3 | 8 | 10 | 7 | 11 | 0 | 3 | 2 | 7 |
| SV=2 | sp\|Q9M0Y8\|NSF_ARATH | 0,044 | 0,4 | 13 | 11 | 13 | 4 | 4 | 1 | 6 | 5 |
| GN=RPL18AB PE=2 SV=2 (sp\|P51418\|R18A2_ARATH) | sp\|P51418\|R18A2_ARATH [2] | 0,024 | 0,4 | 12 | 17 | 11 | 21 | 4 | 7 | 6 | 11 |
| GN=NOP5-2 PE=2 SV=1 | sp\|Q9MAB3\|NOP5B_ARATH | 0,042 | 0,5 | 14 | 10 | 9 | 9 | 2 | 9 | 4 | 7 |
| GN=Ighg1 PE=1 SV=1 | (+1) | 0,014 | 1,3 | 18 | 26 | 23 | 26 | 29 | 30 | 29 | 34 |
| thaliana GN=VHA-B1 PE=2 SV=2 (sp\|P11574\|VATB1_ARATH) | sp\|P11574\|VATB1_ARATH [3] | 0,047 | 1,4 | 35 | 31 | 32 | 25 | 37 | 34 | 51 | 51 |
| SV=1 (sp\|Q6VAG0\|TBA2_GOSHI) | sp\|Q6VAG0\|TBA2_GOSHI [13] | 0,047 | 1,5 | 19 | 17 | 24 | 29 | 26 | 31 | 45 | 36 |
| GN=RPS16A PE=2 SV=1 | sp\|Q9SK22\|RS161_ARATH | 0,0062 | 1,9 | 4 | 6 | 4 | 6 | 12 | 8 | 10 | 8 |
| GN=RPL40A PE=1 SV=1 | (+36) | 0,044 | 2,5 | 3 | 4 | 4 | 6 | 12 | 7 | 16 | 6 |
| GN=RPL21A PE=2 SV=2 | sp\|Q43291\|RL211_ARATH (+1) | 0,012 | 2,8 | 1 | 1 | 2 | 0 | 2 | 3 | 4 | 3 |

**Table S2.** Label-free quantification of ATHE-GFP IP upon Fo5176 infection. MS-2 spectra from three (1-3) independent experiments of mock (m) and Fo-infected (i) root samples. ATHE and MIK2 are highlighted in yellow and green, respectively.

| | | | | LFQ inten | | | | | |
|---|---|---|---|---|---|---|---|---|---|
| | | | | ATHE_m | | | ATHE_i | | |
| Name | AGI number | ANOVA p value | Significant pairs | Rep1 | Rep2 | Rep3 | Rep1 | Rep2 | Rep3 |
| Ribosomal protein L16p/L10e family protein | AT1G14320 | 0,008298 | ATHE_i vs Lti6B_i | 24,8725 | 24,5984 | 23,6086 | 25,1565 | 25,9403 | 24,3695 |
| S18 ribosomal protein | AT1G22780; AT1G34030; AT4G09800 | 0,000504 | ATHE_i vs Lti6B_i | 26,7774 | 27,5393 | 27,3347 | 27,5667 | 27,1137 | 27,3753 |
| Ribosomal protein L22p/L17e family protein | AT1G27400 | 4,83E-12 | ATHE_i vs Lti6B_i | 24,3077 | 25,4234 | 24,6435 | 24,6331 | 25,4416 | 24,0699 |
| Jacalin lectin family protein | AT1G33790 | 8,70E-10 | ATHE_i vs Lti6B_i | 27,245 | 27,8203 | 28,1788 | 27,1618 | 23,5967 | 25,508 |
| choice-of-anchor C domain protein 2C putative (Protein of unknown function 2C DUF642) | AT1G80240 | 0,00628 | ATHE_i vs Lti6B_i | 0 | 21,8731 | 22,2392 | 21,5674 | 23,0374 | 21,3167 |
| Ribosomal protein L18ae/LX family protein | AT2G34480 | 0,008855 | ATHE_i vs Lti6B_i | 23,4998 | 24,1471 | 23,1017 | 24,852 | 24,5922 | 24,1818 |
| Ribosomal protein S5 family protein | AT2G41840 | 0,006585 | ATHE_i vs Lti6B_i | 23,6181 | 24,2698 | 24,1575 | 23,8807 | 24,1066 | 24,689 |
| Ribosomal protein S11 family protein | AT2G36160; AT3G11510 | 0,01195 | ATHE_i vs Lti6B_i | 26,1268 | 27,4118 | 26,9652 | 26,4577 | 26,1737 | 26,9131 |
| Ribosomal protein L13 family protein | AT3G24830 | 0,008753 | ATHE_i vs Lti6B_i | 24,9437 | 24,7667 | 24,5083 | 25,605 | 25,792 | 24,6563 |
| ATHE, Leucine-rich repeat protein kinase family protein | AT3G46330 | 4,32E-12 | ATHE_i vs Lti6B_i; ATHE_m vs Lti6B_m | 30,7313 | 31,2323 | 31,5607 | 31,5828 | 32,2116 | 29,8411 |
| MIK2, Leucine-rich repeat receptor-like protein kinase family protein | AT4G08850 | 0,00409 | ATHE_i vs Lti6B_i; ATHE_i vs ATHE_m | 0 | 20,6896 | 0 | 21,8766 | 22,9121 | 27,6643 |
| Ribosomal protein L23/L15e family protein | AT4G16720; AT4G17390 | 0,008139 | ATHE_i vs Lti6B_i | 23,0431 | 22,9245 | 22,6448 | 23,5655 | 23,9098 | 23,3741 |
| Ribosomal protein S3Ae | AT4G34670 | 2,26E-11 | ATHE_i vs Lti6B_i | 24,375 | 25,7278 | 23,9886 | 25,3535 | 24,4643 | 24,853 |
| Heavy metal transport/detoxification superfamily protein | AT4G35060 | 0,005643 | ATHE_i vs Lti6B_i | 0 | 0 | 21,272 | 21,7031 | 21,9929 | 26,225 |
| Raffinose synthase family protein | AT5G20250 | 2,79E-07 | ATHE_i vs Lti6B_i; ATHE_i vs ATHE_m | 0 | 0 | 0 | 21,6381 | 22,9084 | 28,2754 |

| sity (a.u.) | | | | | |
|---|---|---|---|---|---|
| Lti6B_m | | | Lti6B_i | | |
| Rep1 | Rep2 | Rep3 | Rep1 | Rep2 | Rep3 |
| 24,2555 | 24,8417 | 0 | 0 | 0 | 0 |
| 27,8165 | 27,934 | 26,6726 | 25,3926 | 24,4964 | 24,9804 |
| 24,4246 | 24,6045 | 24,5659 | 0 | 0 | 0 |
| 27,5948 | 27,6808 | 26,9666 | 0 | 0 | 0 |
| 0 | 0 | 0 | 0 | 0 | 0 |
| 23,9352 | 24,4101 | 0 | 0 | 0 | 0 |
| 24,4016 | 0 | 0 | 0 | 0 | 0 |
| 26,4901 | 26,7021 | 26,569 | 25,6912 | 25,3671 | 25,3519 |
| 24,9961 | 25,2114 | 0 | 0 | 0 | 0 |
| 0 | 0 | 0 | 0 | 0 | 0 |
| 0 | 0 | 0 | 0 | 0 | 0 |
| 23,1694 | 22,8857 | 0 | 0 | 0 | 0 |
| 24,8753 | 24,8036 | 24,1327 | 0 | 0 | 0 |
| 0 | 0 | 0 | 0 | 0 | 0 |
| 0 | 0 | 0 | 0 | 0 | 0 |

**Table S3.** List of primers used in this work

| Name | Sequence (5’ □ 3’) | Use |
|---|---|---|
| SALK_108641 LP | CAAACTTTCTAGATGCGCCAG | Genotyping *athe-1* |
| SALK_108641 RP | GGCGAATGATACTTACATCGC | |
| SALK_LBb1 | CGCTTTCTTCCCTTCCTTTCTC | |
| *mik2-1* LP | AACGGATCGATTCCTTCTGA | Genotyping *mik2-1* |
| *mik2-1* RP | TTTTGCCTGATAGCCGATTC | |
| pNew_EcoRI_pATHE_Fw | CGAATTGGAGCTGCGGCCGCGAATTCATCCTCTTACTTTCCAAGCAAAATCAAGA | Cloning *pAHTE:ATHE-GFP* and *pATHE:mCherry-N7* |
| pNew_pATHE_Rev | GATTCCTCCGATCACAATACACGC | Cloning *pATHE:ATHE-GFP* |
| gATHE_Fw | GTATTGTGATCGGAGGAATCATGAAGAATCTTTGTTGGGTTTTTCTGTC | |
| gATHE_Ala_Rev | CGCCGCTGCTGCGGCGCCTCTTGCCTTAGGCTTCACATCAGTATC | |
| Ala_GFP_Fw | GGCGCCGCAGCAGCGGCGATGGTGAGCAAGGGCGAGG | |
| GFP_XbaI_Rev | TTTCATCTTCATCTTCATATTCTAGATTACTTGTACAGCTCGTCCATGCCG | |
| pATHE_mCh_Rev | CCTCGCCCTTGCTCACCATGATTCCTCCGATCACAATACACGC | Cloning *pATHE:mCherry-N7* |
| mCh_Fw | ATGGTGAGCAAGGGCGAGG | |
| mCh_Ala_Rev | GGCGCCCGCCGCCGCCGCCGCTCCCTTGTACAGCTCGTCCATGCC | |
| Ala_Fw | GGAGCGGCGGCGG | |
| N7_XbaI_Rev | TTTCATCTTCATCTTCATATTCTAGATCACTCTTCTTCTTGATCAGCTTCTGTGTCG | |
| ATHE_RT_Fw | ATGAAGAATCTTTGTTGGGTTTTTCTGTC | Examining the knock-out mutation in *athe-1* *(ACTIN1, At2g37620, was used as a control)* |
| ATHE_RT_Rev | CCACGGTTCAGGTTTATTTCTTG | |
| ACTIN1_RT_Fw | TGGTTGGGATGGGGCAAAAG | |
| ACTIN1_RT_Rev | ATTTCACGCTCTGCTGTGGTGG | |
| WRKY45_Fw | GAACAATCCATTCCCCAGGAG | qRT-PCR (de Azevedo Souza *et al* ., 2017) |
| WRKY45_Rev | GGAGGGAAGATGTGCATTTGTG | |
| WRKY53_Fw | GCGACAAGACACCAGAGTCA | qRT-PCR (Masachis *et al* ., 2016) |
| WRKY53_Rev | ACCGTTGGATTGAACCAGTC | |

| | | |
|---|---|---|
| AT1G51890_Fw | CTAGCCGACTTTGGGCTATC | qRT-PCR (Van der Does *et al*., 2017) |
| AT1G51890_Rev | CCAGTTTGTTCTGTAATACTCAGG | |
| AT1G05340_Fw | TCGGTAGCTCAGGGTAAAGTGG | qRT-PCR (Deckers *et al*., 2020) |
| AT1G05340_Rev | CCAGGGCACAACAGCAACA | |
| EXLA2_Fw | AGGAGGCCAAACTGAAGTGG | qRT-PCR (Tsugama *et al*., 2016) |
| EXLA2_Rev | CATTTTGCCGTCGTAGCCTG | |
| XTH23_Fw | TGGTTCGTGGTTGTCTCAGG | qRT-PCR (Tsugama *et al*., 2016) |
| XTH23_Rev | CTAAGCACTCGCGTGGAAGA | |
| WRKY30_Fw | CGGAGCCAAATTTCCAAGAGG | qRT-PCR (de Azevedo Souza *et al*., 2017) |
| WRKY30_Rev | GACGGAGAGTTTGATGCTGAG | |
| WRKY40_Fw | AGCCCTCCCAAGAAACGCAATC | |
| WRKY40_Rev | GCTTGGAGCACAAGCACATTTGAAG | |
| ATHE_Fw | CATCGAGTGGCTTGAGTGGG | qRT-PCR (This work) |
| ATHE_Rev | TAGTGGCTAGAAACTCGGGC | |
| GAPDH 600b_Fw | AGGTGGAAGAGCTGCTTCCTTC | qRT-PCR (Czechowski *et al*., 2005) |
| GAPDH 600b_Rev | GCAACACTTTCCCAACAGCCT | |
| ATHE_F-XhoI | GCCTCGAGATGAAGAATCTTTGTTGGGT | Cloning p35S::ATHE-HA |
| ATHE_R-HindIII | CTAAGCTTCTAAGCGTAATCTGGAACATCGTATGGGTATCTTGCCTTAGGCTTCAC[illegible] | |

**Table S4.** Sequences of RALF peptides used in this work

| RALF peptide | amino acid sequence |
|---|---|
| RALF1 | ATTKYISYQSLKRNSVPCSRRGASYYNCQNGAQANPYSRGCSKIARCRS |
| RALF23 | ATRRYISYGALRRNTIPCSRRGASYYNCRRGAQANPYSRGCSAITRCRRS |
| RALF34 | YWRRTKYYISYGALSANRVPCPPRSGRSYYTHNCFRARGPVHPYSRGCSSITRCRR |
| RALF Fo5176 (Fo-RALF) | PAAKPQGEISYGALNRDHIPCSVKGASAANCRPGAEANPYNRGCNAIEKCRGGVGDN |
